# A Temporal Single-Cell Multi-Omics Atlas of Murine Pancreatic Islet Remodeling During Hyperglycaemia Progression

**DOI:** 10.1101/2025.05.29.656754

**Authors:** Simran Singh, Musale Krushna Pavan, Luiz F. Barella, Jayesh Telang, Ajita Shree, Shruti Agarwal, Ayush Goel, Saahiba Thaleshwari, Jürgen Wess, Hamim Zafar, Sai Prasad Pydi

**Affiliations:** Molecular Metabolism & Cell signaling lab, Department of Biological Sciences & Bioengineering, Indian Institute of Technology Kanpur, UP 208016, India; CoSmic Lab, Department of Biological Sciences & Bioengineering, Indian Institute of Technology Kanpur, UP 208016, India; Department of Computer Science & Engineering, Indian Institute of Technology Kanpur, UP 208016, India; Molecular Signalling Section, Laboratory of Bioorganic Chemistry, National Institute of Diabetes and Digestive and Kidney Diseases, Bethesda, MD 20892, USA; Mehta Family Centre for Engineering and Medicine, Indian Institute of Technology Kanpur, UP 208016, India; Gangwal School of Medical Sciences and Technology, Indian Institute of Technology Kanpur, UP 208016, India

**Keywords:** Single-cell RNA-sequencing, Single-cell ATAC-sequencing, pancreatic islets, cell-cell communication, insulin resistance, type 2 diabetes

## Abstract

Pancreatic islets undergo coordinated cellular remodeling during obesity-induced insulin resistance (IR). However, the associated molecular changes across endocrine and non-endocrine compartments remain largely unexplored. Here, using longitudinal single-cell RNA sequencing (scRNA-seq) and single-cell ATAC sequencing (scATAC-seq) on islets from C57BL/6 mice subjected to high-fat diet (HFD) feeding for 8, 16, and 24 weeks, along with age-matched controls on regular chow, we mapped dynamic changes in islet cell composition and transcriptional states. Beta cells demonstrated pronounced stress-induced reprogramming, with the emergence of proliferative and dysfunctional subsets. Alpha and delta cell fractions declined under HFD, despite increased polyhormonal biosynthesis, suggesting functional rather than numerical adaptation. Immune profiling showed robust expansion of proinflammatory M1 macrophages and upregulation of NF-κB and chemotaxis pathways, particularly at 16 weeks. Notably, cell-cell communication analyses revealed diet-specific disruption in signaling networks. Under HFD conditions, intercellular communication among beta cells, macrophages, and delta cells was markedly altered, leading to the disruption of key signaling pathways such as the gastric inhibitory polypeptide receptor (GIPR) and major histocompatibility complex-I (MHC-I). Notably, C-C motif chemokine ligand 27A (*Ccl27a*) expression and chromatin accessibility were significantly altered in a distinct subpopulation of beta cells under HFD condition, indicative of a niche-specific regulatory mechanism. Integration with human islet datasets from obese and type 2 diabetes (T2D) donors confirmed conserved shifts in beta cell identity and immune activation. This study presents a comprehensive high-resolution atlas of islet remodeling under metabolic stress, identifying key communication nodes and transcriptional programs pertinent to T2D pathogenesis.

**Graphical Abstract:** 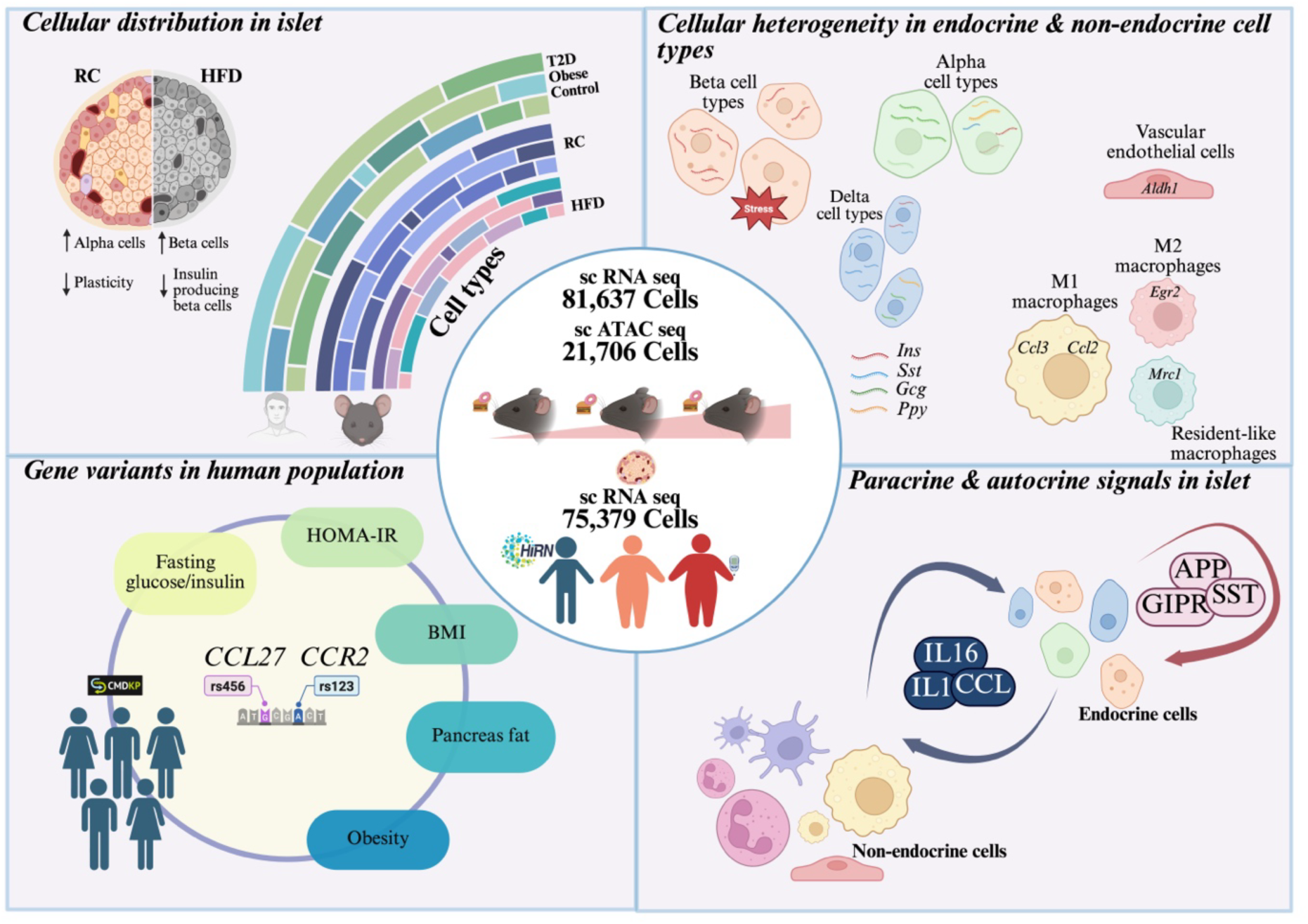

## Introduction

Pancreatic islets are dynamic microorgans composed of various endocrine cell types, including glucagon-secreting alpha cells, insulin-producing beta cells, and somatostatin-releasing delta cells (1-4). These endocrine cells work in tandem to maintain blood glucose and energy homeostasis (5, 6). A sophisticated intercellular communication network is vital for the coordinated function of islet cells under various metabolic states. The inherent cellular heterogeneity of pancreatic islets and their unique functional states contribute to the ability of the endocrine pancreas to adjust to changing metabolic demands (7-9). Factors including age, metabolic status, and environmental stressors add extra layers of complexity to islet biology (10, 11). Non-endocrine cells, such as immune, endothelial, and stellate cells, modulate islet activity, creating a complex microenvironment vulnerable to metabolic stressors like obesity and insulin resistance (IR) (12-14). Obesity is the primary driver of IR and a critical hallmark of type 2 diabetes (T2D) pathogenesis (10). As T2D progresses from IR to overt diabetes, significant disruptions occur in islet cell architecture and dynamics, particularly impacting immune and hormone-secreting cells (15-18). Although extensive studies have explored beta cell function, the contributions of other endocrine and non-endocrine cells in the islet microenvironment during IR have received relatively less attention. Recent studies, however, emphasize the importance of these cells under nutritional stress. For instance, alpha cells can elevate glucagon secretion or even transdifferentiate into insulin-producing cells under certain conditions (19). Similarly, delta cells, responsible for the production of somatostatin, play a pivotal role in the paracrine regulation of beta and alpha cell function; however, their contribution to islet dysfunction during the progression of T2D remains poorly understood (20). Hence, there is an urgent need for a comprehensive, cell type-specific understanding of islet biology during metabolic stress.

Recent advances in single-cell RNA sequencing (scRNA-seq) have improved our understanding of pancreatic islet biology by allowing precise characterization of cellular heterogeneity and functional diversity within islets. While numerous studies have employed a variety of model systems and experimental manipulations, including chemical treatments, genetic modifications, and dietary interventions, to investigate molecular changes in islet cells, the primary focus has often been on beta cells (21-29). These targeted studies have provided valuable insights but did not explore the intricate intercellular communication and molecular pathways regulating islet function in health and disease. Moreover, few studies employed integrative approaches that combine multiple datasets to capture the diverse cellular states within islets (24). The challenge associated with scRNA-seq studies lies in harmonizing data across varying experimental conditions, genetic backgrounds, and sequencing methodologies. This complexity complicates direct comparisons and necessitates high operational standards.

To address these knowledge gaps, we employed a longitudinal, multi-omics approach to dissect the spatiotemporal dynamics of mouse pancreatic islets during metabolic stress. Our findings revealed dynamic reprogramming of both endocrine and non-endocrine cell populations. Key findings include dynamic shifts in cell composition and transcriptional reprogramming within beta cell subclusters that underpin the progression to T2D, as well as adaptive plasticity in alpha, delta, and gamma cells characterized by lineage-mixed states. In addition to endocrine cells, we observed significant changes in endothelial and immune cell populations, emphasizing the critical role of the islet microenvironment in T2D pathogenesis. Furthermore, cell-cell communication analysis revealed disrupted chemokine signaling networks in endocrine-immune coordination. Together, these results outline a hierarchical map of islet cell-state transitions during metabolic adaptation, offering novel mechanistic insights into early T2D pathogenesis. Clearly, these data hold considerable translational relevance.

## Results

### Cellular heterogeneity in pancreatic islets under euglycemic and hyperglycemic conditions

To investigate molecular changes in pancreatic islet cells during the progression towards IR, we performed scRNA-seq on islets from C57/BL6 mice (*Mus musculus-*Mm) fed with a high fat diet (HFD) for 8, 16, and 24 weeks starting at 6 weeks postnatal, along with age-matched maintained on regular chow (RC) for 8, 14, 22, and 30 weeks, as well as 8-week-old RC mice, thus covering a range of prediabetic conditions (**Figure 1A**). Using the 10x Genomics Chromium platform, we generated 21 scRNA datasets, with each condition including three replicates and 100 islets per mouse for analysis. As expected, HFD feeding led to increased body weight, impaired glucose and insulin tolerance, and elevated levels of fed and fasted state blood glucose and plasma insulin, glycerol, and triglycerides over time (**Figure 1B, Figure S1A-H**).

**Figure 1.**
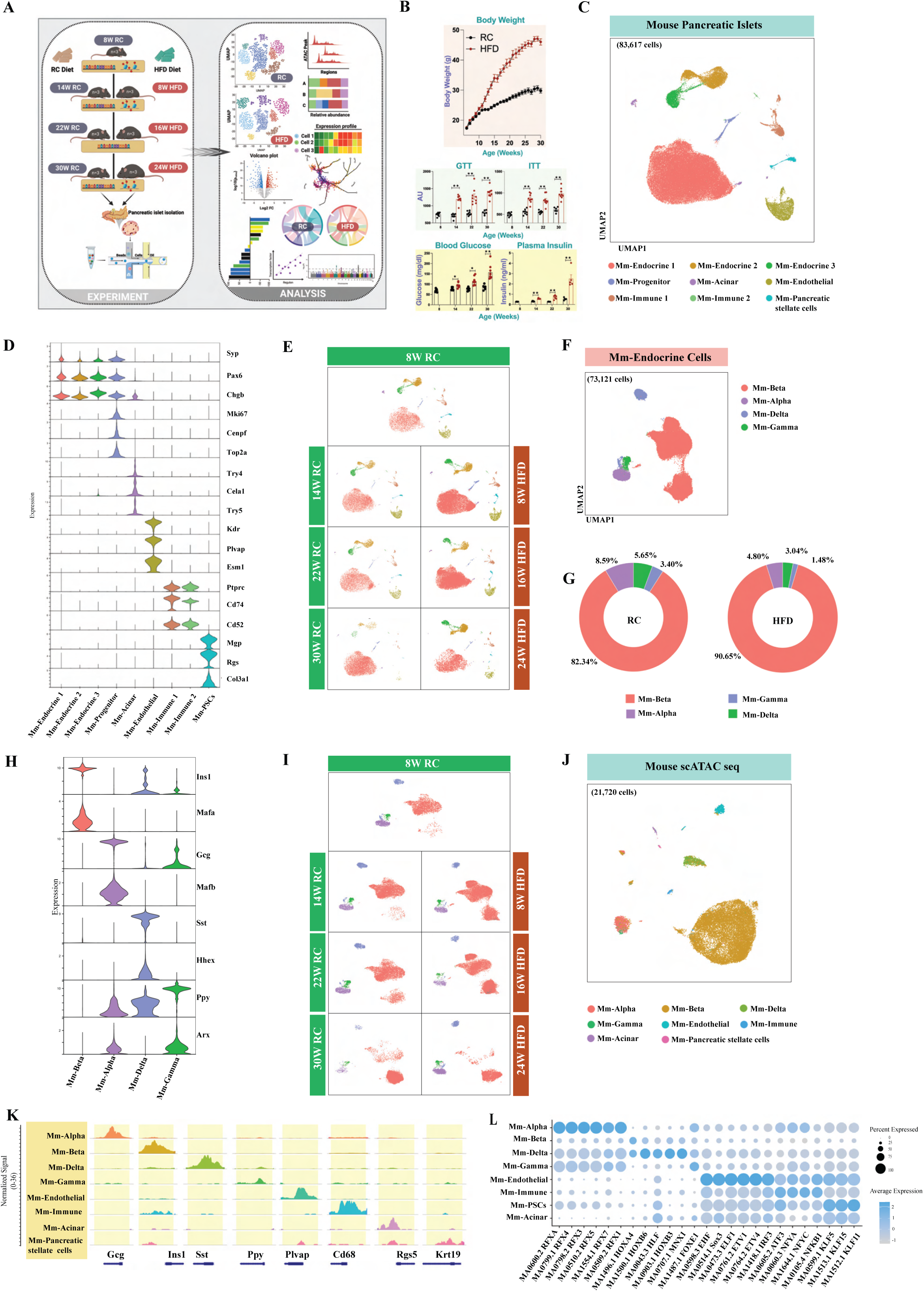
Transcriptomic and chromatin accessibility atlas of mouse pancreatic islets across a spectrum of hyperglycemic and hyperinsulinemic conditions. (A) Overview of the experimental workflow: Single-cell RNA sequencing (scRNA-seq) was performed on pancreatic islets from mice at 8, 14, 22, and 30 weeks on a regular chow diet (RC), as well as at 8, 16, and 24 weeks on a high-fat diet (HFD) (n=3 for each condition). (B) Body weight (upper panel), glucose tolerance test, insulin tolerance test (middle panel) and fasting glucose, plasma insulin (lower panel) in RC and HFD mice. (C) Uniform Manifold Approximation (UMAP) visualization of n=83617 cells from all time points across dietary conditions, derived from scRNA-seq data of pancreatic islets. Cells are colored based on canonical cell types identified through the expression of specific marker genes. (D) Expression profile of marker genes used for annotating different islet cell types. (E) UMAP visualization of islet cell types at 8, 14, 22, and 30 weeks on RC, as well as at 8, 16, and 24 weeks on HFD. (F) UMAP visualization of endocrine cells (n=73121) from all timepoints, obtained from scRNA-seq analysis of islet cells. Cells are colored based on canonical cell types determined by the expression of marker genes or differentially expressed genes. (G) Distribution of endocrine cell types across RC and HFD conditions. (H) Expression profile of marker genes used for annotating different endocrine cell type clusters. (I) UMAP visualizations of endocrine cells at different timepoints, including 8, 14, 22, and 30 weeks of RC, as well as 8, 16, and 24 weeks of HFD. (J) UMAP visualization of *n* = 21720 cells from all time points, obtained from single-cell Assay for Transposase Accessible Chromatin with sequencing (scATAC-seq) of pancreatic islets. (K) Cell-type-specific chromatin accessibility profiles at canonical cell type markers identified in scRNA-seq of pancreatic islets. The height of links represents their significance. (L) Enrichment of motifs for different cell type clusters identified through scATAC-seq of pancreatic islets.

After quality control filtering and doublet removal, 83,617 cells were retained across conditions (36,518 for RC and 47,099 HFD, respectively) in the scRNA-seq dataset (**Figure 1C**). These cells were integrated to remove batch effects, revealing nine major cell subpopulations, including three endocrine, two immune, and one each for endothelial, acinar, progenitor, and pancreatic stellate cells (PSCs) (**Figure 1C & D; Table 1**). Mm-Endocrine cells made up the majority of islet cells, comprising 84.5% (RC) and ∼90% (HFD) of cells, respectively, with notable age-related changes across conditions (**Figure 1E**). Mm-endothelial and Mm-immune cell populations declined with HFD exposure compared to RC, showing distinct age-dependent patterns (**Figure 1E, Figure S2A-C**). Mm-PSCs, Mm-progenitor cells, and Mm-acinar cells exhibited similar trends across both diets. Overall, we successfully identified and characterized the major cell types in the pancreatic islets, enabling a comprehensive analysis of their heterogeneity during the progression of IR.

To explore endocrine population dynamics under different diets, we sub-clustered 73,121 murine endocrine cells (30,706 (RC) and 42,451 (HFD), respectively), identifying seven distinct subpopulations (**Figure 1F-G, Figure S2D**). Differentially expressed gene (DEG) analysis confirmed Mm-Beta (*Ins1*), Mm-Alpha (*Gcg*), Mm-Delta (*Sst*), and Mm-Gamma (*Ppy*) cell populations (**Figure 1H**). The Mm-Alpha, Mm-Gamma, and Mm-Delta populations were lower under HFD compared to the RC feeding (**Figure 1G**, **Table 1**). Mm-Beta cells represented 90% of the endocrine cells under RC and 82% under HFD conditions, respectively. Mm-Beta cells changed dynamically with age in both RC and HFD conditions (**Figure 1I; Figure S2E**). Mm-

Alpha cells were the second-largest cell population, decreasing in RC mice with age but increasing in HFD mice, and a similar trend was observed with Mm-Delta cells (**Figure 1I; Figure S2E**).

To correlate transcriptomic and epigenetic changes and validate the findings from scRNA-seq data, we performed a single-cell sequencing assay for transposase-accessible chromatin sequencing (scATAC-seq) to generate a chromatin accessibility profile for RC islets (8, 14, and 30 weeks of RC feeding), and after HFD islets (8 and 24 weeks of HFD consumption) (**Figure 1A**). After quality control, we generated scATAC-seq data for 25,932 nuclei with 567,365 accessible chromatin sites (**Figure 1J**). Using scRNA-seq dataset as a reference, we identified eight subpopulations in our scATAC-seq data, assigned cell types, and annotated them with canonical markers (**Figure 1J, Figure S2F**). Differential chromatin accessibility was noted in Mm-Beta (*Ins1*), Mm-Alpha (*Gcg*), Mm-Delta (*Sst*), Mm-Gamma (*Ppy*), Mm-Endothelial (*Plvap*), Mm-Immune (*Cd68*), Mm-PSCs (*Rgs5*), and Mm-Acinar cells (*Krt19*) (**Figure 1K, Table-2**). Motif enrichment via ChromVAR highlighted cell type-specific motifs, including *RFXs* in Mm-Alpha cells, *HOXA4* in Mm-Beta and Mm-Delta cells, *HOXB3/6* and *MNK1* in Mm-Delta cells, *SOX3/6* in Mm-Endothelial cells, *ATF3* and (**Figure 1L**).

To compare human and mouse pancreatic islets at single-cell resolution across healthy, obese, and T2D conditions, we analysed human (*Homo sapiens-* Hs) pancreas data from PancDB (**Figure S3A**) (30). We categorized the PancDB data based on body weight, age, disease condition, and HbA1c levels **(Table 3)**. After applying quality control, we retained 10,868, 41,181, and 21,699 cells for healthy, obese, and T2D groups, respectively, totalling 80,211 cells for further analysis. After clustering, we identified nine cell types, including Hs-Endocrine, Hs-Acinar, Hs-PSCs, Hs-Endothelial, Hs-Immune, and Hs-Ductal cells (**Figure S3B, Figure S3C**). Hs-Endocrine cells were most abundant, followed by Hs-Acinar cells (**Figure S3D**). Further analysis of Hs-Endocrine cells revealed three main clusters (Hs-Alpha, Hs-Beta, Hs-Gamma/Delta) (**Figure S3E-H**), and Hs-Alpha cells were predominant, unlike mice, where Mm-Beta cells were most abundant (**Figure S3G-I**).

### Stage-specific transcriptional and epigenetic changes in beta cells upon HFD challenge

To investigate the transcriptional and epigenetic alterations in Mm-Beta cell responses during HFD feeding, we further analyzed the Mm-Beta cell population (63,399 cells; 25,062 RC; 38,337 HFD) and identified five subpopulations (Beta-1 through 5), including two HFD-specific populations (clusters 4 and 5 consisting of 154 and 17 cells, respectively) (**Figure 2A-B**). Mm-Beta 1 cells, the dominant insulin-secreting population, comprised 62.5% of beta cells in RC mice but decreased to 40.8% in HFD mice. In contrast, Mm-Beta 2 and Mm-Beta 3 populations expanded under HFD, representing 43.5% and 15.3% of the population, respectively, compared to 33.0% and 4.5% in RC mice (**Figure 2C, Figure S4A, Table 1**). Age-related adaptations were also evident, with Mm-Beta 1 comprising >90% of beta cells at 8 and 14 weeks and ∼80% at 22 weeks in RC mice. In HFD mice, Mm-Beta 1 decreased to 60% and 30% at 8 and 16 weeks, respectively, while Mm-Beta 2 dominated (∼95%) by 30 weeks of RC feeding and 24 weeks of HFD consumption. Mm-Beta 3 showed minimal temporal changes in RC mice but was enriched at 8 and 16 weeks under HFD conditions. However, it nearly disappeared by 24 weeks of HFD (**Figure 2D, Figure S4B**). DE and GSEA analyses revealed functional distinctions between beta cell subpopulations. Mm-Beta 1 exhibited enrichment of pathways such as intracellular protein transport, glucose-stimulated insulin secretion, and exocytosis and high expression of insulin secretion-related genes (31) (*Pdx1, Mafa, Neurod1, Nkx6.1*, and *FoxO1*) confirming its role as a classical insulin-secreting beta cell (**Figure 2E-F, Figure S4C**). Mm-Beta 2 displayed upregulation of ribosomal protein genes, with GSEA showing enrichment in ribosomal biosynthesis, translation, mitochondrial processes, and general metabolism, while pathways related to insulin signaling and vesicle transport were downregulated (**Figure 2B & E**). Mm-Beta 3 exhibited increased expression of stress-related genes (*Fosb, Nktr, Jun*), and GSEA indicated upregulation of hypoxia, vesicle-mediated transport, inflammatory response, and NF-κB-mediated TNFα signaling, alongside downregulation of mitochondrial respiration and TCA cycle genes (**Figure 2E**). As Mm-Beta 1 is a classical population, to further characterize Mm-Beta 2 and Mm-Beta 3, we performed a pairwise analysis, which revealed Mm-Beta 2 as a polyhormonal population expressing *Ppy* and *Sst*, while Mm-Beta 3 was identified as a proinflammatory cluster with elevated expression of NF-κB mediated TNFα pathway genes (**Figure S4D; Table 4**).

**Figure 2.**
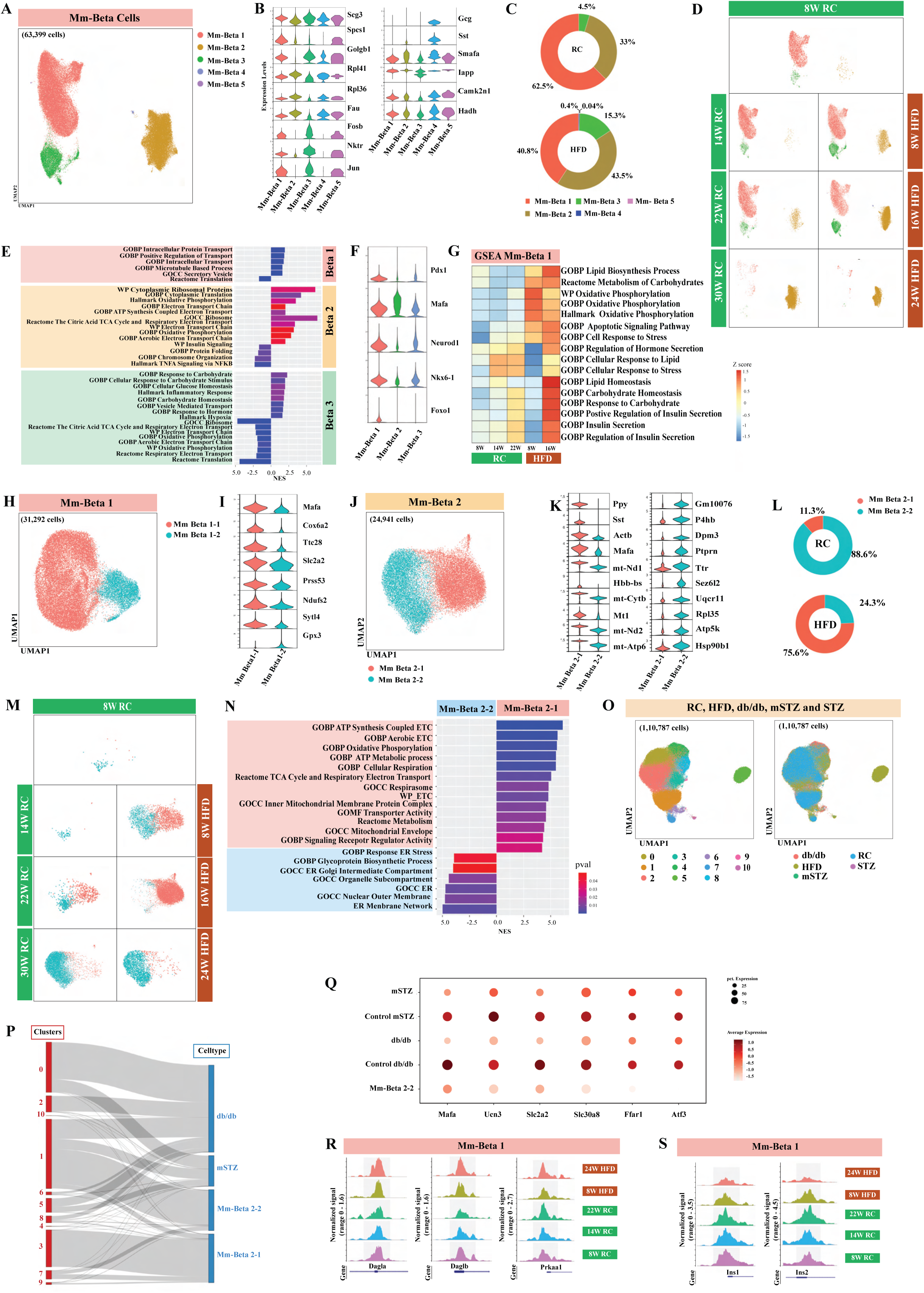
Temporal transcriptional and epigenomic adaptations in mouse beta cells during HFD exposure. (A) UMAP visualization of n = 63,399 beta cells clustered into 5 populations. Cells are colored by the beta population (Mm-Beta 1, Mm-Beta 2, Mm-Beta 3, Mm-Beta 4 and Mm-Beta 5). (B) Expression profile of differentially expressed genes in various beta cell subpopulations. (C) Distribution of beta cell subpopulations across RC and HFD conditions. (D) UMAP visualization of beta cell subpopulations across different time points across diet. (E) Pathways enriched in beta cell subpopulations inferred using GSEA. (F) Expression profile of key genes (*Pdx1*, *Mafa*, *Neurod1*, *Nkx6-1*, and *FoxO1*) in different beta cell subpopulations. (G) Heatmap of Gene Ontology (GO) and KEGG pathways enriched in Mm-Beta 1 subpopulation across different time points across diet conditions. (H) UMAP visualization of Mm-Beta 1 (*n* = 31292) subpopulations. (I) Expression profile of differentially expressed genes in various Mm-Beta 1 subpopulations. (J) UMAP visualization of n = 24941 Mm-Beta 2 cells clustered into two subpopulations. (K) Expression profile of differentially expressed genes across Mm-Beta 2 cell subpopulations. (L) Distribution of Mm-Beta 2 subpopulations under RC and HFD conditions. (M) UMAP visualization of Mm-Beta 2 subpopulations across different dietary conditions (RC, HFD) and timepoints. (N) Hallmark pathways enriched in Mm-Beta 2 subpopulations. (O) UMAP visualization of cellular embeddings after the integration of beta cells from RC and HFD mice with the beta cell populations from diabetes mice models from Feng et al., Sachs et al., and Hrovatin et al., Cells are colored by unsupervised clusters inferred from the integrated embeddings and conditions. (P) Sankey plot representing the correspondence between clusters inferred from integrated embeddings and beta cell subpopulations (Mm-Beta 2-1, Mm-Beta 2-2, db/db and mSTZ). (Q) Expression of beta-cell function genes across different diabetes mice models, their corresponding healthy controls and Mm-Beta 2-2 subpopulation in our dataset. (R) Chromatin accessibility profile of Mm–Beta 1 population across RC and HFD conditions across timepoints for genes involved in lipid and carbohydrate metabolism (*Dagla, Daglb, Prkaa1)*. (S) Chromatin accessibility profile of Mm–Beta 1 population across RC and HFD conditions across timepoints for *Ins1* and *Ins2*.

To investigate beta cell adaptations to different dietary conditions, we performed DE and pathway analysis of the Mm-Beta 1 population in obesity progression with age-matched healthy/lean conditions. In the HFD group, pathways related to lipid biosynthesis, carbohydrate metabolism, oxidative phosphorylation, cellular stress, and apoptotic signaling were upregulated at 8 and 16 weeks (**Figure 2G**). Conversely, pathways associated with lipid and carbohydrate homeostasis and insulin secretion increased with age under RC but were disrupted after 8 weeks of HFD feeding before significantly increasing at 16 weeks. This observation suggests a potential transition to IR and compensatory mechanisms by cells in response to elevated glucose levels (**Figure 2G**). Further, subclustering of Mm-Beta 1 cells revealed two distinct groups: Mm-Beta 1-1 and Mm-Beta 1-2 (**Figure 2H, Figure S4E and F**). Mm-Beta 1-1 cells exhibited increased expression of genes linked to insulin secretion, including *Mafa, Slc2a2, Sytl4*, and *Ttc28* (**Figure 2I**). Enhanced expression of *Sytl4*, known to regulate dense-core granule exocytosis, suggests enhanced secretory function (32, 33). In contrast, Mm-Beta 1-2 cells expressed higher levels of *GPX3*, an antioxidant enzyme that protects against oxidative damage (**Figure 2I**). The Mm-Beta 2 population expanded significantly with the duration of HFD feeding and aging (**Figure S4E**), and we further subclustered it into Mm-Beta 2-1 and Mm-Beta 2-2 populations (**Figure 2J**). Mm-Beta 2-1 had higher expression of *Ppy, Sst, Actb, Mafa,* and *Mt1*, while Mm-Beta 2-2 expressed high levels of *P4hb, Ptprn, Ttr,* and *Sez6l2* (**Figure 2K**). Under RC conditions, Mm-Beta 2-1 generally comprised <12% of the entire Mm-Beta 2 population, except for 22 weeks, during which it expanded to represent > 50% of the Mm-Beta 2 population. In contrast, under HFD conditions, Mm-Beta 2-1 and Mm-Beta 2-2 were equally distributed at 8 weeks, but by 16 weeks of HFD feeding, Mm-Beta 2-1 dominated (97%), followed by a sharp transition to Mm-Beta 2-2 predominance (92%) at 24 weeks (**Figure 2L-M, Figures S4G-H**). Pathway analysis of Mm-Beta 2-1 showed the upregulation of pathways related to oxidative phosphorylation, TCA cycle, and electron transport, indicative of increased metabolic demand at 8 weeks of HFD feeding (**Figure S4I**). In Mm-Beta 2-1, the elevated expression of *Mt1*, a negative regulator of insulin secretion (34), indicates beta cell dysfunction, and high expression of *Sst, Ppy,* and *Mafa* is suggestive of probable transdifferentiation processes **(Figure 2K)**. In contrast, Mm-Beta 2-2 was enriched in ER stress and Golgi-intermediate pathways, consistent with impaired function under sustained metabolic stress (**Figure 2N, S4J**). Analysis of the third beta subpopulation (Mm-Beta 3) resulted in three subpopulations: Mm-Beta 3-1, Mm-Beta 3-2, and Mm-Beta 3-3 (**Figure S5A**). Mm-Beta 3-1, characterized by upregulation of glucose response genes (*Nktr, Mlxipl, Neat1*) (**Figure S5B**), was absent under RC conditions but was predominant (90%) in HFD mice at 16 weeks, suggesting adaptation to increased blood glucose levels (**Figures S4C-D**). *Mlxipl* (ChREBP) and its downstream target *TXNIP* are implicated in glucose-induced inflammation and apoptosis, aligning with beta-cell lipotoxicity in obesity (35). The Mm-Beta 3-2 subpopulation, expressing *Rps10, Eef1a1, Tpt1,* was associated with ribosome biogenesis and cell cycle processes. Mm-Beta 3-3, enriched in transcriptional regulators (e.g., *Atf3, Etv1,* and *Klf4)*, was absent in RC mice but emerged under HFD conditions (**Figure S4B&E**).

To further characterize our beta cell subpopulations and compare them against known beta cell states enriched in diabetes mouse models (21), we integrated the beta cells in RC and HFD mice with beta cells from T2D (db/db) mouse model (36), streptozotocin (STZ)-induced (mSTZ, multiple low doses) beta-cell ablation model (37), and a single high dose STZ-induced diabetes model (23) (**Figure 2P, Figure S6A**). All the 110,187 beta cells were mapped to a common latent space using scDREAMER (38), and their joint analysis revealed nine beta cell states. Interestingly, Mm-Beta 2-2 subpopulation mapped to cluster 1 along with beta cells from db/db and mSTZ models, indicating their transcriptomic similarity (**Figure 2P, Figure S6B**). We further observed that genes which are known to be downregulated in the diabetic mouse models as compared to their control counterpart (e.g., genes involved in beta cell maturation (*Mafa, Ucn3, Slc2a2*), insulin secretion (*Slc30a8*), insulin synthesis and processing (*Ffar1*) and unfolded protein response (*Atf3*)) were also downregulated in Mm-Beta 2-2 (**Figure 2Q**) indicating their shared phenotype and the involvement of Mm-Beta 2-2 in metabolic stress. Moreover, Mm-Beta 2-2 marker genes (*P4hb, Dpm3, Ptprn, Ttr, Sez6l2, Rpl35, Atp5k* and *Hsp90b1*) had high expression in db/db and mSTZ beta cells (**Figure S6C)**. In contrast, Mm-Beta 2-1 did not cluster with any established beta cell state; rather, it mapped to a completely separate cluster (**Figure 2P),** highlighting that its polyhormonal signature is distinct from known diabetic murine beta cell states.

To corroborate the scRNA-seq findings, we analyzed the scATAC-seq profiles of pancreatic beta cells. A total of 18052 nuclei were analyzed across 5 conditions, including 3 RC (7813 nuclei) and 2 HFD (10239 nuclei) conditions. In sc-ATAC, we primarily found Mm-Beta 1 cells, while other beta cell subpopulations were present at low abundance (**Figure S6D**). In line with scRNA-seq data, genes involved in lipid and carbohydrate metabolism (*Dagla*, *Daglb* and *Prkaa*) were highly accessible under HFD conditions (**Figure 2G and 2R**). However, the chromatin accessibility for *Ins1* and *Ins2* genes was reduced in HFD mice (**Figure 2G, 2S and Figure 2SE**).

Trajectory analysis using MARGARET (39) traced beta-cell differentiation, where the progenitor cells gave rise to Mm-Beta 1 cells; which further branched into two lineages giving rise to Mm-Beta 2 and Mm-Beta 3 (**Figures S7A-D**). To further characterize the beta cell lineages, we investigated the expression trends of transcription factors along the pseudotime. Interestingly, *Atf5* and *Tshz1* expression increased in the Mm-Beta 1 population (**Figures S7E-H**). Consistent with DE analysis, *Fos* and *Jun* were found to be expressed in the Mm-Beta 3 lineage (**Figures S7I-L**). Overall, the trajectory analysis indicated that Mm-Beta 1 was the classical insulin-producing population, which transdifferentiated into Mm-Beta 2 and Mm-Beta 3.

#### Beta-cell cluster patterns in Human islets

We analyzed human beta cells from PancDB and identified three distinct Hs-Beta cell subpopulations (**Figure S8A**). Of these, Hs-Beta 1, exhibited a polyhormonal phenotype characterized by co-expression of *SST* and *GCG*, similar to the Mm-Beta 4 and 2 populations (**Figure S8B**). GSEA revealed that Hs-Beta 1 was enriched in pathways related to translation and signaling receptor regulatory activity pathways. Whereas Hs-Beta 2 showed enrichment in pathways associated with ATP metabolic processes and mitochondrial metabolism (**Figure S8C**). Notably, Hs-Beta 1 cells were predominantly found in obese and T2D individuals, with 75.7% of their differentially expressed genes (n = 2898) shared between these two conditions (**Figure S8D-E**). Key polyhormonal markers such as *GCG*, *PPY*, and *SST* were significantly upregulated in Hs-Beta 1 cells from both obese and T2D subjects compared to healthy controls (**Figure S8F**). In contrast, Hs-Beta 2 cells from T2D subjects exhibited upregulation of genes involved in immune response (*PLCG2*, *CD99*, *JUND*, *HSP90AA1*, *HSPA5*) and mitochondrial processes (*GNAS*, *AGPAT5*, *MT-ND4L*, *MT-ATP8*), while genes related to hormone activity (*SST*, *TTR*, *IAAP*, *NPY*) were downregulated (**Figure S8G**).

### Trans-differentiation of alpha cells into polyhormonal cells under nutritional stress

To explore the alterations in alpha cells during the development of IR, we subclustered alpha cells into Mm-Alpha 1 and Mm-Alpha 2 populations, comprising 75% and 25% of alpha cells, respectively (**Figure 3A**). DEG and pathway analysis showed that Mm-Alpha 1 expresses genes linked to glucagon biosynthesis and secretion (e.g., *Prrc2c, Neurod1, Itpr1, Foxa2*), whereas Mm-Alpha 2 was enriched in ribosomal proteins (Rpl37a, Rpl38Rps27, Smim22) and non-alpha-related endocrine hormones (e.g., *Ins2*, *Ppy*) (**Figure 3B&C, Figure S9A).** Thus, Mm-Alpha 1 can be designated as high glucagon-secreting cells (HGSC), and Mm-Alpha 2 as transdifferentiating alpha cells (TAC). The distribution of these two populations varied significantly across diet, in RC condition, Alpha-1 was the dominant population, while Alpha-2 significantly increased in HFD islets (**Figure 3D; Figure S9B**). Mm-Alpha 1 remained prevalent (90-95%) in early stages (8, 14, 22 weeks, RC; 8, 16 weeks, HFD), but Mm-Alpha 2 dominated after 24 weeks of HFD and 30 weeks of RC feeding, respectively (**Figure 3E; Figure S9C; Table 1**). Given the importance of transcription factors *Pax6*, *Foxa2*, *Arx* and *MafB* for *Gcg* expression in alpha cells (40-42), we compared their expression across the alpha subpopulations and found *Pax6*, *Arx*, and *MafB* to have higher expression in Mm-Alpha 1 compared to Mm-Alpha 2, confirming Mm-Alpha 1 as the primary glucagon-secreting cell population (**Figure 3F**).

**Figure 3.**
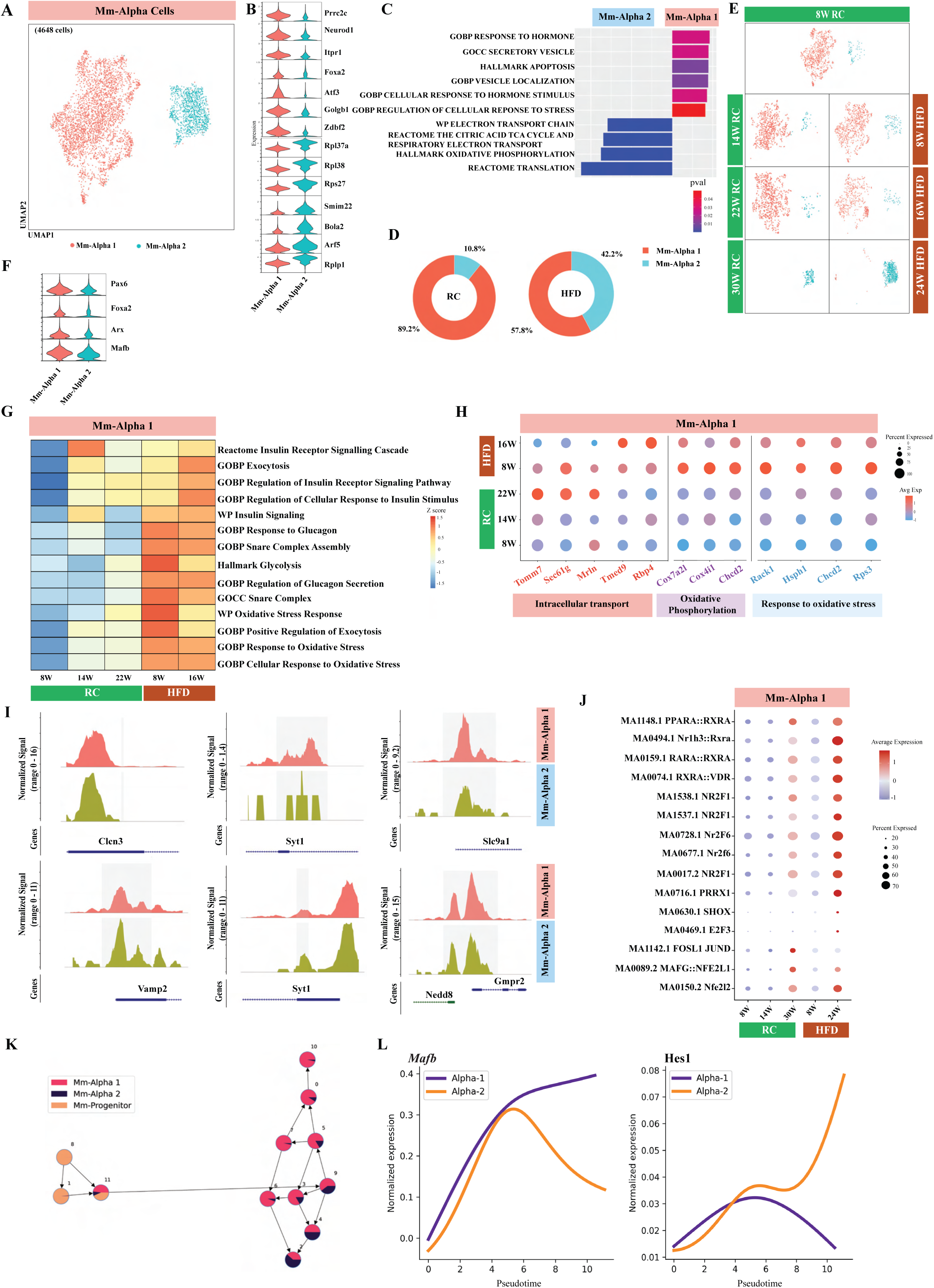
HFD-induced metabolic stress causes alpha cells to adopt a polyhormonal identity. (A) UMAP visualization of pancreatic alpha cell subpopulations (n=4648). Cells are colored by subpopulation identities (Mm-Alpha 1 and Mm-Alpha 2). (B) Expression profile of differentially expressed genes across alpha cell subpopulations. (C) Hallmark pathways enriched in Mm-Alpha 1 and Mm-Alpha 2 subpopulations inferred using GSEA. (D) Distribution of Mm-Alpha 1 and Mm-Alpha 2 subpopulations under RC and HFD conditions. (E) UMAP visualization of Alpha cell subpopulations across different time points across diet conditions. (F) Expression profiles for *Pax6, Foxa2, Arx*, and *Mafb* genes in Mm-Alpha 1 and Mm-Alpha 2 subpopulations. (G) Heatmap of Gene Ontology (GO) or Kyoto Encyclopedia of Genes and Genomes (KEGG) pathways enriched in Mm-Alpha 1 subpopulation across different timepoints under RC and HFD conditions. (H) Expression profiles of genes associated with specific GO and KEGG pathways in Mm-Alpha 1 subpopulation across different timepoints under RC and HFD conditions. (I) Chromatin accessibility profile of differentiable accessible genes (*Clcn3, Syt1, Slc9a1, Syt1*, and *Gmpr2*) in Mm–Alpha 1 and Mm–Alpha 2 subpopulations. (J) Enrichment of motifs as derived from scATAC-seq data in Mm–Alpha 1 cell subpopulation across various timepoints under RC and HFD conditions. (K) Trajectory of Mm-Progenitor, Mm-Alpha 1 and Mm-Alpha 2 cell populations inferred using MARGARET. (L) Gene expression trends for *Mafb* (left) and *Hes1* (right) for Mm-Alpha 1 and Mm-Alpha 2 lineages.

Given its primary role in glucagon secretion, we probed Mm-Alpha 1 further using GSEA across time points and feedings conditions. This analysis revealed significant enrichment of glucagon and insulin signaling related pathways (e.g., glucagon secretion regulation, response to glucagon, insulin signaling), and metabolic processes under dietary stress (**Figure 3G**). Consistent with previous studies linking glucagon secretion to Ca²⁺- and SNARE protein-mediated exocytosis (43, 44), SNARE complex and exocytosis pathways were enriched in Mm-Alpha 1 under HFD condition (**Figure 3G**) (45). These alpha cells further exhibited upregulation of intracellular transport genes (*Tomm7, Sec61g, Mrln, Tmed9, Rbp4*) after 8 and 16 weeks of HFD and after 22 weeks of RC feeding (**Figure 3H).** Genes related to oxidative and cellular stress (*Rack1, Hsph1, Manf, Rps3*) were elevated in Mm-Alpha 1 cells after 8 weeks HFD consumption, indicating early metabolic stress responses (**Figure 3G**). Metabolic pathway analysis in Alpha-1 showed increased expression of oxidative phosphorylation genes (*Cox7a2l, Cox4i1, Chchd2*) after 8 weeks of HFD feeding, followed by a decline at 16 weeks of HFD consumption. Glycolysis-related genes remained enriched at both time points compared to controls (**Figure 3G, H**).

Label transfer from scRNA-seq to scATAC-seq resulted in 1350 mouse alpha cell ATAC-seq profiles, where Mm-Alpha 1 was the primary population in both RC and HFD islets (**Figure S4N, S9D)**. Transcription motifs for *Jun*-*Junb*, *Fosl*-*Jun*, and *Jun* were highly enriched in Alpha-1, indicating enhanced cellular stress handling compared to Mm-Alpha 2 (**Figure S9E**). Mm-Alpha 1 showed higher accessibility of Slc9a1 (glucagon secretion), while vesicle transport genes like *Syt1* and *Vamp2* were more accessible in Mm-Alpha 2 (**Figure 3I**). *Clcn3*, a key insulin secretion regulator, was specifically accessible in Mm-Alpha 2 (**Figure 3I**), aligning with a previous study reporting that *Clcn3*−/− mice exhibited impaired insulin exocytosis and granular acidification (46). In Mm-Alpha 1 ATAC-seq, motifs associated with mitochondrial regulation (*PPARA*: *RXRA*, *Nr1h3:Rxra*, *RXRA:VDR*) and glucose metabolism (SHOX) were highly enriched after 24 weeks of HFD feeding (**Figure 3J**).

Given the expansion of the Mm-Alpha 2 population after 30 weeks of RC and 24 weeks of HFD consumption in our scRNA-seq data, we hypothesized that Mm-Alpha 2 may originate from the Mm-Alpha 1 population. Using MARGARET(39), we inferred the differentiation trajectory of alpha cell types, which was initiated from progenitor cells in the 8W RC condition. This analysis revealed a bifurcating trajectory where a progenitor-enriched cluster (cluster 11) gave rise to intermediate populations (clusters 9, 5, 7, 3, 6), predominantly composed of Mm-Alpha 1 cells. These intermediates further bifurcated into two lineages: one leading to a terminal Mm-Alpha-1 population (cluster 10) and the other containing both Mm-Alpha 1 and Mm-Alpha 2 cells (clusters 4, 2), indicating that the Mm-Alpha 2 population emerges from Mm-Alpha 1 cells (**Figure 3K**). Lineage trend of transcription factors further revealed important transcription factors associated with the Mm-Alpha 1 (e.g., *Atf3*, *Mafb*) and Mm-Alpha 2 lineages (*Hes1*, *Tbx1*) (**Figure 3L, Figure S9F**).

#### Alpha-cell cluster patterns in Human islets

To enable cross species comparison, we performed reference mapping between mouse and human alpha cell populations. Human alpha cells (n=15,412) were subclustered into Hs-Alpha 1 and Hs-Alpha 2 groups. Similar to mouse Mm-Alpha 1, Hs-Alpha 1 expressed *PAX6*, *MAFB*, and *CHGA*, while *INS* was enriched in Hs-Alpha 2 (**Figure S10A–C**). Pathway analysis revealed that Hs-Alpha 1 aligned with Mm-Alpha 1, being involved in vesicle-mediated transport, while Hs-Alpha 2 corresponded to Mm-Alpha 2, showing enrichment in polyhormone genes (*GCG*, *INS*, *SST*), oxidative phosphorylation and ETC pathways (**Figure S10D&E**). As observed with mice, in obese and T2D human subjects, Hs-Alpha 2 populations expanded, while Hs-Alpha 1 decreased (**Figure S10F&G)**. Unlike Mm-Alpha 1, genes for exocytosis and glucagon response were upregulated in control human cells, whereas cellular and oxidative stress pathways were enriched only in obese populations (**Supp Figure S10H)**.

### Delta Cell Heterogeneity and Adaptive Plasticity in Response to Nutritional Stress

Delta cells, the third most abundant endocrine cells in the pancreas, play a key role in the release of islet hormones despite their low abundance. In our murine scRNA-seq dataset, 3008 delta cells were identified under RC and HFD conditions (**Table 1**). Sub-clustering revealed three distinct populations: Mm-Delta 1, Mm-Delta 2, and Mm-Delta3 (**Figure 4A-C**). Under RC conditions, Mm-Delta 1 constituted ∼70%, Mm-Delta 2 ∼24.5%, and Mm-Delta 3 ∼5.75%, respectively. In contrast, the proportions shifted under HFD conditions, with Mm-Delta 2 comprising 50.6%, Mm-Delta 1 30.2%, and Mm-Delta 3 19.1%, respectively (**Figure 4B**). Notably, Mm-Delta 1 remained stable during RC feeding, while Mm-Delta 2 showed minor variations. Under HFD conditions, Mm-Delta 1 increased from 8 weeks to 16 weeks and decreased after 24 weeks. Mm-Delta 2 disappeared at 24 weeks, and Mm-Delta 3 increased significantly at this time point (**Figure 4D, Figure S11A&B**). DEG analysis revealed that Mm-Delta 1 predominantly expressed *Sst* and Sst secretion-associated genes (*Stxbp5l, Syne1*), Mm-Delta 2 expressed beta cell markers (*Ins1, Ins2, G6pc2*), and Mm-Delta 3 expressed *Sst* alongside *Gcg, Pyy, and Ppy* (**Figure 4C**). These populations were characterized as High Sst-Expressing Cells (HSEC, Mm-Delta 1), Beta-Cell Transdifferentiating Delta Cells (BTDC, Mm-Delta 2), and Polyhormonal Delta Cells (PDC, Mm-Delta 3). GSEA showed enrichment of peptide metabolic processes in Mm-Delta 1, hormone secretion and insulin activity in Mm-Delta 2, and increased ribosomal pathway activation with vesicle transport downregulation in Mm-Delta 3 (**Figure 4E**). Increased Mm-Delta 2 proportions in HFD mice suggest that delta cells attempt to compensate for beta cell loss by producing insulin during IR.

**Figure 4.**
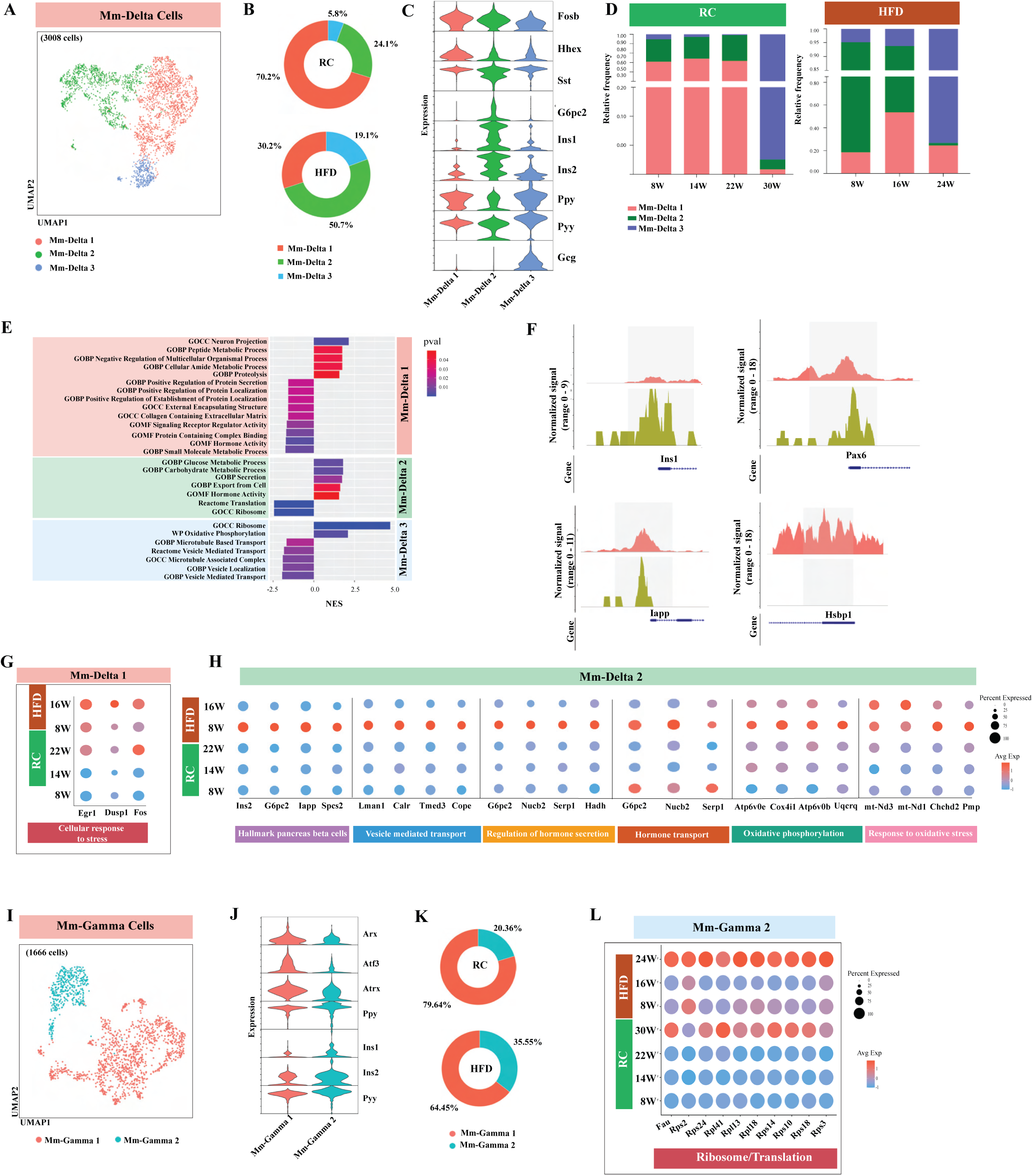
Transcriptomic and epigenomic heterogeneity of delta and gamma cells under HFD exposure. (A) UMAP visualization of mice pancreatic delta cell subpopulations. Cells are colored by subpopulation identities (Mm-Delta 1, Mm-Delta 2, and Mm-Delta 3). (B) Distribution of Mm-Delta 1, Mm-Delta 2, and Mm-Delta 3 subpopulations across RC and HFD conditions. (C) Expression profile of differentially expressed marker genes for mice delta subpopulations. (D) Distribution of Mm-Delta 1, Mm-Delta 2, and Mm-Delta 3 subpopulations across different timepoints across RC and HFD conditions. (E) Hallmark pathways enriched in Mm-Delta 1, Mm-Delta 2, and Mm-Delta 3 subpopulations inferred using GSEA. Positive Normalized Enrichment Score (NES) values indicates upregulated pathways and a negative NES indicates dowregulated pathways, respectively. (F) Chromatin accessibility profile for *Ins1* (left top), *Pax6* (right top), *Iapp* (left bottom), and *Hspb1* (right bottom) in Mm-Delta 1, and Mm-Delta 2 subpopulations. (G) Expression of stress response genes in Mm-Delta 1 cells across different time points across RC and HFD conditions. (H) Expression of genes associated with specific pathways (Hallmark pancreas beta cells, Vesicle mediated transport, Regulation of hormone secretion, Hormone transport, Oxidative phosphorylation, and Response to oxidative stress) in Mm-Delta 2 cells across different time points across RC and HFD conditions. (I) UMAP visualization of mice pancreatic gamma cell subpopulations. Cells are colored by subpopulation identities (Mm-Gamma 1 and Mm-Gamma 2). (J) Expression profile of differentially expressed marker genes for Mm-Gamma 1 and Mm-Gamma 2 subpopulations. (K) Distribution of Mm-Gamma 1 and Mm-Gamma 2 subpopulations under RC and HFD conditions. (L) Expression of genes associated with Ribosome/translation in Mm-Gamma 2 cells across different timepoints across RC and HFD conditions.

ScATAC-seq analysis of 935 nuclei (RC and HFD) revealed distinct chromatin accessibility patterns (**Table 5**). Mm-Delta 2 showed increased chromatin accessibility of *Ins1*, *Pax6*, and *Iapp*, consistent with gene expression data, while Mm-Delta 1 exhibited increased *Hspb1* accessibility (**Figure 4F**). Motif enrichment in Mm-Delta 2 indicated activation of *Sox* and *Atf4*, markers of progenitor-like states and beta cell stress responses (47, 48) (**Figure S11C**).

Due to their low abundance, the role of delta cells in somatostatin secretion and signaling during T2D progression remains poorly understood. Taking advantage of our data, we focused on Mm-Delta 1 and Mm-Delta 2 populations and examined their responses under nutritional stress. *Egr1, Dusp1*, and *Fos* expression increased in Mm-Delta 1 under RC (22 weeks) and HFD (8 and 16 weeks) conditions, correlating with enriched oxidative stress TF motifs (*Fos*, *Jun*) in scATAC-seq data (**Figure 4G, Figure S11D**). Mm-Delta 2 displayed upregulation of beta cell markers and insulin secretion genes (*Ins2, G6pc2, Iapp, Spcs1*), along with genes involved in vesicle transport (*Lman1, Calr*) and hormone regulation pathways after 8 weeks of HFD feeding (**Figure 4H**). Oxidative phosphorylation genes (*Atp6v0e, Cox4i1*) were elevated after 8 and 16 weeks of HFD consumption, alongside with mitochondrial stress genes (*mt-Nd3, mt-Nd1*; **Figure 4H**). These findings highlight that Mm-Delta 2 cells transdifferentiate toward a beta-like phenotype, potentially as a compensatory mechanism under IR. To investigate the lineage relationship between the delta cell populations, we conducted trajectory analysis (MARGARET), which indicated Mm-Delta 2 as a precursor to Mm-Delta 1 and Mm-Delta 3, with Mm-Delta 3 as a terminal transdifferentiating population (**Figure S11E**).

Subclustering of 1466 human delta cells revealed two distinct subpopulations (Hs-Delta 1 and Hs-Delta 2) (**Figure S11F-H**). Hs-Delta 2 showed increased expression of *INS*, *GCG*, and *PPY*, similar to the murine Mm-Delta 2 population (**Figure S11G&H**). GSEA revealed the downregulation of stress pathways, TCA cycle, and oxidative phosphorylation in Hs-Delta 2, while Hs-Delta 1 showed decreased peptide transport and insulin secretion pathways (**Figure S11I**). Hs-Delta 1 was predominant in healthy and T2D individuals, while Hs-Delta 2 increased in the obese state **(Figure S11J)**. These findings align with the mouse data, suggesting an increase in the Hs-Delta 2 population in IR.

Ppy-expressing gamma cells constitute a minor, understudied population in pancreatic islets. Subclustering of 1,666 gamma cells (RC and HFD combined) revealed two distinct subpopulations: Mm-Gamma 1 and Mm-Gamma 2 (**Figure 4I; Figure S12A-B**). Mm-Gamma 1 cells showed elevated expression of *Ppy, Arx, Atf3, and Atrx* genes associated with hormonal secretion, while Gamma-2 cells expressed ribosomal proteins (*Rsp29, Rpl37, Rplp1, Rpl41*), *Ins2, Sst*, and high *Ppy*/*Pyy* levels (**Figure 4J**). Mm-Gamma 1 predominated in early stages (8, 14, 22 weeks of RC; 8, 16 weeks of HFD), while Mm-Gamma 2 was enriched at later stages (30 weeks of RC, 24 weeks of HFD) (**Figure 4K, Figure S12B&C, Table 1**). Pathway analysis revealed that microtubule organization, protein folding, and stress response were upregulated in Mm-Gamma 1, whereas Mm-Gamma 2 showed enrichment of genes involved in translation, RNA metabolism, and OXPHOS pathways (**Figure S12D**). Under HFD conditions, Mm-Gamma 1 exhibited increased enrichment of genes associated with hormone transport, secretion, and glucose metabolism pathways, particularly at 16 weeks of HFD feeding (**Figure S12E**). Mm-Gamma 2 displayed significant upregulation of transcription and ribosomal genes (*Fau, Rsp2/24, and Rpl3*), with stronger expression after 24 weeks of HFD consumption, as compared to 30 weeks of RC feeding (**Figure 4L**).

### Islet vascular cells (IVC) transcriptional and epigenetic changes during the IR state

Pancreatic islet vascular cells (IVCs) play crucial roles in islet development, hormone secretion, and immune cell recruitment. To investigate changes in IVCs during obesity and IR, we subclassified 4025 IVCs (2505 RC; 1520 HFD) into four subclusters, identified as endothelial cells and pericytes (*Notch3, Acta2, Pdgfrb*) (**Figure 5A**). Endothelial cells were further categorized into Mm-Vascular endothelial cells (VECs) (*Rgcc, Car4, Thbs1, Aldh2*), Mm-Progenitor endothelial cells (*Mal, Pbx1, Dkk2, Tgfb2*), and Mm-Lipid-handling endothelial cells (*Fbln2, Cavin2, Ptgs1, Scarb1*) based on DEG analysis (**Figure 5B**). VECs accounted for >50% of the endothelial cell population under RC and HFD conditions, except at 30 weeks of RC and 24 weeks of HFD feeding, where endothelial cells were markedly reduced (**Figure 5C&D; Figure S13A&B**). Mm-Progenitor endothelial cells disappeared with age across conditions, while Mm-Lipid-handling endothelial cells showed an increase at 16 weeks of HFD feeding, followed by a significant decline at 24 weeks of HFD consumption (**Figure 5D; Figure S13B**).

**Figure 5.**
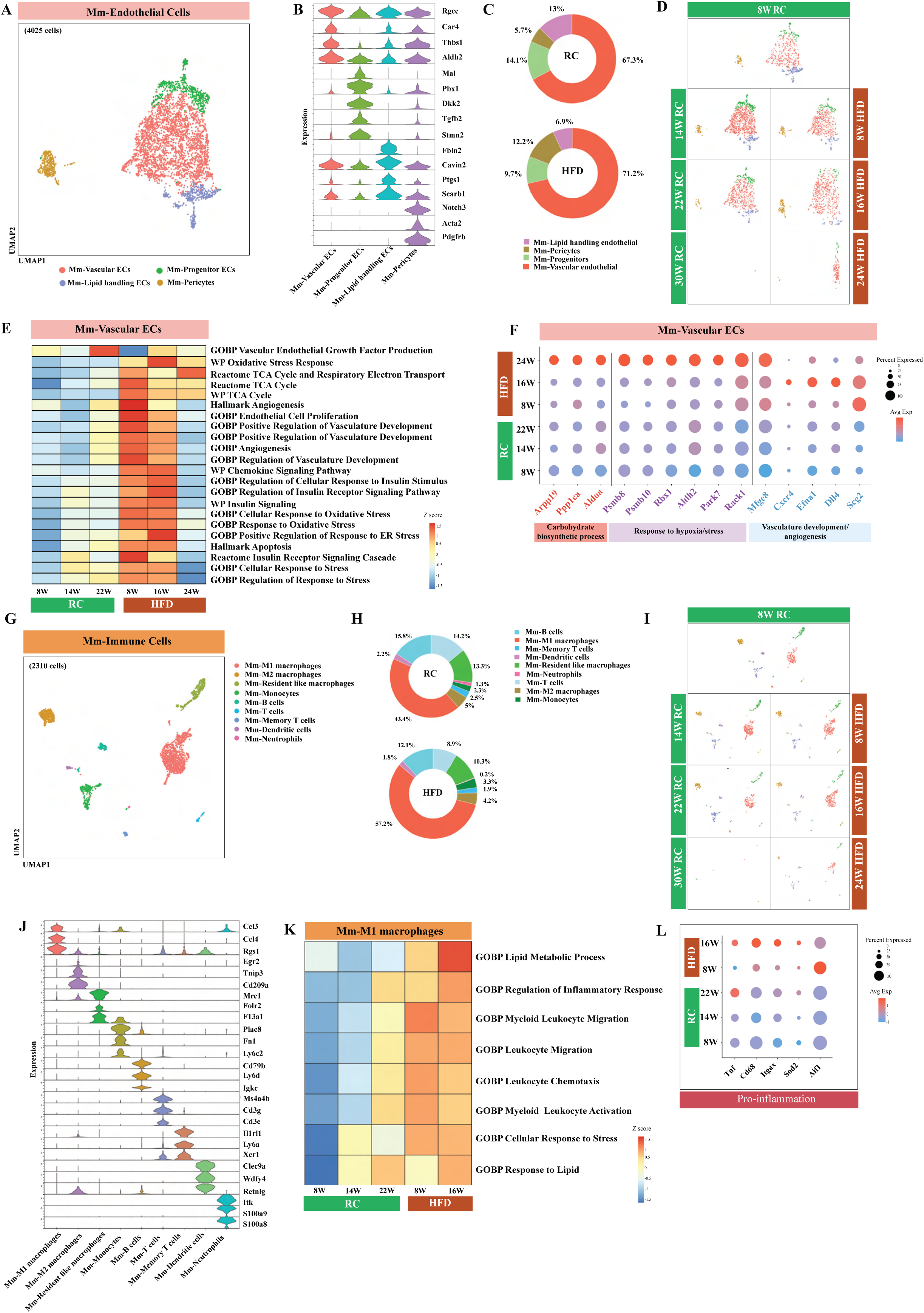
Transcriptomic heterogeneity of murine islet endothelial and immune cells across RC and HFD conditions. (A) UMAP visualization of murine pancreatic endothelial cell subpopulations. Cells are colored by subpopulation identities (Mm-Vascular ECs, Mm-Lipid handling ECs, Mm-Progenitor ECs, and Mm-Pericytes). (B) Expression profile of differentially expressed genes across endothelial cell subpopulations. (C) Distribution of endothelial cell subpopulations under RC and HFD conditions. (D) UMAP visualization of temporal distribution of endothelial cell subpopulations across RC and HFD conditions. (E) Heatmap of pathways enriched in Mm-Vascular ECs across different timepoints across RC and HFD conditions. (F) Expression of genes associated with specific pathways (Angiogenesis, Oxidative stress and Chemokine signaling) in Mm-Vascular ECs across different timepoints across RC and HFD conditions. (G) UMAP visualization of mice pancreatic immune cell types. Cells are colored by immune cell types marker. (H) Distribution of immune cell types across dietary conditions. (I) UMAP visualization of temporal distribution of immune cell types across RC and HFD conditions. (J) Expression profile of marker genes corresponding to immune cell types. (K) Heatmap of pathways enriched in Mm-M1 macrophages across different timepoints across RC and HFD conditions. (L) Expression of proinflammatory pathway-associated genes in Mm-M1 macrophages across different timepoints across RC and HFD conditions.

To further study VECs, we performed GSEA analysis between RC and HFD populations and characterized the changes in major pathways. We observed that pathways like angiogenesis, oxidative stress, and chemokine signaling pathways were upregulated with prolonged HFD feeding (**Figure 5E)**. Blood glucose sensing in endothelial cells is affected in diabetes(49); thus, to understand how the major metabolic pathways regulate glucose metabolism, we studied the expression pattern of genes involved in carbohydrate biosynthetic process (*Aldoa, Arpp19, Ppp1ca*) and oxidative phosphorylation (*Cox4i1, Uqcrq, Uqcrh, Ndufb9*) (**Figure 5F)**. Interestingly, genes involved in these pathways were upregulated under HFD conditions as compared to their age-matched RC mice, and the expression of these genes was significantly upregulated after 24 weeks of HFD feeding. Earlier studies have shown that exposure of endothelial cells to high glucose and free fatty acid (FFA) induces the expression of proinflammatory genes and leads to apoptosis (50). As expected, genes involved in cytokine responses (*Cxcr4, Efna1, Dll4, Scg2, Mfge8*) showed increased expression after 8 and 16 weeks of HFD consumption but a reduction after 24 weeks of HFD feeding (**Figure S13C)**. A similar pattern was observed with pathways like insulin signaling, apoptosis, and angiogenesis/vascular development (*Cxcr4, Efna1, Dll4, Scg2, Mfge8*). However, genes involved in the cellular response to stress/hypoxia (*Psmb8, Psmb10, Rbx1, Aldh2, Park7, Rack1*) showed maximum expression at 24 weeks (**Figure 5F)**.

For cross-species comparison, we analyzed endothelial cells from PancDB human data, where we identified four subpopulations of human endothelial cells: Hs-Vascular, Hs-Metabolically active, Hs-Proinflammatory, and Hs-Immunoregulatory endothelial cells (**Figure S14A-B**). Hs-Metabolically active endothelial cells increased in obesity but decreased significantly in T2D, while Hs-Proinflammatory endothelial cells increased under both conditions (**Figure S14C-D**). The endothelial subclusters were annotated and characterized based on GSEA pathways enriched in the subclusters (**Figure S14E**). Similar to mouse endothelial cells, genes involved in angiogenesis and programmed cell death were enriched in obesity and T2D, whereas peptidase activity was selectively elevated in T2D (**Figure S14F**).

### Transcriptomic heterogeneity of immune cells in pancreatic islets of RC and HFD mice

To investigate immune cell changes in response to diet, we subclustered 2310 immune cells (1411 RC; 899 HFD) into nine subpopulations based on differentially expressed genes: Mm-M1 macrophages (*Ccl3, Ccl4, Rgs1*), Mm-M2 macrophages (*Egr2, Tnip3, Cd209a*), Mm-Resident-like macrophages (*Mrc1, Folr2, F13a1*), Mm-Monocytes (*Plac8, Fn1, Ly6c2*), B cells (*Cd79b, Ly6d, Igke*), Mm-T cells (*Ms4a4b, Cd3g, Cd3e*), Mm-Memory T cells (*Il1rl1, Ly6a, Itk*), Mm-Dendritic cells (*Clec9a, Idh4*), and Mm-Neutrophils (*Retnlg, S100a9, S100a8*) (**Figure 5G-J, Figure S15A**). Mm-M1 macrophages were the most abundant population in both RC and HFD mice, with a 15% increase in number after HFD feeding (**Figure 5H**). B cells were the second most abundant cell type, but were not further analyzed due to low abundance. Interestingly, no significant changes in Mm-M1 macrophage number were observed at 8 weeks of HFD consumption compared to age-matched RC mice. However, after 16 weeks of HFD feeding, the Mm-M1 macrophage population significantly increased, as compared to RC control mice, with no further increase in 24-week HFD mice, indicative of niche saturation in islets (**Figure 5I, Figure S15B**). Our findings are in agreement with a previous study carried out with C57BL/6J mice reporting a doubling of CD11b+ myeloid cells after 8 weeks of HFD feeding (13). Pathway analysis revealed the upregulation of genes involved in inflammatory responses, leukocyte migration, chemotaxis, and myeloid activation after 8 and 16 weeks of HFD consumption (**Figure 5K**). Additionally, genes associated with proinflammation were upregulated in Mm-M1 macrophages, more prominently after 16 weeks than after 8 weeks of HFD feeding (**Figure 5L**). To provide context for our data, we compared our results from M1 macrophages with published findings on intra-islet CD11c+ macrophages, which indicated a positive correlation under HFD feeding (**Figure S15C)** (13). Interestingly, tissue-resident macrophage populations remained largely unchanged under HFD-induced obesity.

We also analyzed immune cell populations in human islets from PancDB, where we identified four subpopulations: Hs-T cells (*CD3D*, *CCL5*, *KLRB1*), Hs-Granulocytes (*TSPB2*, *TPSAB1*, *KIT*), Hs-Macrophages (*CD68*, *LYZ*, *CXCL8*), and Hs-APCs (*HLA*-*DRA*, *HLA*-*DPB1*, *HLA*-*DPA*) (**Figure S16A-C**). Consistent with the mouse data, Hs-Macrophages were the most abundant population and exhibited upregulated inflammatory pathways, leukocyte chemotaxis, and NF-κB signaling in obese and T2D states, whereas antigen presentation and translational processes were enriched under control conditions (**Figure S16D**).

### Cell-cell communication in pancreatic islets of healthy mice and under nutritional stress mice

Pancreatic islets host a complex microenvironment where diverse cells interact to regulate the secretion of hormones critical for glucose and cellular homeostasis (1). Using CellChat (51)we identified intercellular ligand-receptor interactions in healthy (islets from RC mice) and IR states (islets from HFD mice). Analysis of interactions within the endocrine cell populations revealed 41 signaling pathways: 21 were conserved across RC and HFD conditions, 16 were specific to RC islets, and 4 were specific to HFD islets **(Figure 6A)**. Pathways enriched exclusively in RC islets included GIPR (GCG-GIPR), MHC-1, CD200 (Cd200-Cd200r1), THBS (Thbs3-ITGA3_ITGB), and protease-activated receptors signaling, all absent in HFD islets **(Figure 6A**, **Table 6)**. Although insulin signaling was conserved in both states, its overall strength decreased in HFD islets, except for enhanced signaling between Mm-Beta 1 and Mm-Gamma 1 cells **(Figure 6B)**. Four pathways, APP, PDGF, HSPG, and PTN were enriched specifically in HFD islets and absent in RC islets **(Figure 6A)**. APP signaling predominantly involved Mm-Delta 1 as the source of the APP ligand and Mm-Beta 2 as the recipient, with Cd74 as a target receptor **(Figure 6C)**. This finding is consistent with a previous study reporting that Cd74 expression is differentially regulated in diet-induced obesity (48). HSPG and PDGF signaling were mediated by Mm-Beta 1-derived ligands targeting multiple endocrine cells (**Figure S16A**), while Mm-Delta 3 cells uniquely secreted Hspg2 in HFD islets (**Table 6**). Interestingly, Mm-Delta 3 cells displayed exclusive interactions under HFD conditions, including somatostatin, CDH, and HSPG signaling (**Figure S17A-C**). PTN signaling, mediated by Mm-Delta 1-derived Ptn, engaged various receptors across endocrine cells (Ptprz1, Ncl; **Figure S17A-C, Table 6**). While most endocrine cells exhibited reduced incoming and outgoing interaction strength in HFD islets, Mm-Delta 3 cells showed increased incoming and outgoing interaction strength, suggesting that it has a potential role in regulating islet function under diet-induced metabolic stress **(Figure 6D)**.

**Figure 6.**
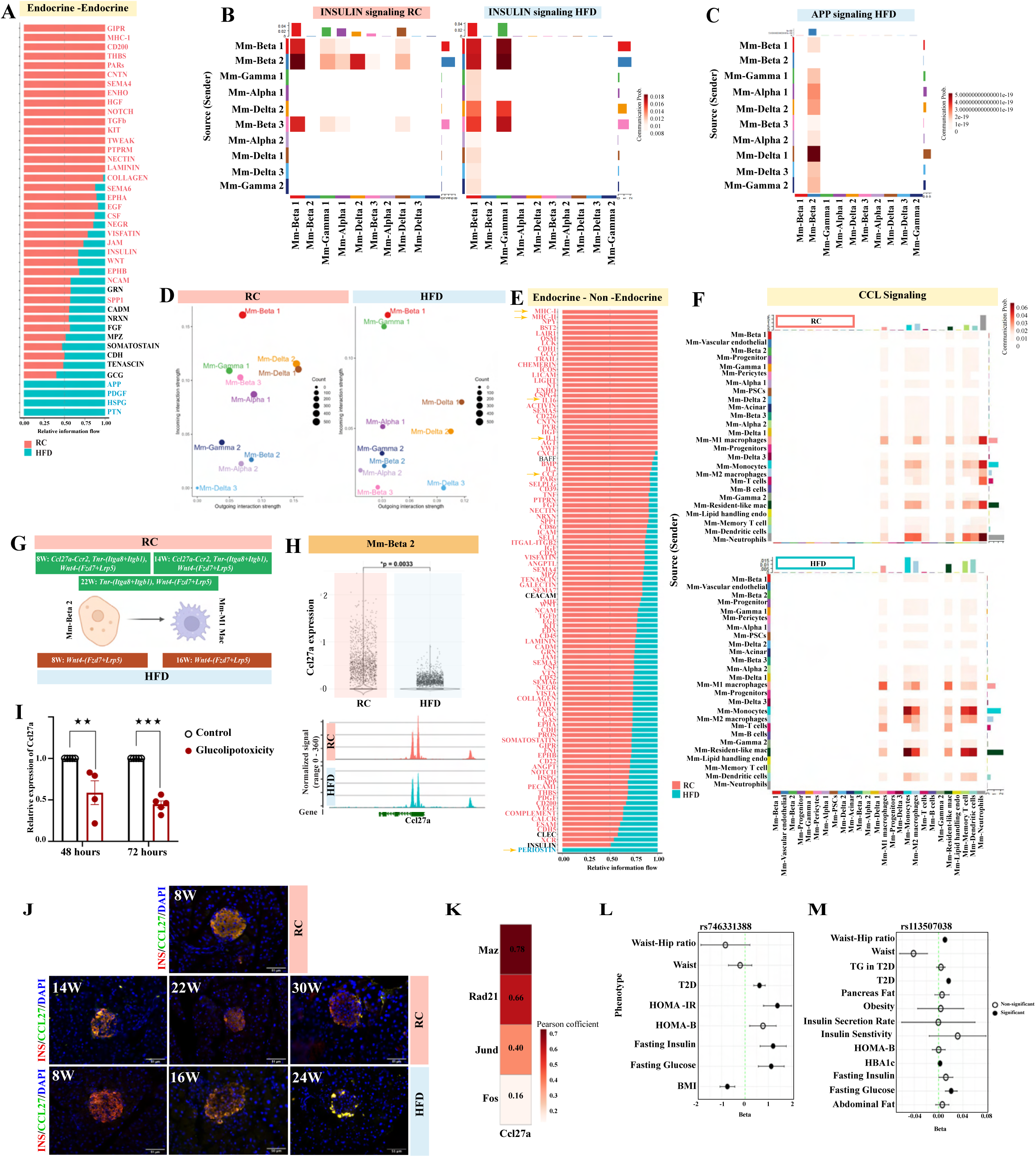
Cell-cell interactions in murine pancreatic islets under different metabolic conditions. (A) Comparison of overall information flow for the top signaling pathways inferred for communication within the endocrine cells across RC and HFD conditions. (B) Heatmap representing insulin signaling among endocrine cells under RC (left) and HFD (right) conditions. (C) Heatmap representing APP signaling among endocrine cells under HFD conditions. (D) 2D projection of incoming/outgoing interaction strength among endocrine cell types under RC and HFD conditions. (E) Comparison of overall information flow for the top signaling pathways inferred for communication within endocrine and nonendocrine cells across RC and HFD mice. (F) Heatmap representing CCL signaling, within endocrine and nonendocrine under RC and HFD conditions. (G) Graphical representation of ligand-receptor interactions between Mm-Beta 2 and Mm-M1 macrophages in RC and HFD mice at different time points. (H) Comparison of *Ccl27a* gene expression and chromatin accessibility in Mm-Beta 2 under RC and HFD conditions. (I) Comparison of relative expression of *Ccl27a* in *Ins1* cells upon treatment with glucolipotoxic media for 48 and 72 hours. (J) Immunofluorescence staining of pancreatic slices. Slices were co-stained with either an anti-CCL27 antibody (Alexa Fluor, green) and an anti-insulin antibody (Alexa Fluor, red). The inserts in the upper right of each panel show enlarged islets areas. Scale bar in inserts: 51 μm: Contrast was adjusted for improved visualization. (K) Heatmap of correlation of expression of *Ccl27a* and transcription factors (*Maz, Rad21, Jund, Fos*) in Mm-Beta (RC mice). (L) Association of SNPs (rs746331388) in *Ccl27a* with anthropometric and metabolic traits in Common Metabolic Disease Knowledge Portal. The filled circles are significant after correction (p < 1e−5). (M) Association of SNPs (rs113507038) in *Ccr2* with anthropometric and metabolic traits in Common Metabolic Disease Knowledge Portal. The filled circles are significant after correction (p < 1e−5). TG, Triglycerides; T2D, Type 2 Diabetes.

Using CellChat, we next analyzed interactions between endocrine and non-endocrine cells, identifying 104 signaling pathways. Of these, 27 were specific to RC islets, 1 was unique to the HFD islets, and the remaining pathways were conserved across both states **(Figure 6E)**. Immune cell types, including macrophages, dendritic cells, neutrophils, and monocytes, showed reduced interaction strengths in HFD islets. Interestingly, periostin signaling emerged in HFD islets, linking PSCs and M1 macrophages as interacting partners **(Figure S18A).** Periostin modulates adipose tissue macrophage infiltration (52), and deficiency of periostin in mice showed increased sensitivity towards streptozotocin (53). Notably, pathways such as MHC-I, MHC-II, interleukin (e.g., IL1, IL16) were completely absent in HFD islets **(Figure 6E, Figure S18B and C),** and cytokine and chemokine signaling was significantly reduced in HFD islets (**Figure 6E**). CCL27a-mediated chemokine signaling, observed between endocrine and immune cells (Mm-Beta 2 and Mm-M1 macrophages) under RC conditions, was entirely absent in HFD islets **(Figure 6F & G, Figure S18D)**. We further analyzed Ccl27a RNA expression and chromatin accessibility using scRNA-seq and scATAC-seq data, respectively. Consistent with our cell-cell interaction analysis, Ccl27a expression and chromatin accessibility were significantly reduced in Mm-Beta 2 cells in HFD islets **(Figure 6H)**. To further validate our findings, we conducted in vitro studies using INS-1 cells, where lipotoxic and glucotoxic conditions were induced to mimic HFD exposure (54). Following 48 and 72 hours of treatment, INS-1 cells exhibited a significant decrease in the expression of Ccl27a, in agreement with our single cell data **(Figure 6I)**. We confirmed from immunohistochemical imaging the colocalization of insulin and Ccl27 at all different dietary timepoints **(Figure 6J)**. Consistent with our findings from cell-cell communication analysis, we found the colocalization was more prominent in the RC 8 week condition. The expression of Ccl27 was reduced with HFD feeding (**Figure 6J, Figure S19A,B**). We further investigated the transcription factors that might regulate the expression of Ccl27a in Mm-Beta 2 cells using SCENIC (55) and identified Maz, Rad21, Jund, and Fos as potential regulators **(Figure 6K**, **Table 7)**.

The role of the Ccl27-Ccr2 axis in the context of obesity, IR, or T2D remains unexplored. To assess the translational relevance of our findings, we analysed the correlation of CCL27 gene expression and genetic associations with anthropometric traits. Within the CCL27 genetic locus, a missense single nucleotide polymorphism (SNP), rs746331388, was identified as being positively associated with multiple metabolic traits, including T2D, Homeostasis Model Assessment of Insulin Resistance (HOMA-IR), and fasting insulin and glucose **(Figure 6L)**. Similarly, analysis of the CCR2 genetic locus revealed an upstream variant, rs113507038, which was positively correlated with waist-hip ratio, T2D, HbA1c, and fasting glucose **(Figure 6M)**.

## Discussion

In the current study, we used single-cell sequencing technology to provide a high-resolution transcriptional and epigenomic map of mouse islets under healthy and nutritional stress conditions. While previous investigations have characterized islet cell heterogeneity under metabolic challenges (25, 29, 56, 57), our longitudinal approach characterized temporal cellular and molecular changes, unveiling distinct adaptive trajectories across endocrine and non-endocrine cell populations.

Our findings underscore the remarkable plasticity of islet endocrine cells in response to metabolic perturbations. Specifically, we identified the transdifferentiation of endocrine cells as a crucial adaptive mechanism in response to metabolic stress. Under HFD conditions, alpha (Mm-Alpha 2) and delta (Mm-Delta 2) cell subpopulations acquired beta-like identities, marked by upregulated *Ins1*, *Ins2*, and *Pdx1* expression. This phenomenon aligns with prior reports demonstrating alpha to beta cell conversion under beta cell loss or stress conditions (19, 58-60). For example, Thorel et al. (2010) demonstrated that alpha cells act as a reservoir for beta-cell regeneration in mice with diphtheria toxin-induced beta-cell loss (58). Moreover, Chakravarthy et al. (2017) demonstrated that alpha-to-beta transdifferentiation can be induced by targeting epigenetic regulators such as *Arx (19),* while the overexpression of Pax4 also facilitates the reprogramming of alpha cells into insulin-producing cells (61).

We further established that delta cells exhibit a distinct dual response under metabolic stress condition. Alongside beta-like transdifferentiation, a polyhormonal subpopulation emerged that co-expressed somatostatin, insulin, and glucagon, among other hormones. This dual phenotype is consistent with prior evidence from zebrafish and STZ-treated mice, where delta-to-beta reprogramming has been observed following beta cell loss (62). The presence of polyhormonal delta cells also mirrors findings in human T2D islets, where mixed-hormone expression patterns have been reported (63, 64). Importantly, this polyhormonal identity was associated with significant downregulation of *Pax6*, a transcription factor essential for maintaining delta cell fate and overall islet endocrine identity (65). Reduced *Pax6* expression may permit the destabilization of delta cell identity and enable the co-expression of non-canonical hormones.

Interestingly, while polyhormonal gamma cells showed no beta-like reprogramming, they displayed a broadening of their secretory profile, consistent with their role in modulating islet function through PPY and peptide YY (66). This observation suggests that gamma cells retain their identity but adapt their function to metabolic challenges, contributing to the regulatory intra-islet network. This selective plasticity may be due to developmental lineage constraints, as gamma cells derive from progenitors distinct from those that give rise to alpha, beta, and delta cells (67). Our findings show that, despite being prone to dedifferentiation under metabolic stress, beta cells did not transdifferentiate into other endocrine types, a pattern consistent with previous reports emphasizing identity loss rather than lineage conversion(15, 31, 68). The decline of the canonical Mm-Beta 1 population and the rise of Mm-Beta 2 and Mm-Beta 3 populations highlight the plasticity of beta cells under obesogenic stress, consistent with early signs of dedifferentiation and inflammation in diabetes (17, 31). Mm-Beta 2 cells increased ribosomal activity alongside reduced insulin signaling pointing to a metabolic shift likely prioritizing biosynthesis over secretion, mirroring known stress adaptations in nutrient-overloaded beta cells (69). The temporary emergence of Mm-Beta 3, defined by stress markers (*Fosb*, *Jun*) and NF-κB activation, reflects acute glucolipotoxic stress, while its later disappearance may signal either successful adaptation or a slide into irreversible dysfunction, a critical transition that warrants closer study. Mm-Beta 2-2 resemblance to beta cells from db/db and STZ models underscores shared stress responses, particularly ER overload and disrupted protein folding (*P4hb*, *Hsp90b1*), classic features of T2D-related beta cell failure (70). In contrast, Mm-Beta 2-1 cells distinct polyhormonal profile (*Ppy*, *Sst*) and divergence from diabetic states may reflect a short-term dedifferentiated phase, typical of early hyperglycemia adaptation (31). The progression from Mm-Beta 2-1 to Mm-Beta 2-2 with prolonged HFD consumption suggests increasing mitochondrial and ER stress, accompanied by reduced *Ins1/2* chromatin accessibility, indicating deepening epigenetic disruption. Interestingly, Mm-Beta 1 subpopulations also diverge functionally, with Mm-Beta 1-1 focused on insulin secretion and Mm-Beta 1-2 enriched in antioxidant genes, suggesting internal specialization to manage secretion and stress (56). The presence of a similar polyhormonal profile in human Hs-Beta 1 cells from obese/T2D donors supports a conserved adaptive mechanism, while stronger immune activation in Hs-Beta 2 suggests species-specific responses shaped by the human islet environment (57).

The adaptive responses of non-endocrine islet cells, such as endothelial and immune cells, are crucial for understanding how metabolic stress from HFD feeding islet function and the islet compensatory mechanisms. We found that a specific group of endothelial cells, known as VECs, undergo significant metabolic reprogramming when exposed to HFD. This reprogramming is marked by enhanced activity in oxidative phosphorylation (OXPHOS) and carbohydrate biosynthesis pathways. This metabolic shift likely represents an adaptation to increased energy demands for vascular remodeling and nutrient delivery to insulin-producing beta cells, especially as hyperglycemia and lipotoxicity place pressure on the islet microvasculature. Previous research has indicated that endothelial glycolysis and OXPHOS play vital roles in vascular development, suggesting that these metabolic changes in VECs may initially promote islet vascularization in the early stages of HFD exposure (71). However, as metabolic stress becomes prolonged, this balance is disrupted, which is reflected in the decline of all endothelial cell subtypes. A comparable phenomenon is seen in lipid-handling endothelial cells, which are abundant following short-term HFD consumption but diminish with extended exposure to HFD, indicating a decline in endothelial function. Additionally, progenitor endothelial cell (PEC) reduction during chronic HFD consumption aligns with clinical observations that obesity and IR interfere with PEC-mediated vascular repair (72). This decline further aggravates the rarefaction of islet microvasculature and contributes to beta-cell dysfunction. These findings highlight the essential role of vascular dysfunction in the adaptation and failure of islets under metabolic stress (6, 73).

Similar to endothelial cells, HFD feeding triggered significant immune cell remodeling in islets, characterized by an increase in pro-inflammatory Mm-M1 macrophages. These macrophages show heightened lipid metabolism pathways and inflammatory signaling processes, which play a crucial role in islet inflammation. This finding is in agreement with earlier research by Lumeng et al. (2007), demonstrating that obesity causes M1 polarization in adipose tissue macrophages, thereby contributing to systemic IR (74). In the islets, lipid-enriched macrophages are likely to induce beta-cell toxicity through cytokine-driven apoptosis and endoplasmic reticulum stress, as shown in murine models of lipotoxicity (17, 18). Notably, the plateau observed in M1 macrophage numbers during prolonged HFD may suggest niche saturation. This phenomenon has been observed within adipose tissue, where the infiltration of macrophages reaches its peak once the lipid storage capacity is exceeded (75). This saturation could restrict further immune cell recruitment while maintaining a chronic inflammatory environment, which accelerates beta-cell dedifferentiation and loss (76). Importantly, the interaction between metabolic and immune stressors is two-way: lipotoxicity activates macrophages and hinders their shift to anti-inflammatory M2 states, in agreement with a study by Kratz et al. (2014) (77), in obese adipose tissue. In islets, this imbalance may worsen inflammation-related fibrosis and amyloid deposition, which are hallmarks of advanced T2D (78, 79).

A key novel aspect of our study is the characterization of cell-cell communication within the islet microenvironment, which plays a critical role in maintaining glucose homeostasis. We identified a significant disruption in immune-endocrine communication within islets prepared from HFD mice, marked by the loss of key signaling pathways such as MHC-I, MHC-II, and interleukin-mediated interactions. This signaling deficit may impair immune surveillance and contribute to islet dysfunction under metabolic stress (80). Conversely, HFD-specific pathways, such as APP and periostin signaling, may reflect adaptive responses to chronic nutrient overload, with Mm-Delta 3 cells emerging as key regulators through somatostatin and HSPG interactions. HFD-stressed Mm-Beta 2 cells exhibited reduced *Ccl27a* expression, mirroring our findings with INS1 cells grown under glucolipotoxic conditions. Earlier studies showed that *Ccl27*a is expressed in keratinocytes and adipose tissue, playing a role in tissue regeneration and wound healing (81). However, the function of *Ccl27a* in islet tissue remained unclear so far. The difference in *Ccl27a* expression suggests a new mechanism of islet dysregulation. Given the association of *CCL27* and *CCR2* genetic variants with IR and T2D (82), these findings are of significant potential translational relevance.

## Limitations

While our study provides a detailed atlas of islet cell adaptations under HFD stress, its cross-sectional design limits causal inferences about cell-state transitions. Longitudinal studies are needed to track the dynamics of cellular plasticity over time. Furthermore, our findings are based on a murine model, which may not fully recapitulate human islet biology. Finally, while scRNA-seq and scATAC-seq provide high-resolution data, integrating these findings with proteomic and metabolomic analyses would offer an even more comprehensive understanding of islet adaptations to metabolic stress.

## Acknowledgments

We thank Dr. Yinghon Cui (Molecular Signaling Section, NIDDK) for technical assistance in procuring mouse models, and H. Smith, I. Akan, and S. Yun (NIDDK Genomics Core) for their support with RNA sequencing. We are grateful to Dr. Prosenjit Mondal (IIT-Mandi) for generously providing INS-1 cells. We also acknowledge the members of the MMCSL laboratory for their valuable discussions and critical input throughout the study. Research in the SPP laboratory is supported by the DBT/Wellcome Trust India Alliance (IA/I/21/1/505613), the Department of Biotechnology (DBT) [B1/PR44526/MED/30/2376/2021], the Department of Science and Technology - Science and Engineering Research Board (DST-SERB) [CRG/2021/004502], the Indian Council of Medical Research (ICMR) [5/4/8-18/Obs/SPP/2022-NCD-II], and an Initiation Grant from IIT Kanpur. HZ is supported by an IIT Kanpur Initiation Grant [IITK /CS /2019236], DBT/Wellcome Trust India Alliance Early Career Fellowship [IA/E/21/1/506298], Science and Engineering Research Board (SERB), Government of India Startup Research Grant [SRG/2020/001333], and Har Gobind Khorana Innovative Young Biotechnologist Award Grant [BT/13/IYBA/2020/05]. The JW laboratory is funded by the Intramural Research Program of the NIDDK, NIH (Bethesda, MD, USA). SS acknowledges support from the DBT Junior Research Fellowship (DBTHRDPMU/JRF/BET/-20/1/2020/AL/352). KPM acknowledges support from Prime Minister’s Research Fellowship [MOE/CS/PMR9009]. Authors utilized digital tools, including Grammarly and AI-assisted platforms, for the refinement of grammar and sentence construction.

## Author contributions

SPP and HZ designed the study. SS, LFB, JW, and SPP conceived and designed the in vivo and in vitro experiments. MKP, JT, AS, SA, SS, and HZ performed all computational analyses. SS and ST conducted tissue staining and imaging. AG carried out the GWAS analysis. SS, MKP, LFB, JW, HZ and SPP contributed to manuscript writing and revisions. All authors approved the manuscript.

## Declaration of interests

The authors declare no competing interests.

## Supplementary data

Supplementary figures and tables will be provided upon request.

## Materials and Methods

### Mice

All experiments were conducted using male C57BL/6NTac mice (Taconic Biosciences). To establish a diet-induced obesity model, mice were fed a high-fat diet (Research Diet: D12492, 60 kcal % fat) starting at six weeks of age for durations of 8, 16, or 24 weeks. Control groups were maintained on a standard chow diet (Safe Diet: D131, 12.6 kcal% Fat). Body weight was measured weekly for all mice. Mice were housed under standard conditions with a 12-hour light/dark cycle at a constant room temperature (23°C) and ad libitum access to food and water. The research described in this manuscript complies with all relevant ethical regulations. All animal experiments were approved by the National Institute of Diabetes and Digestive and Kidney Diseases/NIH Animal Care and Use Committee or Institutional Animal Ethics Committee (IAEC), Indian Institute of Technology Kanpur.

### In vivo metabolic test

Metabolic tests were performed with mice fed either regular chow (RC) (8, 14, 22, and 30 weeks) or a high-fat diet (HFD) (8, 16, and 24 weeks), following previously described protocols (83). For the glucose tolerance test (GTT), mice were fasted overnight, injected intraperitoneally with 1 g/kg glucose, and blood glucose levels were measured at 0, 15, 30, 60, 90, and 120 min. For the insulin tolerance test (ITT), mice were fasted for 6 hours and then injected with 1 U/kg insulin (Humulin, Eli Lilly) intraperitoneally, followed by monitoring blood glucose at the same time intervals.

Additionally, fasting and fed blood glucose levels and plasma insulin, glycerol, and triglycerides were quantified at all seven time points using commercially available kits, as detailed in the resource table.

### Islet Isolation

Islets were isolated from the pancreata of RC and HFD mice after collagen digestion as described (84). Briefly, the pancreatic duct was cannulated with a 27-G needle connected to a 5 cc syringe, and the pancreas was injected with 3–5 ml of cold (4 °C) enzyme solution (HBSS with 25 mM HEPES buffer and collagenase Type XI; Sigma, Cat. C7657). The pancreas was removed and digested at 37 °C for 12-15 min. Enzymatic activity was terminated by adding cold (4 °C) RPMI (10%). Islets were then washed by spinning and decanting the supernatant. Finally, islets were purified by Ficoll density gradient centrifugation.

### Single-cell suspension and library preparation

Islets were isolated by first pre-warming 0.05% Trypsin, 100% FBS, and Prodo+ media to 37°C for 20 min. Islets were transferred into a 50 mL conical tube and washed with 10 mL of Prodo+ media. After centrifugation at 200 x g for 4 min, the islets were resuspended in 10 mL of Prodo+ media and handpicked under a microscope. The islets were transferred to a 15 mL conical tube with 10 mL of 1X PBS (no Ca²⁺, Mg²⁺), centrifuged at 180 x g for 2 min, and the supernatant was discarded. Islets were then treated with 1 mL of 0.05% Trypsin at 37°C, pipetted to dissociate cells, and incubated at 37°C for 9 min with periodic pipetting every few minutes. The reaction was stopped with 1 mL of FBS, and cells were filtered through a BD FACS tube with a strainer top and then rinsed with PBS + 10% FBS. Cells were centrifuged at 400 x g for 4 min, washed twice with PBS + 10% FBS, and resuspended in the appropriate volume with PBS containing 10% FBS for further single-cell RNA and ATAC library preparation.

Single-cell suspensions with >90% viability were used for library preparation. Single-cell 3ʹ gene expression libraries were generated using the Chromium Single Cell 3ʹ GEM, Library & Gel Bead Kit v3 (10x Genomics, PN-1000075) following the manufacturer’s protocol. Briefly, cells were loaded onto the Chromium Controller to generate Gel Beads-in-Emulsion (GEMs), where individual cells were co-encapsulated with barcoded gel beads in oil droplets, enabling cell-specific transcript tagging. Subsequently, reverse transcription and cDNA amplification were performed according to the 10x Genomics protocol. The resulting amplified cDNA libraries were purified and assessed for quality and size distribution using the Agilent Bioanalyzer 2100. Quantified libraries were then sequenced on an Illumina NovaSeq platform using paired-end sequencing with a single indexing strategy. Sequencing was performed to a depth of at least 20,000 read pairs per cell to ensure robust transcriptome coverage.

Single-cell chromatin accessibility profiling was performed using the Chromium Next GEM Single Cell ATAC Library & Gel Bead Kit v1.1 (10x Genomics, PN-1000175), following the manufacturer’s protocol. Nuclei were isolated from dissociated islets using the 10x Genomics Nuclei Isolation Kit. Subsequently, nuclei were encapsulated with barcoded gel beads in oil droplets using the Chromium Controller. In-GEM transposition and barcoding of accessible chromatin regions were carried out using the Tn5 transposase provided in the kit. Following GEM incubation and breakage, barcoded DNA fragments were purified and PCR-amplified to generate single-cell ATAC-seq libraries. Library quality was assessed using an Agilent Bioanalyzer 2100 to confirm appropriate fragment size distribution. Sequencing was performed on an Illumina HiSeq platform using paired-end, dual-indexed reads, with a sequencing depth of approximately 25,000 read pairs per nucleus.

### INS-1 cell culture and lipoglucotoxic treatment

INS-1 cells (kindly provided by Dr. Prosenjit Mondal at the Indian Institute of Technology, Mandi) were cultured in RPMI supplemented with 10% FBS, 10 mg/ml penicillin/streptomycin, 10 mM HEPES, 50 mM beta-mercaptoethanol, and 1 mM sodium pyruvate at 37°C and 5% CO2. Lipoglucotoxic media was supplement with 30 mM glucose and 0.25 mM palmitate for 48 hours and 72 hours. Subsequently, RNA was isolated using ReliaPrep™ RNA Miniprep Systems (Promega) and subjected to cDNA preparation using Takara PrimeScript™ 1st strand cDNASynthesis Kit and qRT-PCR using TB Green® Premix Ex Taq™ II studies.

### Pancreas staining and imaging

Briefly, mouse pancreata were fixed overnight in 4% paraformaldehyde/phosphate-buffered saline, followed by embedding in paraffin. Pancreatic sections (10 μm thick) were mounted on slides. To determine colocalization of Ccl27a and insulin, three distinct sections per pancreas were blocked with normal donkey serum for 1 h and incubated overnight at 4 °C with a rabbit anti-insulin antibody and a goat anti-Ccl27a antibody. The insulin and Ccl27a antibodies were detected with Alexa Fluor 594 donkey anti-rabbit (red color) and Alexa Fluor 488 donkey anti-goat (green color) secondary antibodies, respectively. All sections were counterstained with DAPI (Vectashield mounting medium with DAPI, Vector Laboratories) to visualize the nuclei (blue color). Slides were imaged using a Leica DM5000B equipped with a DFC420 camera. Image acquisition and merging were performed using ImageJ. Islets from three mice per genotype were analyzed (8 islets per mouse).

### Single-cell RNA-seq data analysis

#### Setup the Seurat Object

In the first step, sequenced data were generated by the 10x Genomics pipeline. The Read10X_h5() function from the Seurat package V3 (85) was used to read the filtered_feature_bc_matrix.h5 matrices generated by 10x Genomics pipeline, which consist of unique molecular identifier (UMI) count matrix of the genes across the cells for each replicate. Next, this count matrix was used to create a Seurat object. For consistency, the individual Seurat objects were made to have the same number of genes across different mice and time points by adding the zero count values to the missing genes. The pre-processing, quality control and decontamination steps were applied to individual objects separately.

#### Pre-processing and quality control

The pre-processing consisted of two main steps, 1. Filtering low-quality cells based on empty droplet detection, total UMI counts, and mitochondrial gene expression, and 2. Filtering low-quality genes based on the expression across cells. First, empty droplets were detected using the emptyDrops() function from the DropletUtils package (86) with a lower parameter set to 2000, and cells with FDR less than 0.05 were retained for further analysis. Subsequently, the low-quality cells were detected based on the quickPerCellQC() of the scuttle package (87) that evaluates low-quality cells based on commonly used quality control (QC) metrics consisting of total UMI counts, UMIs detected, and mitochondrial gene expression percentages. Later, we filtered low-quality genes that were expressed in fewer than 10 cells across all datasets.

#### Decontamination of single-cell RNA-seq data

To perform decontamination of the single-cell RNA-seq data from the ambient RNA, high ambient RNA load was estimated and removed using DecontX (88). Each count matrix for a specific mouse, time point, and diet was used as input to DecontX which resulted in a decontaminated matrix of true expression and contamination fraction. Hyper-parameter delta for DecontX was set to (10, 20) for our dataset.

#### Normalization of data

After quality control, the datasets were normalized using a default global-scaling normalization method, Seurat::LogNormalize, with a scale factor of 10,000. Then batchelor::multiBatchNorm (89) was used to remove the differences in the coverage across the batches.

#### Data Integration and clustering

After normalization, we merged all the individual datasets across different mice and timepoints into a single Seurat object which was used for integration. We first selected the top 2,000 highly variable genes using ‘FindVariableFeatures()’ function, followed by scaling with ScaleData() function, and finally identified the principal components (PCs) using the RunPCA() function. These PCs served as input to the Harmony integration algorithm (90), which outputs the integrated embeddings removing batch effects. Harmony integration was performed using RunHarmony() function by setting group.by.vars parameter to mice replicates. Harmony embeddings were subsequently used for clustering the cells and 2D visualization using the Uniform Manifold Approximation and Projection (UMAP)(91). The clustering was performed using the Louvain algorithm [9] and the optimal resolution was chosen based on the visual inspection of expression of marker genes for the expected cell types. The functions FindNeighbors() and FindClusters() from SeuratV3 were utilized for performing the unsupervised graph-based Louvain clustering.

#### Cell type annotation

We annotated different cell types based on the expression levels of differentially expressed genes and canonical marker genes, visualized on UMAP plots across the Louvain clusters identified using FindClusters() over Harmony embeddings. We identified differentially expressed genes for each cluster using FindAllMarkers() function. We further sub-clustered each cell type and performed sub-clustering within the cell type to identify cell subtypes. We annotated and identified cell types and subtypes by referencing markers from the literature.

#### Differential Expression (DE) analysis across diet and timepoint

To find differentially expressed genes between two groups of cells, we used the limma (92) package. The first group consisted of cells for which we wanted to identify differentially expressed genes, while the second group acted as a control for DE analysis. We then applied lmFit(), eBayes() and topTable() functions from the limma package with default parameters to identify the DE genes. We conducted cell-type, diet, and timepoint-specific DE analysis using limma. DE genes were visualized on volcano plots created using the EnhancedVolcano R package (https://github.com/kevinblighe/EnhancedVolcano).

#### Pathway enrichment analysis

For pathway enrichment analysis, FGSEA (Fast Gene Set Enrichment Analysis) (93) was used to identify upregulated and downregulated pathways. We used fgsea() with the stats parameter set to gene list ranked based of log2 fold change (log2FC) of DE analysis. The pathway parameters were set using MSigDB Collections (https://www.gsea-msigdb.org/gsea/msigdb/mouse/collections.jsp) for mouse species.

#### Computation of gene signature score

We employed VISION (94), a tool designed for gene signature analysis to assess gene signature scores across different conditions of Mm-Alpha and Mm-Beta populations. We selected cells of interest (Mm-Alpha, Mm-Beta) from our dataset and provided their gene expression matrix as input for VISION. Additionally, we provided integrated lower-dimensional embeddings, computed using Harmony, to account for batch effects. We utilized predefined gene sets from the MSigDB Collections (https://www.gsea-msigdb.org/gsea/msigdb/mouse/collections.jsp) for gene signature scoring. VISION computed an overall score for each cell by summarizing the expression levels of genes within each signature. We applied min-max scaling normalization to the computed scores and aggregated the normalized scores for each group by calculating the mean.

#### Trajectory analysis

For trajectory inference, we initially performed preprocessing steps on the subset of cells selected for trajectory inference which usually consisted of the starting cell population i.e. progenitor cells and an endocrine cell population (alpha, beta or gamma). We performed preprocessing on the subset of cells and computed harmony embeddings. Using the computed harmony embeddings on the subset of cells, we ran the MARGARET (39) trajectory inference algorithm using the default parameter setting i.e. “n_episodes” = “10”, “optimizer” = “SGD”, “batch_size” = “256”, learning rate = “0.01”, “n_neighbours” = “30” and “device” = “cuda”. For the “metric_clusters” parameter, we utilized the existing sub-cluster level annotation information, and for the “obsm_data_key” parameter, we used the existing harmony embeddings. We provided the starting cell population as “progenitor cells” for running the algorithm. We used MARGARET’s compute_connectivity_graph() and compute_undirected_cluster_connectivity() to prepare a connectivity graph over cluster nodes inferred by MARGARET. We added the feature of piechart representation at each cluster node for a better representation of population composition at a cluster node level. The terminal cluster ids were calculated using get_terminal_states() function, and the function compute_trajectory_graph() was used for generating the final directed trajectory. We further used compute_pseudo_time() and compute_diff_potential() to compute pseudotime and differentiation potential for all cells. We finally employed the MARGARET’s LinearGAM module to analyse the non-linear gene expression dynamics along the trajectories using the inbuilt function plot_lineage_trends(). The lineage trend analysis was performed for all the mouse transcription factors available in the FANTOM repository (95). https://fantom.gsc.riken.jp/5/sstar/Browse_Transcription_Factors_mm9).

### Mm-ATAC sequencing dataset

#### Data Preprocessing and QC

Raw fastq files from multiple batches were aligned separately to the mm10 reference genome and quantified using CellRanger ATAC v1.2.0 (10x Genomics http://10xgenomics.com). CellRanger generates a peak-barcode counts matrix consisting of the counts of fragment ends within each peak region for each barcode and a fragments file that consists of read fragment ends. The counts matrix and the fragment files generated were utilized for creating Seurat objects via a chromatin assay object.

For each of the objects, quality control was performed separately by a standard pipeline as suggested in the Signac (96) tutorial (https://stuartlab.org/signac/articles/pbmc_vignette), considering peak_region_fragments > 1500 and < 20000, and pct_reads_in_peaks > 15, nucleosome_signal < 4 and TSS.enrichment >2 and were merged using disjoin function from GenomicRanges package. We then performed term frequency-inverse document frequency (TF-IDF) normalization. Later, we carried out singular value decomposition (SVD) on the TD-IDF matrix to obtain the latent semantic indexing (LSI) embeddings on the top features calculated using FindTopFeatures() function. Subsequently, integration was performed on individual mouse replicates using SeuratV3 (85) to acquire integrated LSI reduction. For integration with Seurat, we identified integration anchors between the replicates using FindIntegrationAnchors() function with reduction=‘rlisi’. Finally, we used the IntegrateEmbeddings() function, setting the reduction to LSI embeddings.

#### Multimodal label transfer

We computed a gene activity assay by counting fragments overlapping the gene body and a 2-kb upstream region for each gene in each cell, using the GeneActivity() function in Signac (96). We log-normalized the gene activity counts for the DNA accessibility assay using the NormalizeData() function in Seurat. We then identified anchor cells between the scATAC-seq and scRNA-seq datasets using canonical correlation analysis, with the function FindTransferAnchors() in SeuratV3 with the parameters, reduction=‘cca’. Cell type labels were transferred from the scRNA-seq to scATAC-seq dataset using the TransferData() function, with weight.reduction=query[[‘lsi’]] and dims=2:30 to weight anchors based on nearest-neighbor distances in the LSI space.

#### Finding Differentially Accessible Peaks in Signac

We can perform a differential accessibility (DA) test using the FindMarkers() function with a Wilcoxon rank sum test and min.pct set to 0.1. Peak plots were created from ATAC assay for the accessible chromatin regions with significant p_values (< 0.05) when compared with diet conditions across different time points.

#### Motif enrichment using JASPER and chromVar

A motif class was created in scATAC objects to store information about position weight matrices (PWMs) or position frequency matrices (PFMs) from JASPAR 2020 (97) using AddMotifs() function in Signac (96). RunChromVAR() of chromVAR(98) package in R was used to compute a per-cell motif activity scores. Differentially enriched motifs were identified between cell populations using the FindMarkers() function, only motifs with p_adj_value < 0.05 were considered for further analysis.

### Analysis of single-cell Data from Hs-HIRN Database

Analysis of human islets was performed from the publicly available Human Pancreas Analysis Program (HPAP-RRID: SCR_016202) Database (https://hpap.pmacs.upenn.edu), a Human Islet Research Network (RRID: SCR_014393) consortium (UC4-DK-112217, U01-DK-123594, UC4-DK-112232, and U01-DK-123716) to include Type 2 Diabetic, T1D control and T2D control groups (30). The analysis was performed using SeuratV3 [3]. The data was subset to only include the subjects of interest from both male and female individuals. Individuals with age 22-59 years were included with BMI ranging from 20.8 to 45.49. Individuals with no metabolic disease manifestations including T2D, NAFLD/NASH with BMI < 30 (26.91) were included in the control group (9 individuals). Individuals with BMI >30 (30.85) and HB1Ac ≤ 6 were considered obese (14 individuals) and the individuals designated as “T2D” in the datasets were categorized as T2D (16 individuals).

#### Quality control, dimension-reduction, and clustering

Cells with fewer than 200 or greater than 2500 unique genes, as well as cells with greater than 10% of mitochondrial counts, were excluded from further analysis. Following data filtering, the gene counts matrix was normalized and scaled by using the NormalizeData() and ScaleData() functions, respectively. The top 2000 highest variable genes were used for the principal component analysis (PCA), and the optimal number of PCA components were determined using the ElbowPlot procedure, The resulting PCs 1 to 20 were harmonized across samples using Harmony [8]. Single cells were clustered using the K-nearest neighbor (KNN) graph algorithm in PCA space and visualized using the Uniform Manifold Approximation Projection (UMAP) (91) non-linear dimensionality reduction algorithm over harmony reduction. The FindAllMarkers() function was used to identify novel marker genes for each cluster with min.logfc being set to 0.25 and min.pct being set to 0.20. By using the same parameters, clusters were identified using Louvain clustering (99, 100) based on the harmonized embedding using a resolution of 0.1.

#### Differential expression (DE) analysis

DEGs were determined as genes expressed with an average log (Fold Change) of greater than 0.5 and P_adj_value < 0.05 across all Seurat clusters. DEGs were selected by Seurat FindAllMarkers() based on the Wilcox likelihood-ratio test with default parameters.

For pathway enrichment analysis, we used FGSEA (Fast Gene Set Enrichment Analysis) (101) in the same fashion as in the mouse scRNA-seq analysis, except that pathways were selected from the MSigDB Collections (https://www.gsea-msigdb.org/gsea/msigdb/human/collections.jsp) for the human species.

#### Cell type annotation

The cell type identification of each cluster was manually annotated according to the expression of canonical markers found among the DEGs, combined with data obtained from literature. Violin plots and Dot plots that exhibit the expression of cell-type markers were generated by Seurat VlnPlot() and DotPlot().

#### Sub-clustering of islet cell populations

For in depth analysis of individual cell populations, the populations were isolated from the overall Seurat object and were again subjected to normalization and scaling, followed by default clustering procedure. Sub-clusters were identified using Louvain clustering with an optimal resolution to get clusters with significant differential expression which was validated by single cell level heatmaps using DoHeatmap() of Seurat.

#### Mapping Mm-populations to Hs-clusters: Multimodal reference mapping

Mapping of mouse cells onto human clusters was performed using Seurat multimodal reference mapping (102). For all cells and each subset, the mouse data was prepared by extracting the counts matrix from the mouse single-cell object and mapping the mouse gene names to their human orthologs using a database of ortholog mappings from Mouse Genome Informatics (http://www.informatics.jax.org/homology.shtml). In the case of multi-mapping, the first ortholog pair was used. The mouse object was then split by sample and mapped onto the sNuc-seq data from the matching human all-cell or subset object using the RNA assay and PCA reduction.

#### PheWAS analysis

To evaluate the phenome-wide relationships of the genetic variants associated with the candidate genes across anthropometric and metabolic traits in humans, GWAS summary statistics was examined across 51 variations and 595 phenotypes associated with *CCL27* and *CCR2* genes in the Type 2 Diabetes Knowledge Portal (type2diabetesgenetics.org). *CCL27* Gene page. 2024 Nov 2; https://t2d.hugeamp.org/gene.html?gene=CCL27 (RRID:SCR_003743) and *CCR2* Gene page. 2024 Nov 2; https://t2d.hugeamp.org/gene.html?gene=CCR2 (RRID:SCR_003743) [5]. Briefly, meta-analyzed GWAS summary statistics data was explored across the genomic region harboring candidate genes. The genotyped arrays consisted of individuals across African, American, East Asian, European, and South Asian ancestry. The effect size of the genetic variant was represented by beta values. Association summary statistics were visualized with R forestplot (https://github.com/cran/forestplot) and ggplot2 packages (https://github.com/tidyverse/ggplot2).

#### Comparison between bulk seq data and scRNA seq data

Bulk RNA-seq data (accession number: GSE112002) was retrieved from the Gene Expression Omnibus (GEO) repository. For single-cell RNA-seq analysis, Mm-M1 macrophages from the RC22W and HFD16W samples were subset from a log-normalized Seurat object. Variable genes were identified within these subsets. To compare the transcriptomic profiles of individual Mm-M1 macrophages with the bulk RNA-seq profiles of F4/80 High and CD11c high islet macrophages, Pearson’s correlation coefficients were calculated. This analysis was performed using the 2,000 variable genes common to both the Mm-M1 macrophage subsets and the bulk RNA-seq datasets.

#### Integration and mapping of mouse beta cells with publicly available datasets

To characterize our Mm-Beta 2-1 and Mm-Beta2-2 cell populations, we analyzed two publicly available scRNA sequencing datasets: GSE211799 (the integrated MIA) and GSE137909, focusing on beta cell populations under different metabolic conditions. For the MIA dataset (GSE211799), we extracted beta cells from two conditions: db/db and mSTZ, yielding a total of 46,312 cells. Within the db/db condition, we grouped samples as control_db (chow_WT) and db/db (PF_Lepr-/-, Sham_Lepr-/-, and VSG_Lepr-/-). Similarly, for the mSTZ condition, we grouped samples into control_mSTZ (control) and mSTZ (STZ, STZ_estrogen, STZ_GLP1, STZ_GLP1_estrogen, STZ_GLP1_estrogen + insulin, and STZ_insulin). For the GSE137909 dataset, we preprocessed the raw counts, performed cell type annotation using key marker genes, and extracted beta population under the STZ condition, resulting in 476 beta cells.

To comprehensively analyze beta cell populations, we merged our beta subpopulation with these datasets, yielding 110,187 cells with 15,210 common genes. To integrate these datasets, we used scDREAMER’s (38) unsupervised batch integration method, where CONDITION (db/db, mSTZ, STZ, HFD, RC) was treated as batches. The resulting latent embeddings were used for clustering the cells using Seurat’s "FindNeighbors" and "FindClusters" functions. To visualize the cells, we projected the scDREAMER embeddings into two dimensions using the UMAP algorithm. The resulting clusters were visualized in UMAP space using the "DimPlot" function, grouped by CONDITION and beta subpopulation. To assess the molecular signatures of different beta cell subtypes, we examined the expression of key marker genes across control_db, db/db, control_mSTZ, mSTZ, and Beta2-2 clusters, using “DotPlot” and “VlnPlot” functions. Additionally, we generated Sankey plots using the "plot_sankey_comparison" function from the mapscvi package in R to illustrate mapping between conditions.

#### Gene expression correlation analysis

We used the TFLink (103) to download the transcription factor (TF) list for the Ccl27a gene. We filtered out cells from the Mm-Beta 2 subset having only those cells which show expression for both Ccl27a and the TF. Subsequently, we used the FeatureScatter() function from Seurat to prepare the correlation plots between Ccl27a and each TF. The correlation between Ccl27a and each TF was calculated using the cor.test() function in R (Pearson correlation) to find the R-value and p-value for each TF.

#### Single-cell regulatory network inference and clustering (SCENIC) analysis

To infer transcription factor regulons across Mm-Beta cells, we used SCENIC (55), a regulatory network inference tool and GRNBoost2 algorithm. RNA counts from samples of Mm-Beta2 subset in RC and HFD conditions were exported into a text (.txt) file using the GetAssayData() function from Seurat. The standard pySCENIC (55) workflow was run to find the TFs regulating the expression of Ccl27a gene.

#### Cell-cell communication analysis

For predicting cell-cell interactions between endocrine and non-endocrine cell populations under RC and HFD conditions, the communication networks were inferred using the CellChat R package (version 1.6.1) (51). The Seurat object was formatted into the CellChat format. The data processing and visualization were performed with the default settings of CellChat. The bubble plots were generated by specifying the source and target for the visualization of ligand-receptor pairs. For Mm-Beta 2 cells and macrophages, communication probability was computed using 10% truncated mean (type = “truncatedMean”, trim = 0.1), and other visualizations were performed with default settings. All cell-cell communication analyses were performed between the same timepoints of RC and HFD conditions.

### Data and code Availability

All data were analyzed with standard programs and packages, as detailed in STAR Methods. The code is available at https://github.com/Zafar-Lab/Murine_Pancreatic_Hyperglycemic.

## Figure Legends

**Supp Figure 1. Metabolic profiles of RC and HFD mice at different time points.** (A) Glucose tolerance test (GTT). (B) Insulin tolerance test (ITT). (C) Fed blood glucose, (D) Fed plasma insulin. (E) Fasting blood glycerol. (F) Fed plasma glycerol. (G) Fasting plasma triglycerides. (H) Fed plasma triglycerides. Male mice consumed either regular chow (RC) or a HFD for the indicated time periods. All experiments were performed with male mice after a 6-hour fast except for GTT it was fasted overnight. Data are shown as the mean ± SEM (n = 6 to 9 mice/group). *P < 0.05, ***P < 0.001, and ****P < 0.0001, by 1-way ANOVA followed by Tukey’s post hoc analysis (A-B) and one paired student t-test (C-H)

**Supp Figure 2. Transcriptomic and chromatin accessibility atlas of mouse pancreatic islets.** (A) UMAP visualization of mice sc-RNA seq data for islet types under RC (n= 36518 cells) and HFD (n= 47099 cells) conditions (time course). Cells are colored based on canonical cell types identified through the expression of specific marker genes. (B) Distribution of cells within each cluster assigned to annotated islet cell types under RC and HFD feeding conditions. (C) Distribution of cells within each cluster assigned to islet cell types across different timepoints and dietary conditions. (D) UMAP visualization of mice sc-RNA-seq data for islet types under RC (n= 30706 cells) and HFD (n= 42415 cells) conditions (time course). (E) Percentage distribution of cells within each cluster assigned to annotated islet cell types across different timepoints and dietary conditions. (F) UMAP visualization of islet cells obtained from RC (n= 9844 cells) and HFD (n= 11863 cells) mice, derived from sc-ATAC seq data.

**Supp Figure 3. scRNA-seq analysis of human pancreatic islets.**(A) Graphical representation of the Human Islet Research Network (HIRN) published dataset, including scRNA-seq and scATAC-seq data from control (n=9: scRNA-seq, n=10: scATAC-seq), obese (n=14: scRNA-seq, n=9: scATAC-seq) and T2D (n=16: scRNA-seq, n=5: scATAC-seq) population. (B) UMAP visualization of clusters derived from scRNA-seq data of human islets (n= 43070 cells). Cells are colored based on canonical cell types identified through the expression of specific marker genes. (C) Expression profile of marker genes used for annotating different scRNA-seq islet cell type clusters. (D) Percentage distribution of cells within each cluster assigned to annotated scRNA-seq islet cell types across control, obese, and T2D populations. (**E**) Sankey plots displaying the relationship between mouse and human islet clusters. Mouse cells were mapped onto human scATAC-seq cells using multimodal reference mapping, illustrating the relationship between manually assigned mouse clusters and mapped human clusters for each mouse cell. (F) Combined UMAPs presenting human scRNA-seq pancreatic endocrine cell types (n= 36713 cells). (G) Expression profile of marker genes used for annotating different human islet cell clusters. (H) Percentage distribution of cells within each cluster assigned to annotated human islet cell types across control, obese, and T2D populations. (I) UMAPs presenting human scRNA-seq islet cell types across control, obese, and T2D populations. Cells are colored based on canonical cell types identified through the expression of specific marker genes.

**Supp Figure 4. Temporal transcriptional and epigenomic adaptations in mouse beta cells during HFD feeding.** (A) UMAP embedding of mouse beta cell distribution during RC (n= 30706 cells) and HFD (n= 42415 cells) feeding. Cells are colored by the beta population (Mm-Beta 1, Mm-Beta 2, Mm-Beta 3, Mm-Beta 4 and Mm-Beta 5). (B) Distribution of cells within each cluster assigned to beta cell subtypes across various timepoints. (C) Heatmap for enriched pathways in Mm-Beta 1, 2 and 3 population using VISION. (D) Expression profile of differentially expressed genes in Mm-Beta 2 v/s 3 cell clusters. (E) UMAP embedding depicting Mm-Beta 1 subclusters across RC (n= 15653 cells) and HFD (n= 15639 cells). (F) Percentage of cells within each cluster assigned to annotated Mm-Beta 1 cell subtypes across various timepoints. (G) UMAP embedding showing Mm-Beta 2 subclusters across RC (n= 8275 cells) and HFD (n= 16666 cells) feeding. (H) Percentage of cells within each cluster assigned to annotated Mm-Beta 2 cell subtypes across various timepoints. (I) Heatmap of GO or KEGG pathways enriched in Mm-Beta 2-1 subclusters across different timepoints. (J) Heatmap of GO or KEGG pathways enriched in Mm-Beta 2-2 subclusters across various timepoints. (K) UMAP embedding representing murine Mm-Beta 3 subclusters. (L) Expression profile of differentially expressed genes across Mm-Beta-2 cell subclusters.

**Supp Figure 5. Temporal transcriptional and epigenomic adaptations in mouse Beta 3 cells during HFD feeding.** (A) UMAP embedding representing Mm-Beta-3 (n= 6995 cells) subclusters. Cells are colored by the Mm-Beta-3 population (Mm-Beta 3-1, Mm-Beta 3-2, Mm-Beta 3-3) (B) Expression profile of differentially expressed genes across Mm-Beta 3 cell subclusters. (C) UMAP embedding representing Mm-Beta 3 subclusters across RC (n= 1134 cells) and HFD (n= 5861 cells). (D) Percentage of cells within each cluster assigned to annotated Mm-Beta 3 cell subtypes across various timepoints. (E) GSEA of enriched pathways in murine pancreatic Mm-Beta 3 cell subclusters.

**Supp Figure 6. Comparison of Mm-Beta subpopulation with published data from db/db and STZ treated models.** (A) Joint UMAP embedding from current data, Feng et al., Sachs et al., and Hrovatin et al., (B) Sankey plot representing the correspondence between cluster labels and true cell type labels in current and published data. (C) Set of genes expressed in current data and published data. (D) UMAP visualization of murine scATAC-seq data for Beta cell subtypes across different timepoints. (E) Violin plot of *Ins1&2* expression across RC and HFD conditions.

**Supp Figure 7. Trajectory analysis of Mm-Beta cells using MARGARET:** A) Directed trajectory for Mm-Progenitor, Mm-Beta 1, 2 and 3 cell populations inferred using MARGARET. (B) UMAP visualization of cell-state embeddings inferred by MARGARET, cells are colored by cell identity, (C) MARGARET clusters and (D) MARGARET pseudotime. (E) Expression trend and (F) Normalized expression for *Atf5* gene across Mm-Beta 1, 2 and 3 cell lineages. (G) Expression trend and (H) Normalized expression for *Tshz1* gene across Mm-Beta 1, 2 and 3 cell lineages. (I) Expression trend and (J) Normalized expression for *Fos1* gene across Mm-Beta 1, 2 and 3 cell lineages. (K) Expression trend and (L) Normalized expression for *Jun* gene across Mm-Beta 1, 2 and 3 cell lineages.

**Supp Figure 8. Transcriptional adaptations in human beta cells across control, obese and T2D conditions.** (A) UMAPs of human scRNA-seq beta cell (n= 13506 cells) subclusters across different timepoints. Cells are colored by the Hs-Beta population (Hs-Beta 1, Hs-Beta 2, Hs-Beta 3) (B) Expression profile of differentially expressed genes used for annotating different beta cell type subclusters. (C) GSEA analysis showing enriched pathways in Hs-Beta cell subclusters. (D) Percentage of cells within beta cell subtypes across control, obese and T2D population. (E)Venn diagram of differentially expressed genes in Hs-Beta cells across obese and T2D populations. (F) Correlation analysis between the expression profiles of genes in Hs-Beta 1 across obese and T2D populations. (G) Expression of genes associated with GO and KEGG pathways in pancreatic Hs-Beta-2 subclusters across control, obese, and T2D populations

**Supp Figure 9. Metabolic challenge causes mouse alpha cells to adopt a polyhormonal identity.** (A) UMAP visualization of murine scRNA-seq data for alpha cell subtypes under RC (n= 2615 cells) and HFD (n= 2033 cells) feeding conditions (different timepoints). Cells are colored by subpopulation identities (Mm-Alpha 1 and Mm-Alpha 2). (B) Expression profile of differentially expressed genes used for annotating different alpha cell subclusters. (C) Percentage distribution of cells within each cluster assigned to annotated alpha cell subtypes across different timepoints and dietary conditions. (D) UMAP visualization of murine scATAC-seq data for alpha cell subtypes across different timepoints. (E) Differentially enriched motif profile of Mm-Alpha cell type subclusters. (F) Visualization of normalized expression of *Mafb* (Upper) and *Hesb1* (bottom) transcription factors, respectively on MARGARET inferred trajectory for Mm-Alpha 1 and 2 cells.

**Supp Figure 10. Transcriptional chanbes in human alpha cells across control, obese and T2D conditions.** (A) Combined UMAP plots representing the distribution of human pancreatic alpha cell subtypes (n= 21741 cells). Cells are colored by subpopulation identities (Hs-Alpha 1 and Hs-Alpha 2). (B) Expression profiles of differentially expressed genes used to annotate distinct alpha cell type subclusters. (C) Volcano plot illustrating differentially expressed genes across Hs-Alpha 1 and Hs-Alpha 2 cell types. Red represents upregulated genes in Hs-Alpha 1, and blue represents upregulated genes in Hs-Alpha 2, respectively. (D) Expression patterns of genes associated with GO or KEGG pathways in Hs-Alpha-2 cell clusters across control, obese, and T2D populations. (E) Expression profiles of *GCG*, *INS*, and *SST* in Hs-Alpha 1 and Hs-Alpha 2 cell subclusters. (F) UMAP plots representing the distribution of Hs-Alpha subtypes across different timepoints and in control, obese, and T2D human populations. (G) Percentage distribution of cells within each cluster assigned to annotated Hs-Alpha subtypes across control, obese, and T2D conditions. (H) Results of GSEA hallmark analysis showing enriched gene sets in Hs-Alpha-1 and Hs-Alpha-2 populations.

**Supp Figure 11. Transcriptional heterogeneity of murine delta cells at the transcriptomic and epigenomic levels**. (A-B) UMAP embedding of Mm-Delta cell subclusters across different timepoints and dietary conditions (n= 3008 cells). Cells are colored by subpopulation identities (Mm-Delta 1, Mm-Delta 2 and Mm-Delta 3). (C) Differentially enriched motifs in scATAC-seq Mm-Delta 1 and Mm-Delta 2 populations. (D) Enriched motifs in scATAC-seq of Mm-Delta 1 cells. (E) Visualization of directed pseudotime trajectory for Mm-Progenitor, Mm-Delta 1, 2 and 3 cell populations inferred by MARGARET. (F) UMAP projection of clusters formed by scRNA-seq Hs-Delta cells. Cells are colored by subpopulation identities (Hs-Delta 1 and Hs-Delta 2). (G) Expression profile of differentially expressed genes used for annotating different Hs-Delta cell type clusters. (H) Volcano plots for differentially expressed genes in human scRNA-seq delta cells, with red indicating upregulated genes in Hs-Delta 1 and blue indicating upregulated genes in Hs-Delta 2, respectively. (I) GSEA analysis showing enriched pathways in Hs-Delta 1 and Hs-Delta 2 populations (J) UMAPs displaying human scRNA-seq delta cell subtypes across control, obese, and type 2 diabetic populations.

**Supp Figure 12. Transcriptional heterogeneity of murine gamma cells at the transcriptomic levels**. (A) UMAP visualization of mice scRNA-seq data for gamma cell types across RC (n= 1036), HFD (n= 630 cells) and timepoints. (B) UMAP visualization of scRNA-seq data for Mm-Gamma cell types across different timepoints and dietary conditions. Cells are colored by subpopulation identities (Mm-Gamma 1 and Mm-Gamma 2). (C) Percentage distribution of cells within each cluster assigned to annotated Mm-Gamma cell types across different timepoints and dietary conditions. (D) GSEA analysis showing enriched pathways in Mm-Gamma 1 and Mm-Gamma 2 populations. (E) Heatmap of GO and KEGG pathways in Mm-Gamma 1 subclusters across different timepoints and dietary conditions.

**Supp Figure 13. Heterogeneity of murine islet endothelial cells through transcriptomic profiling.** (A) UMAP visualization of murine islet endothelial cell subtypes across RC (n= 2505 cells), HFD (n= 1520 cells) and timepoints. Cells are colored by subpopulation identities (Mm-Vascular ECs, Mm-Lipid handling ECs, Mm-Progenitor ECs, and Mm-Pericytes). (B) Percentage of cells within each cluster assigned to annotated islet endothelial cell across different dietary conditions and timepoints. (D) Expression of genes associated with pathways like cytokine production and response associated genes in murine islet vascular endothelial cells.

**Supp Figure 14. Heterogeneity in human endothelial cells identified through transcriptomic profiling.** (A) UMAP visualization of human islet endothelial cell subtypes (n= 4128 cells). Cells are colored by subpopulation identities (Hs-Vascular ECs, Hs-Metabolically active ECs, Hs-Proinflammatory ECs, and Hs-Immune regulatory ECs). (B) Expression profile of genes identifying islet endothelial cell subclusters. (C) UMAP embedding of human endothelial cell subclusters across health states. (D) Percentage of cells within each cluster assigned to annotated islet endothelial cell types across control, obese and T2D populations. (E) GSEA analysis showing enriched pathways in islet endothelial cells: top left - Hs vascular endothelial ECs; bottom Left - Hs immune regulatory endothelial ECs; top right - Hs metabolically active endothelial ECs; bottom right – Hs Proinflammatory endothelial ECs. (F) Expression of genes associated with pathway (Vascular development/angiogenesis, Programmed cell death, Peptidase activity) in Hs-Vascular ECs.

**Supp Figure 15. Heterogeneity of murine islet immune cells through transcriptomic profiling.** (A) UMAP visualization of murine islet immune cell subtypes across RC (n= 1408 cells), HFD (n= 899 cells) and timepoints. Cells are colored by immune cell types marker. (B) Percentage of cells within each cluster assigned to annotated islet immune cell subtypes across across different dietary conditions and timepoints. (C) Comparison of identified islet Mm-M1 macrophages with intra-CD11+ macrophages, as defined by Ying et al. (2019).

**Supp Figure 16. Heterogeneity of human islet immune cells through transcriptomic profiling.** (A) Combined UMAP visualization of human islet immune cell subtypes (n= 1667 cells). Cells are colored by immune cell types marker. (B) Expression profile of genes identifying islet immune cell subclusters. (C) UMAP visualization of islet immune cell subtypes across health states. (D) Expression of pathway (Leukocyte chemotaxis, Inflammatory response via *NFkb* signaling, Antigen presentation and processing and Translation) associated genes in Hs-Macrophages cells.

**Supp Figure 17. Endocrine-endocrine cells interactions in murine pancreatic islets under different metabolic conditions.** (A) Comparison of incoming (B) Outgoing (C) Overall information flow for the top signaling pathways inferred for communication within the endocrine cells across RC (left) and HFD (right) conditions.

**Supp Figure 18. Non endocrine-endocrine cells interactions in murine pancreatic islets under different metabolic conditions**.(A) 2D projection of incoming/outgoing interaction strength among endocrine-non endocrine cell types under RC and HFD conditions. (B) Comparison of overall signaling pattern between endocrine and non-endocrine cells across RC (left) and HFD (right) conditions. (C) Heatmap representing MHC II (top); IL1 (middle); and periostin (bottom) signaling from endocrine cells as source (sender) to non-endocrine cells under RC conditions. (E) Dotplot representing signaling axis between Mm-Beta 2 and Mm-M1 macrophages

**Supp Figure 19. CCL27 and Insulin colocalization across all RC and HFD timepoints** (A) RC (B) HFD Immunofluorescence staining of pancreatic slices. Slices were co-stained with either an anti-CCL27 antibody (Alexa Fluor, green) and an anti-insulin antibody (Alexa Fluor, red). The inserts in the upper right of each panel show enlarged islets areas. Scale bar in inserts: 51 μm: Contrast was adjusted for improved visualization.

## Key Reagents and Resources

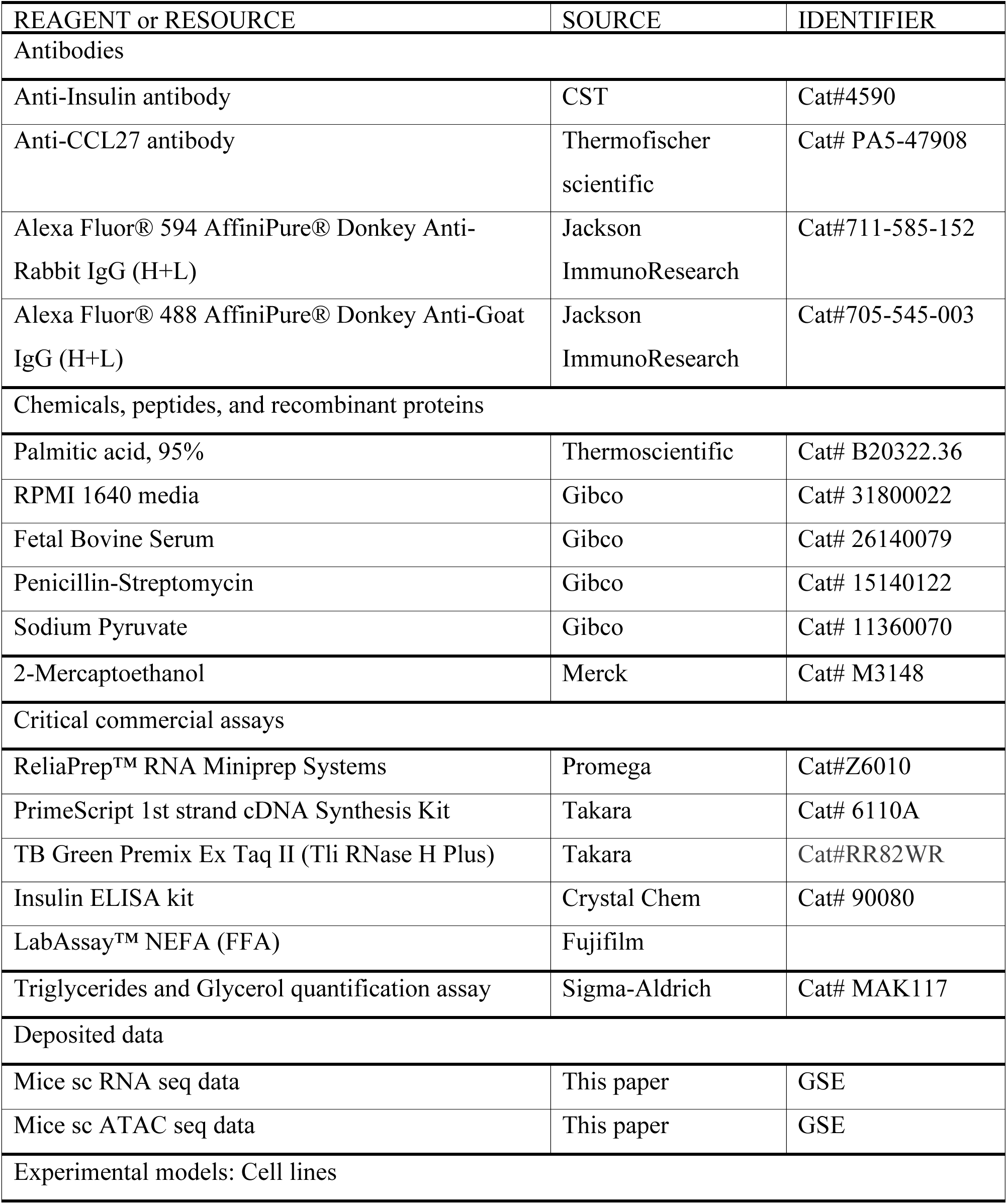
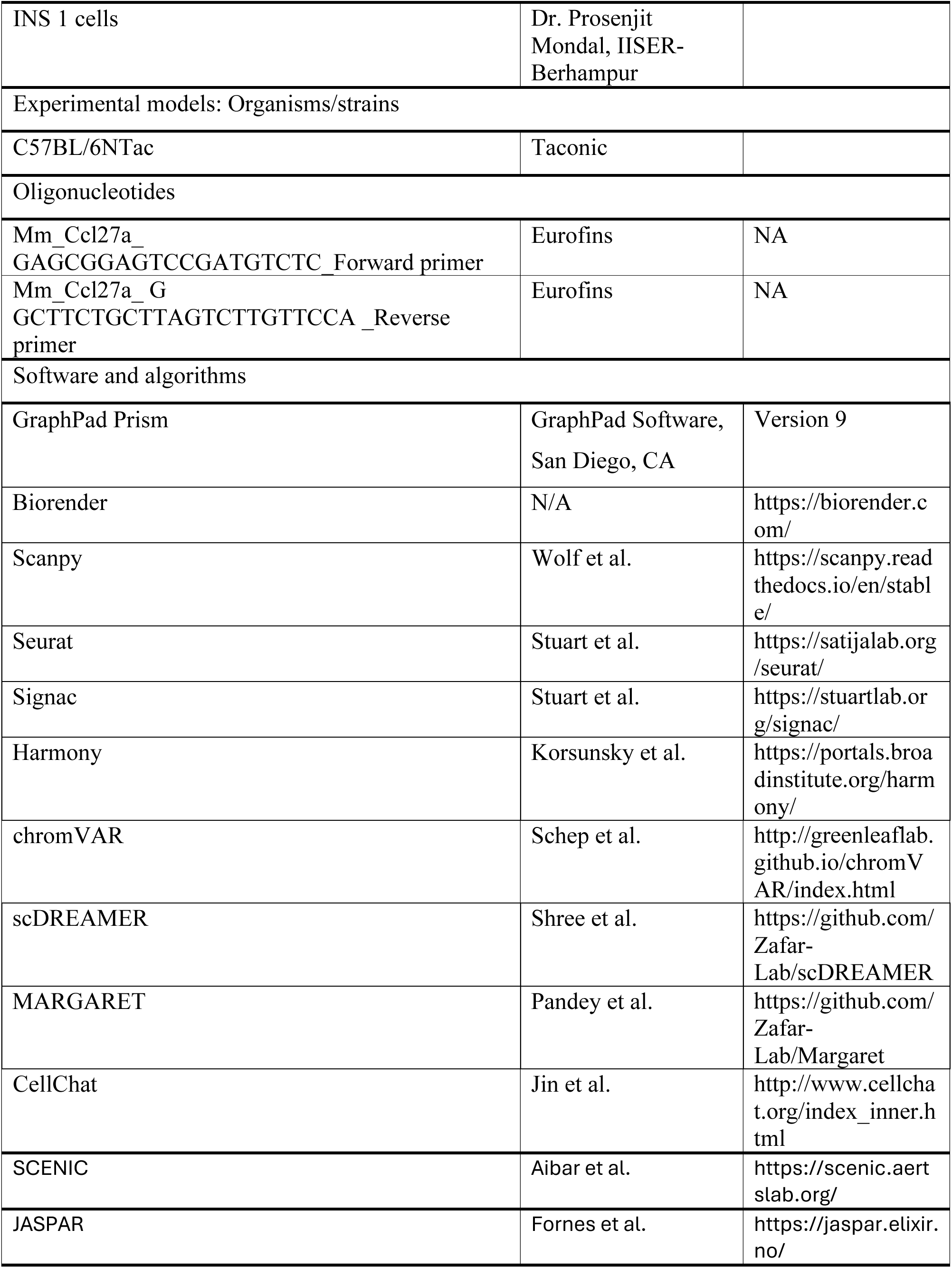
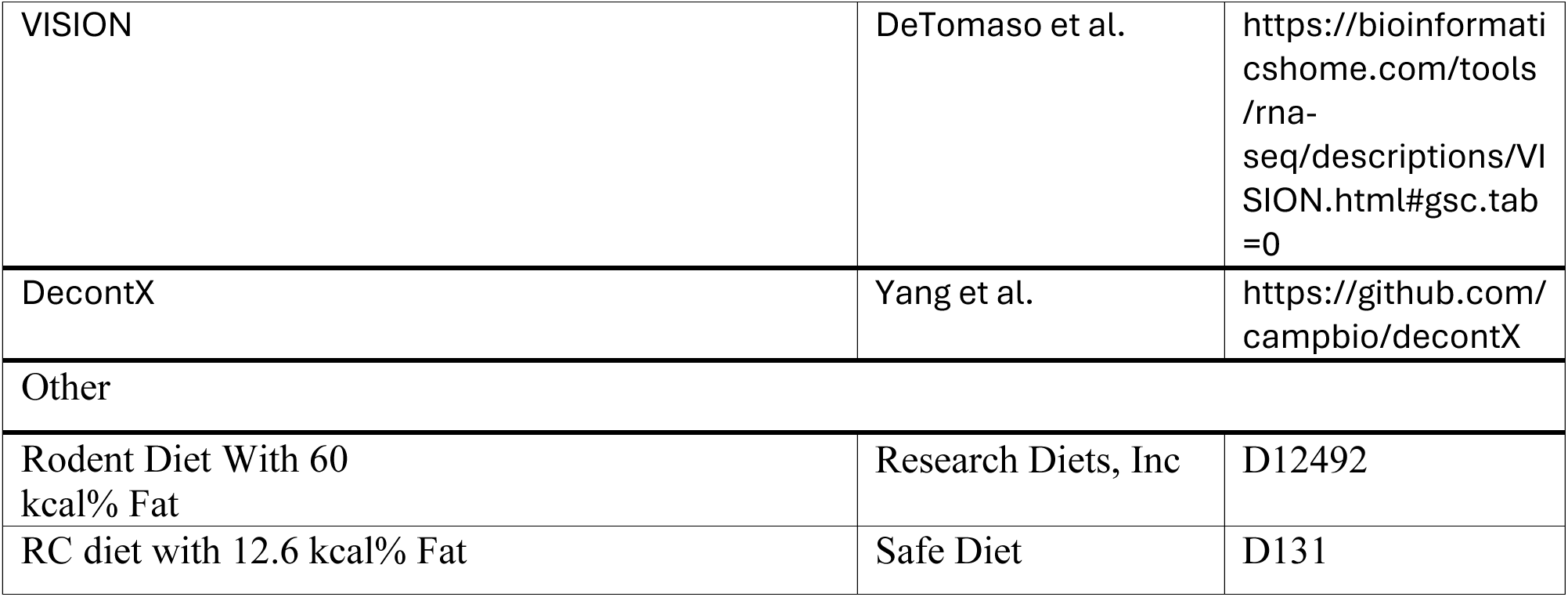

## Notes

### Competing Interest Statement

The authors have declared no competing interest.

