## Supplementary Figures for "A Temporal Single-Cell Multi-Omics Atlas of Murine Pancreatic Islet Remodeling During Hyperglycaemia Progression"

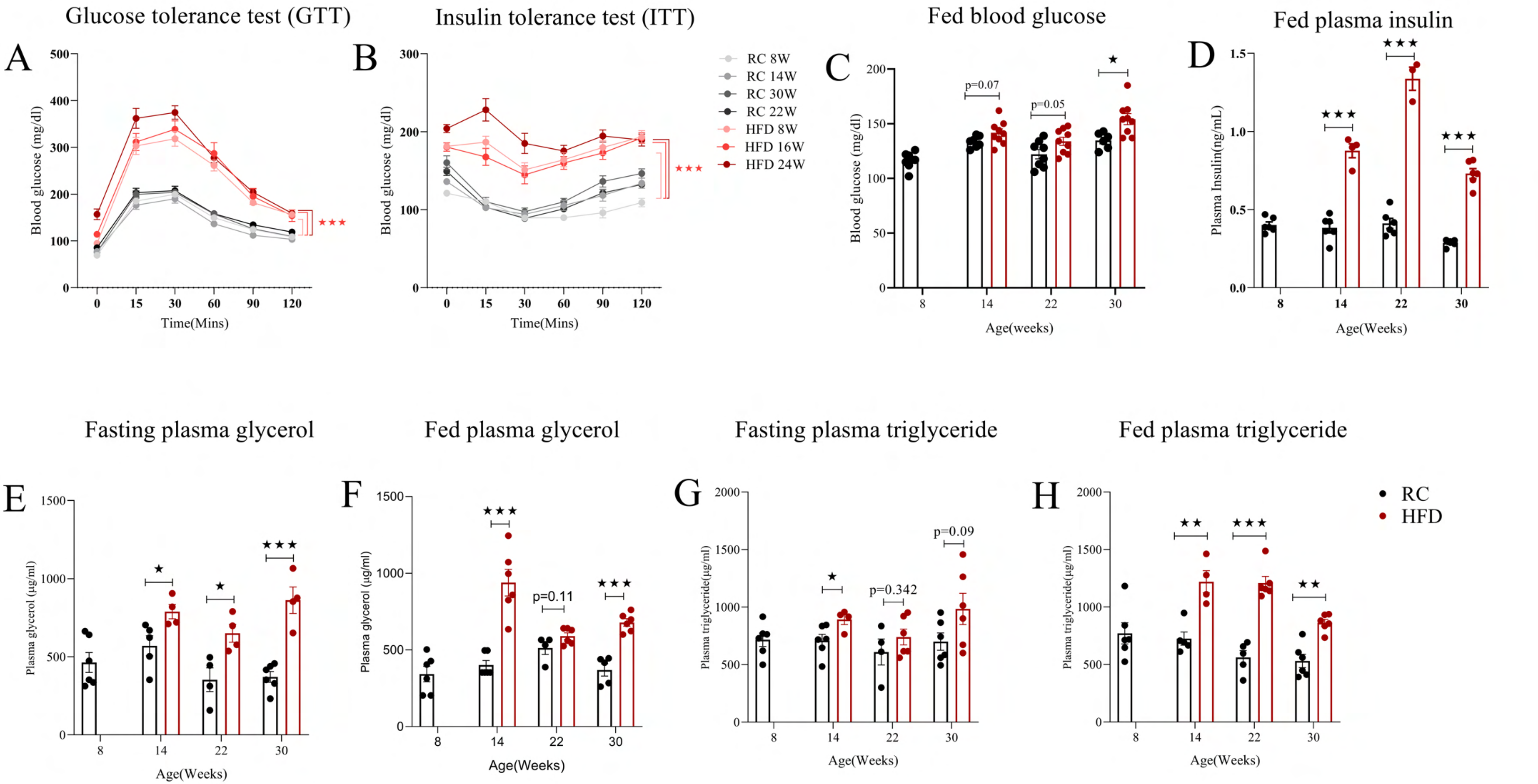

Glucose tolerance test (GTT)
A
Blood glucose (mg/dl)
Time(Mins)
Insulin tolerance test (ITT)
B
Blood glucose (mg/dl)
Time(Mins)
RC 8W
RC 14W
RC 30W
RC 22W
HFD 8W
HFD 16W
HFD 24W
Fed blood glucose
C
Blood glucose (mg/dl)
p=0.07
p=0.05
Age(weeks)
Fed plasma insulin
D
Plasma Insulin(ng/mL)
Age(Weeks)
Fasting plasma glycerol
E
Plasma glycerol (μg/ml)
Age(Weeks)
Fed plasma glycerol
F
Plasma glycerol (μg/ml)
p=0.11
Age(Weeks)
Fasting plasma triglyceride
G
Plasma triglyceride(μg/ml)
p=0.342
p=0.09
Age(Weeks)
Fed plasma triglyceride
H
Plasma triglyceride(μg/ml)
Age(Weeks)
RC
HFD

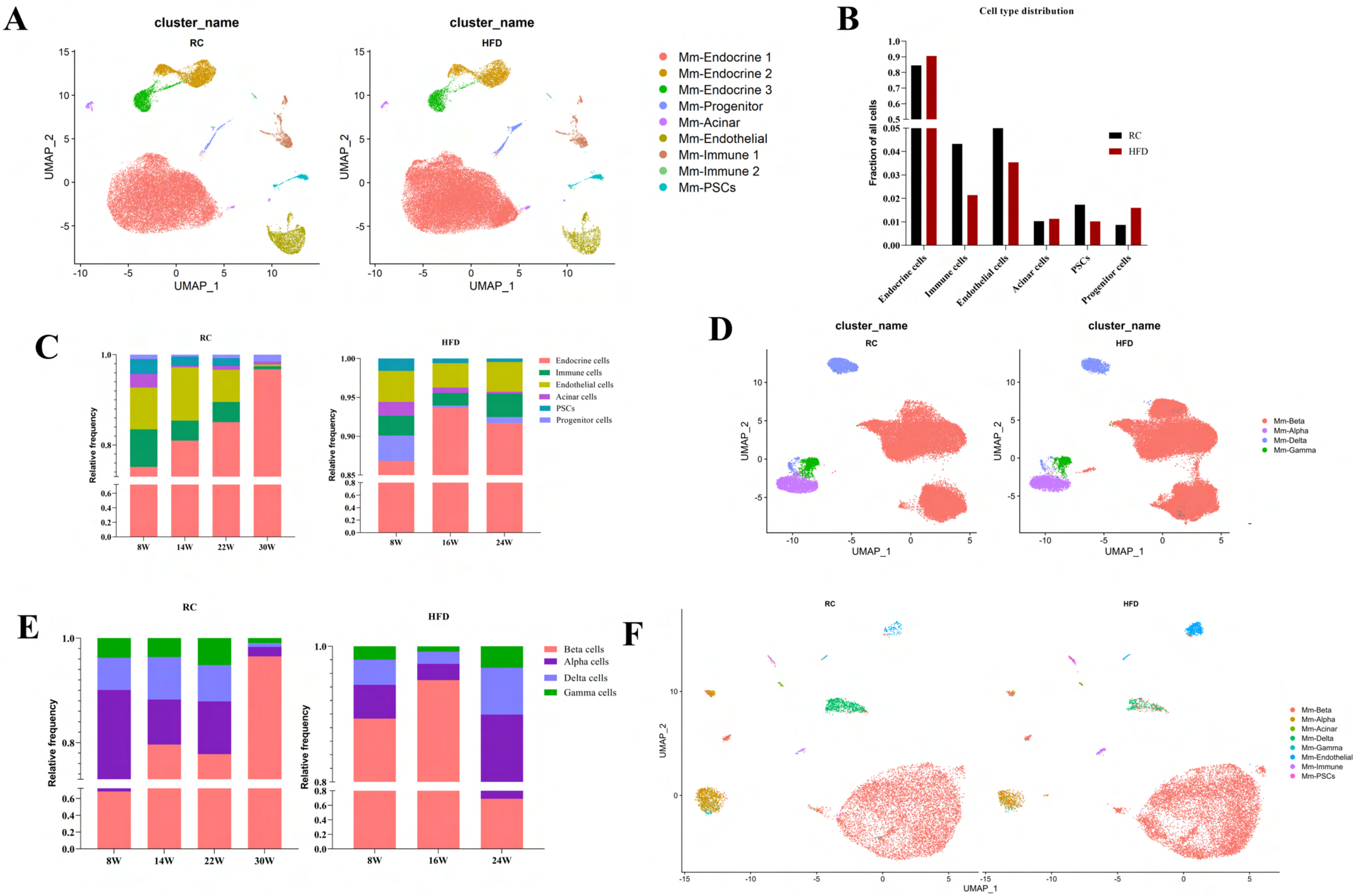

A
cluster_name
RC
HFD
UMAP_1
UMAP_2
Mm-Endocrine 1
Mm-Endocrine 2
Mm-Endocrine 3
Mm-Progenitor
Mm-Acinar
Mm-Endothelial
Mm-Immune 1
Mm-Immune 2
Mm-PSCs
B
Cell type distribution
Fraction of all cells
RC
HFD
Endocrine cells
Immune cells
Endothelial cells
Acinar cells
PSCs
Progenitor cells
C
RC
HFD
Relative frequency
Endocrine cells
Immune cells
Endothelial cells
Acinar cells
PSCs
Progenitor cells
8W
14W
22W
30W
16W
24W
D
cluster_name
RC
HFD
UMAP_1
UMAP_2
Mm-Beta
Mm-Alpha
Mm-Delta
Mm-Gamma
E
RC
HFD
Relative frequency
Beta cells
Alpha cells
Delta cells
Gamma cells
8W
14W
22W
30W
16W
24W
F
RC
HFD
UMAP_1
UMAP_2
Mm-Beta
Mm-Alpha
Mm-Acinar
Mm-Delta
Mm-Gamma
Mm-Endothelial
Mm-Immune
Mm-PSCs

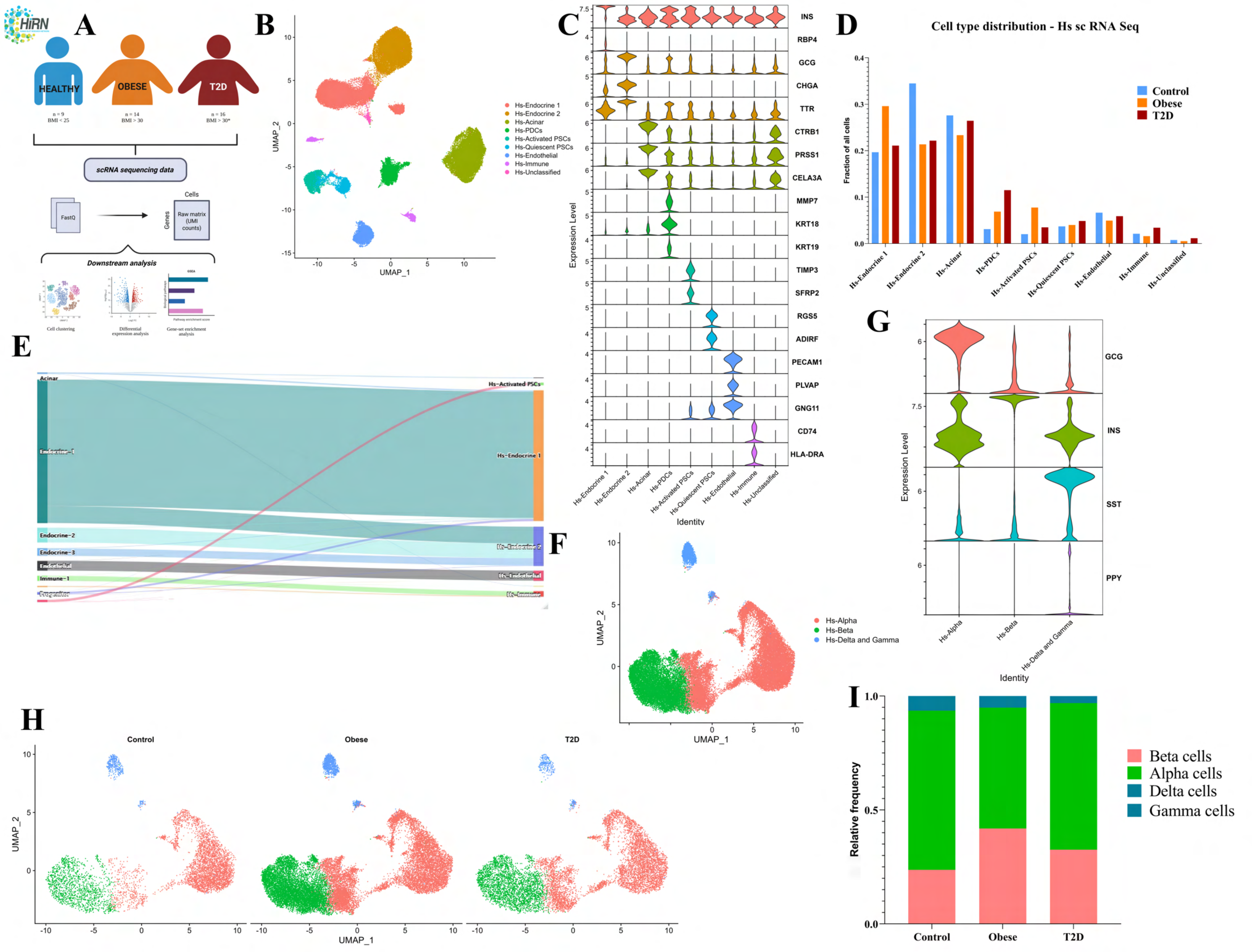
A
HiRN
HEALTHY
OBESE
T2D
n = 9
BMI < 25
n = 14
BMI > 30
n = 16
BMI > 30*
scRNA sequencing data
FastQ
Cells
Genes
Raw matrix (UMI counts)
Downstream analysis
Cell clustering
Differential expression analysis
Gene-set enrichment analysis
B
UMAP_1
UMAP_2
Hs-Endocrine 1
Hs-Endocrine 2
Hs-Acinar
Hs-PDCs
Hs-Activated PSCs
Hs-Quiescent PSCs
Hs-Endothelial
Hs-Immune
Hs-Unclassified
C
INS
RBP4
GCG
CHGA
TTR
CTRB1
PRSS1
CELA3A
MMP7
KRT18
KRT19
TIMP3
SFRP2
RGS5
ADIRF
PECAM1
PLVAP
GNG11
CD74
HLA-DRA
Expression Level
Identity
D
Cell type distribution - Hs sc RNA Seq
Fraction of all cells
Control
Obese
T2D
E
Acinar
Endocrine-1
Endocrine-2
Endocrine-3
Endothelial
Immune-1
Progenitor
Hs-Activated PSCs
Hs-Endocrine 1
Hs-Endocrine 2
Hs-Endothelial
Hs-Immune
F
Hs-Alpha
Hs-Beta
Hs-Delta and Gamma
UMAP_1
UMAP_2
G
GCG
INS
SST
PPY
Expression Level
Identity
H
Control
Obese
T2D
UMAP_1
UMAP_2
I
Relative frequency
Beta cells
Alpha cells
Delta cells
Gamma cells
Control
Obese
T2D

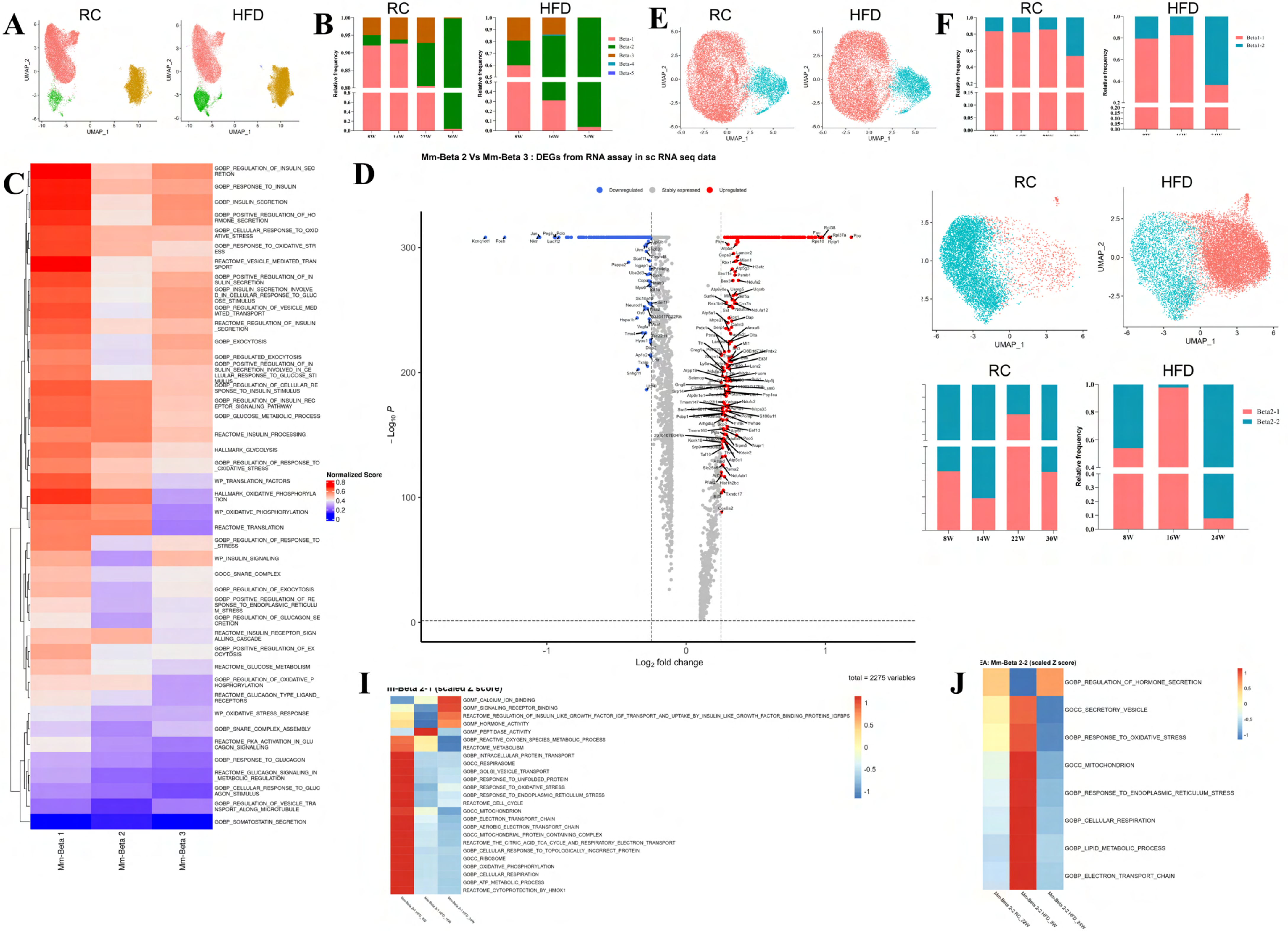
A
RC
HFD
UMAP_1
UMAP_2
B
RC
HFD
Relative frequency
Beta-1
Beta-2
Beta-3
Beta-4
Beta-5
E
RC
HFD
UMAP_1
UMAP_2
F
RC
HFD
Relative frequency
Beta1-1
Beta1-2
C
GOBP_REGULATION_OF_INSULIN_SECRETION
GOBP_RESPONSE_TO_INSULIN
GOBP_INSULIN_SECRETION
GOBP_POSITIVE_REGULATION_OF_HORMONE_SECRETION
GOBP_CELLULAR_RESPONSE_TO_OXIDATIVE_STRESS
GOBP_RESPONSE_TO_OXIDATIVE_STRESS
REACTOME_VESICLE_MEDIATED_TRANSPORT
GOBP_POSITIVE_REGULATION_OF_INSULIN_SECRETION
GOBP_INSULIN_SECRETION_INVOLVED_IN_CELLULAR_RESPONSE_TO_GLUCOSE_STIMULUS
GOBP_REGULATION_OF_VESICLE_MEDIATED_TRANSPORT
REACTOME_REGULATION_OF_INSULIN_SECRETION
GOBP_EXOCYTOSIS
GOBP_REGULATED_EXOCYTOSIS
GOBP_POSITIVE_REGULATION_OF_INSULIN_SECRETION_INVOLVED_IN_CELLULAR_RESPONSE_TO_GLUCOSE_STIMULUS
GOBP_REGULATION_OF_CELLULAR_RESPONSE_TO_INSULIN_STIMULUS
GOBP_REGULATION_OF_INSULIN_RECEPTOR_SIGNALING_PATHWAY
GOBP_GLUCOSE_METABOLIC_PROCESS
REACTOME_INSULIN_PROCESSING
HALLMARK_GLYCOLYSIS
GOBP_REGULATION_OF_RESPONSE_TO_OXIDATIVE_STRESS
WP_TRANSLATION_FACTORS
HALLMARK_OXIDATIVE_PHOSPHORYLATION
WP_OXIDATIVE_PHOSPHORYLATION
REACTOME_TRANSLATION
GOBP_REGULATION_OF_RESPONSE_TO_STRESS
WP_INSULIN_SIGNALING
GOCC_SNARE_COMPLEX
GOBP_REGULATION_OF_EXOCYTOSIS
GOBP_POSITIVE_REGULATION_OF_RESPONSE_TO_ENDOPLASMIC_RETICULUM_STRESS
GOBP_REGULATION_OF_GLUCAGON_SECRETION
REACTOME_INSULIN_RECEPTOR_SIGNALLING_CASCADE
GOBP_POSITIVE_REGULATION_OF_EXOCYTOSIS
REACTOME_GLUCOSE_METABOLISM
GOBP_REGULATION_OF_OXIDATIVE_PHOSPHORYLATION
REACTOME_GLUCAGON_TYPE_LIGAND_RECEPTORS
WP_OXIDATIVE_STRESS_RESPONSE
GOBP_SNARE_COMPLEX_ASSEMBLY
REACTOME_PKA_ACTIVATION_IN_GLUCAGON_SIGNALLING
GOBP_RESPONSE_TO_GLUCAGON
REACTOME_GLUCAGON_SIGNALING_IN_METABOLIC_REGULATION
GOBP_CELLULAR_RESPONSE_TO_GLUCAGON_STIMULUS
GOBP_REGULATION_OF_VESICLE_TRANSPORT_ALONG_MICROTUBULE
GOBP_SOMATOSTATIN_SECRETION
Normalized Score
Mm-Beta 1
Mm-Beta 2
Mm-Beta 3
Mm-Beta 2 Vs Mm-Beta 3 : DEGs from RNA assay in sc RNA seq data
D
Downregulated
Stably expressed
Upregulated
– Log10 P
Log2 fold change
total = 2275 variables
RC
HFD
UMAP_1
UMAP_2
RC
HFD
Relative frequency
Beta2-1
Beta2-2
8W
14W
22W
16W
24W
I
Mm-Beta 2-1 (scaled Z score)
GOMF_CALCIUM_ION_BINDING
GOMF_SIGNALING_RECEPTOR_BINDING
REACTOME_REGULATION_OF_INSULIN_LIKE_GROWTH_FACTOR_IGF_TRANSPORT_AND_UPTAKE_BY_INSULIN_LIKE_GROWTH_FACTOR_BINDING_PROTEINS_IGFBPS
GOMF_HORMONE_ACTIVITY
GOMF_PEPTIDASE_ACTIVITY
GOBP_REACTIVE_OXYGEN_SPECIES_METABOLIC_PROCESS
REACTOME_METABOLISM
GOBP_INTRACELLULAR_PROTEIN_TRANSPORT
GOCC_RESPIRASOME
GOBP_GOLGI_VESICLE_TRANSPORT
GOBP_RESPONSE_TO_UNFOLDED_PROTEIN
GOBP_RESPONSE_TO_OXIDATIVE_STRESS
GOBP_RESPONSE_TO_ENDOPLASMIC_RETICULUM_STRESS
REACTOME_CELL_CYCLE
GOCC_MITOCHONDRION
GOBP_ELECTRON_TRANSPORT_CHAIN
GOBP_AEROBIC_ELECTRON_TRANSPORT_CHAIN
GOCC_MITOCHONDRIAL_PROTEIN_CONTAINING_COMPLEX
REACTOME_THE_CITRIC_ACID_TCA_CYCLE_AND_RESPIRATORY_ELECTRON_TRANSPORT
GOBP_CELLULAR_RESPONSE_TO_TOPOLOGICALLY_INCORRECT_PROTEIN
GOCC_RIBOSOME
GOBP_OXIDATIVE_PHOSPHORYLATION
GOBP_CELLULAR_RESPIRATION
GOBP_ATP_METABOLIC_PROCESS
REACTOME_CYTOPROTECTION_BY_HMOX1
J
Mm-Beta 2-2 (scaled Z score)
GOBP_REGULATION_OF_HORMONE_SECRETION
GOCC_SECRETORY_VESICLE
GOBP_RESPONSE_TO_OXIDATIVE_STRESS
GOCC_MITOCHONDRION
GOBP_RESPONSE_TO_ENDOPLASMIC_RETICULUM_STRESS
GOBP_CELLULAR_RESPIRATION
GOBP_LIPID_METABOLIC_PROCESS
GOBP_ELECTRON_TRANSPORT_CHAIN
Mm-Beta 2-2 RC_22W
Mm-Beta 2-2 HFD_8W
Mm-Beta 2-2 HFD_24W

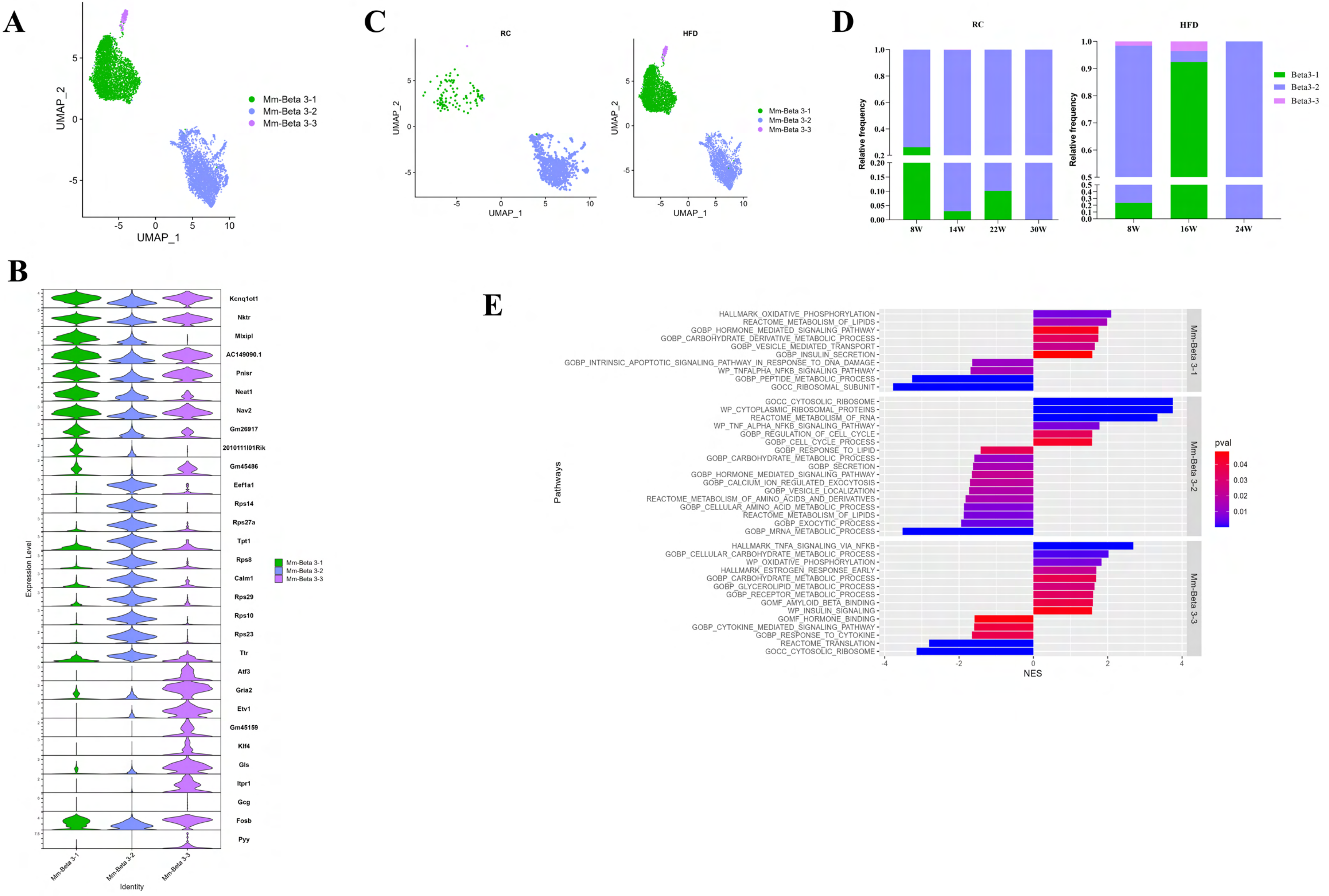
A
UMAP_1
UMAP_2
Mm-Beta 3-1
Mm-Beta 3-2
Mm-Beta 3-3
C
RC
HFD
UMAP_1
UMAP_2
Mm-Beta 3-1
Mm-Beta 3-2
Mm-Beta 3-3
D
RC
HFD
Relative frequency
8W
14W
22W
30W
8W
16W
24W
Beta3-1
Beta3-2
Beta3-3
B
Kcnq1ot1
Nktr
Mlxipl
AC149090.1
Pnisr
Neat1
Nav2
Gm26917
2010111I01Rik
Gm45486
Eef1a1
Rps14
Rps27a
Tpt1
Rps8
Calm1
Rps29
Rps10
Rps23
Ttr
Atf3
Gria2
Etv1
Gm45159
Klf4
Gls
Itpr1
Gcg
Fosb
Pyy
Expression Level
Identity
Mm-Beta 3-1
Mm-Beta 3-2
Mm-Beta 3-3
E
Pathways
HALLMARK_OXIDATIVE_PHOSPHORYLATION
REACTOME_METABOLISM_OF_LIPIDS
GOBP_HORMONE_MEDIATED_SIGNALING_PATHWAY
GOBP_CARBOHYDRATE_DERIVATIVE_METABOLIC_PROCESS
GOBP_VESICLE_MEDIATED_TRANSPORT
GOBP_INSULIN_SECRETION
GOBP_INTRINSIC_APOPTOTIC_SIGNALING_PATHWAY_IN_RESPONSE_TO_DNA_DAMAGE
WP_TNFALPHA_NFKB_SIGNALING_PATHWAY
GOBP_PEPTIDE_METABOLIC_PROCESS
GOCC_RIBOSOMAL_SUBUNIT
GOCC_CYTOSOLIC_RIBOSOME
WP_CYTOPLASMIC_RIBOSOMAL_PROTEINS
REACTOME_METABOLISM_OF_RNA
WP_TNF_ALPHA_NFKB_SIGNALING_PATHWAY
GOBP_REGULATION_OF_CELL_CYCLE
GOBP_CELL_CYCLE_PROCESS
GOBP_RESPONSE_TO_LIPID
GOBP_CARBOHYDRATE_METABOLIC_PROCESS
GOBP_SECRETION
GOBP_HORMONE_MEDIATED_SIGNALING_PATHWAY
GOBP_CALCIUM_ION_REGULATED_EXOCYTOSIS
GOBP_VESICLE_LOCALIZATION
REACTOME_METABOLISM_OF_AMINO_ACIDS_AND_DERIVATIVES
GOBP_CELLULAR_AMINO_ACID_METABOLIC_PROCESS
REACTOME_METABOLISM_OF_LIPIDS
GOBP_EXOCYTIC_PROCESS
GOBP_MRNA_METABOLIC_PROCESS
HALLMARK_TNFA_SIGNALING_VIA_NFKB
GOBP_CELLULAR_CARBOHYDRATE_METABOLIC_PROCESS
WP_OXIDATIVE_PHOSPHORYLATION
HALLMARK_ESTROGEN_RESPONSE_EARLY
GOBP_CARBOHYDRATE_METABOLIC_PROCESS
GOBP_GLYCEROLIPID_METABOLIC_PROCESS
GOBP_RECEPTOR_METABOLIC_PROCESS
GOMF_AMYLOID_BETA_BINDING
WP_INSULIN_SIGNALING
GOMF_HORMONE_BINDING
GOBP_CYTOKINE_MEDIATED_SIGNALING_PATHWAY
GOBP_RESPONSE_TO_CYTOKINE
REACTOME_TRANSLATION
GOCC_CYTOSOLIC_RIBOSOME
Mm-Beta 3-1
Mm-Beta 3-2
Mm-Beta 3-3
NES
pval
0.04
0.03
0.02
0.01

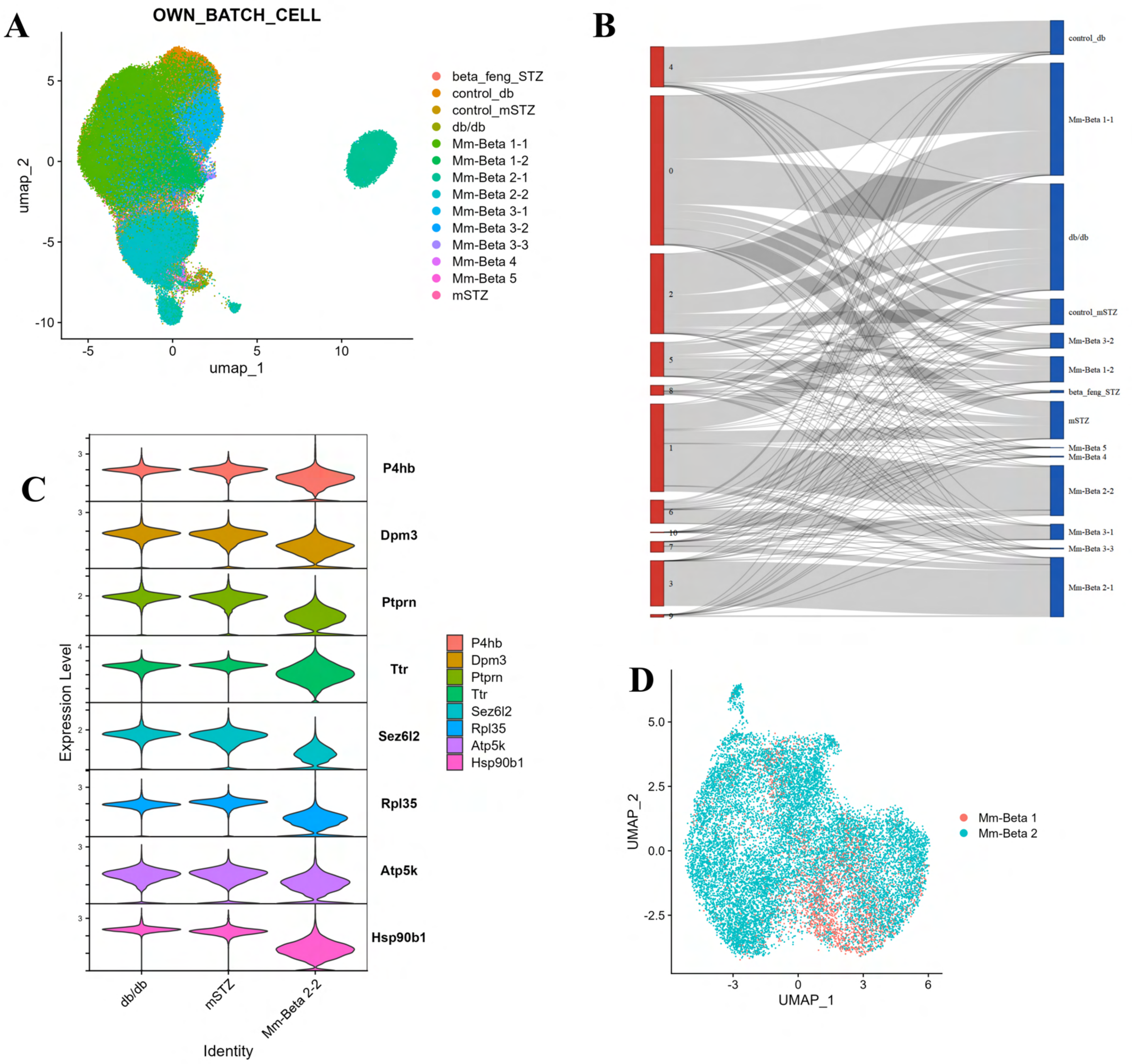
A
OWN_BATCH_CELL
umap_2
umap_1
beta_feng_STZ
control_db
control_mSTZ
db/db
Mm-Beta 1-1
Mm-Beta 1-2
Mm-Beta 2-1
Mm-Beta 2-2
Mm-Beta 3-1
Mm-Beta 3-2
Mm-Beta 3-3
Mm-Beta 4
Mm-Beta 5
mSTZ
B
control_db
Mm-Beta 1-1
db/db
control_mSTZ
Mm-Beta 3-2
Mm-Beta 1-2
beta_feng_STZ
mSTZ
Mm-Beta 5
Mm-Beta 4
Mm-Beta 2-2
Mm-Beta 3-1
Mm-Beta 3-3
Mm-Beta 2-1
C
Expression Level
P4hb
Dpm3
Ptprn
Ttr
Sez6l2
Rpl35
Atp5k
Hsp90b1
db/db
mSTZ
Mm-Beta 2-2
Identity
D
UMAP_2
UMAP_1
Mm-Beta 1
Mm-Beta 2

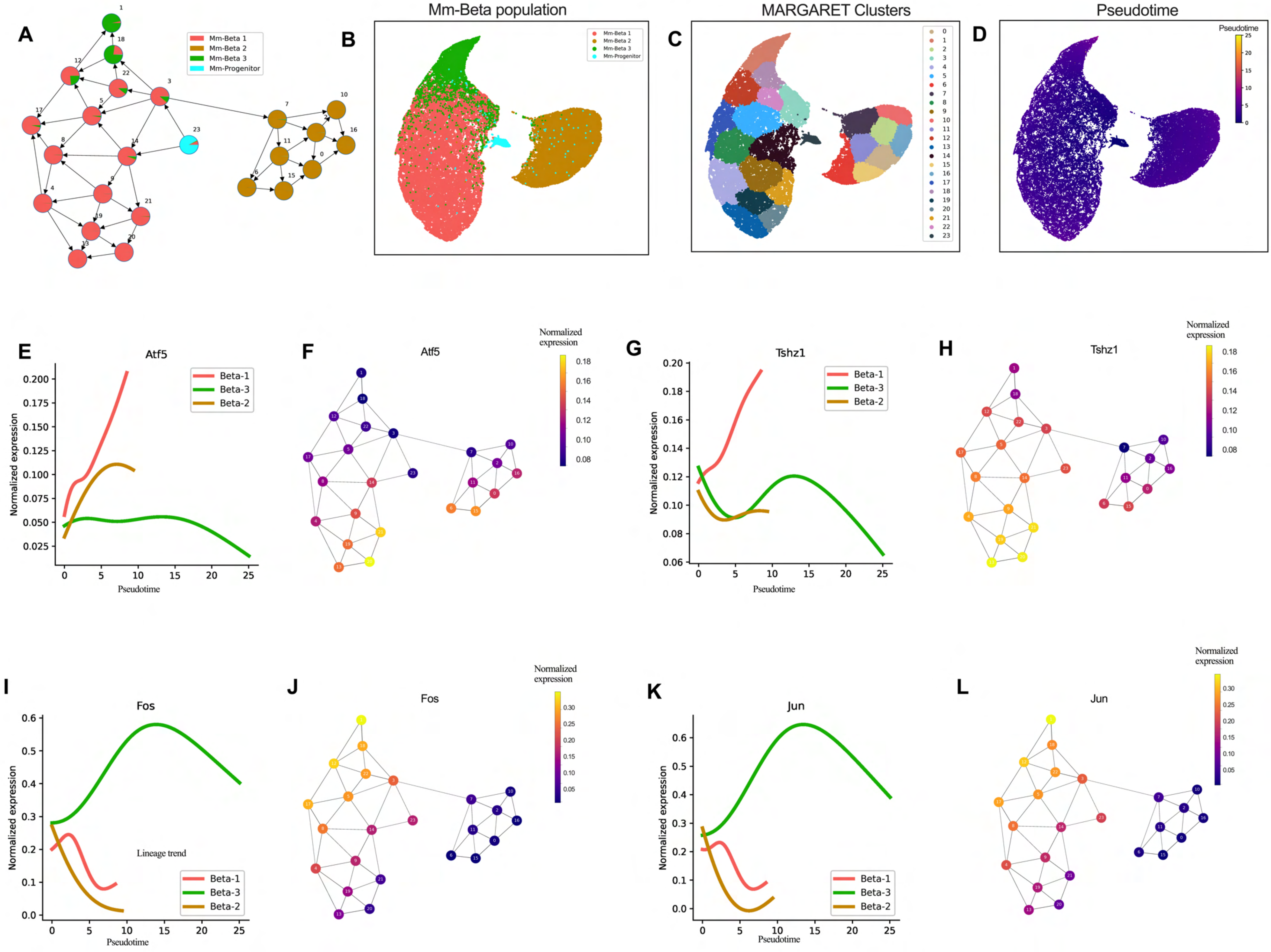

A
Mm-Beta 1
Mm-Beta 2
Mm-Beta 3
Mm-Progenitor
B
Mm-Beta population
C
MARGARET Clusters
D
Pseudotime
E
Atf5
Normalized expression
Pseudotime
Beta-1
Beta-3
Beta-2
F
Atf5
Normalized expression
G
Tshz1
H
Tshz1
I
Fos
Lineage trend
J
Fos
K
Jun
L
Jun

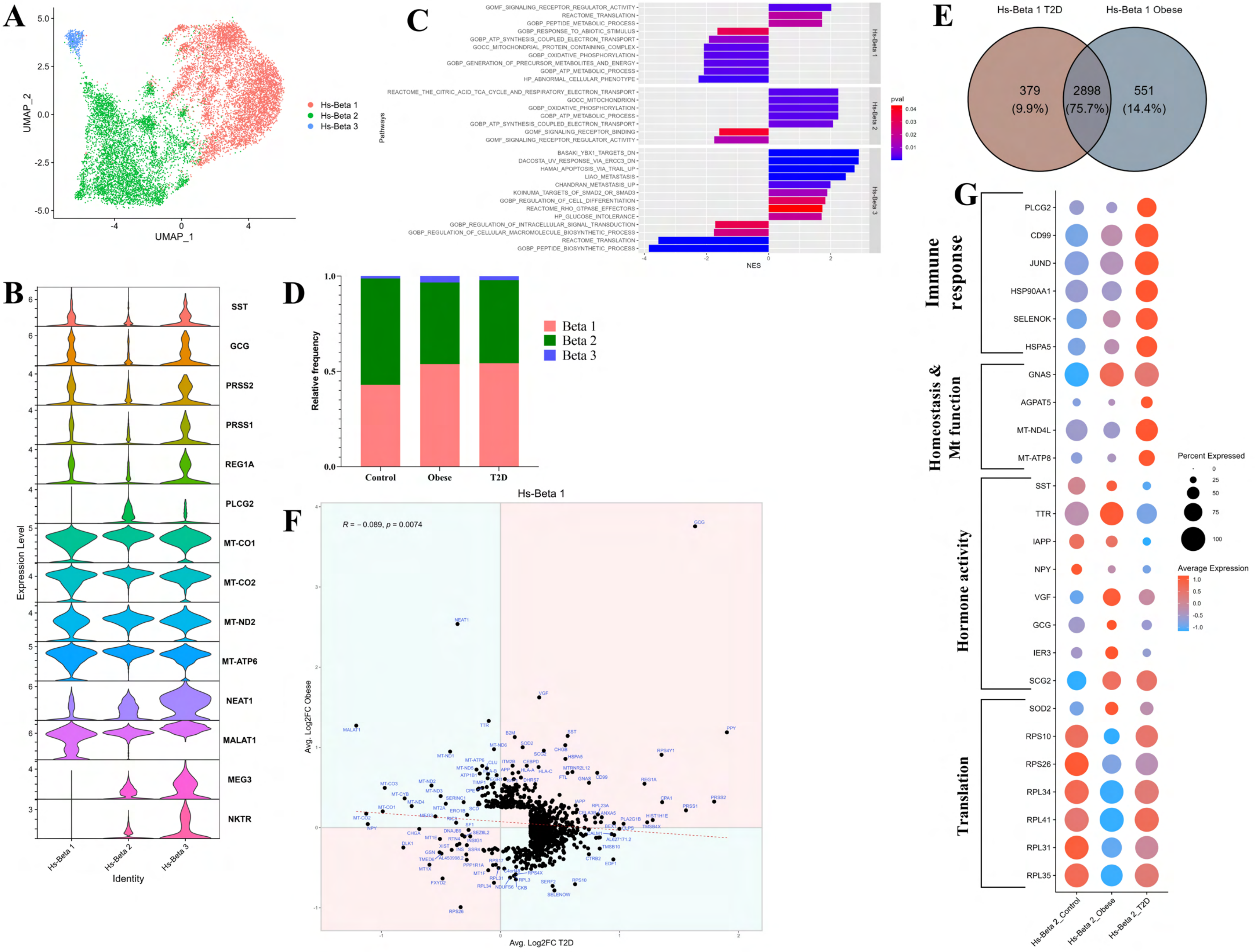

A
UMAP_1
UMAP_2
Hs-Beta 1
Hs-Beta 2
Hs-Beta 3
B
SST
GCG
PRSS2
PRSS1
REG1A
PLCG2
MT-CO1
MT-CO2
MT-ND2
MT-ATP6
NEAT1
MALAT1
MEG3
NKTR
Expression Level
Identity
Hs-Beta 1
Hs-Beta 2
Hs-Beta 3
C
GOMF_SIGNALING_RECEPTOR_REGULATOR_ACTIVITY
REACTOME_TRANSLATION
GOBP_PEPTIDE_METABOLIC_PROCESS
GOBP_RESPONSE_TO_ABIOTIC_STIMULUS
GOBP_ATP_SYNTHESIS_COUPLED_ELECTRON_TRANSPORT
GOCC_MITOCHONDRIAL_PROTEIN_CONTAINING_COMPLEX
GOBP_OXIDATIVE_PHOSPHORYLATION
GOBP_GENERATION_OF_PRECURSOR_METABOLITES_AND_ENERGY
GOBP_ATP_METABOLIC_PROCESS
HP_ABNORMAL_CELLULAR_PHENOTYPE
REACTOME_THE_CITRIC_ACID_TCA_CYCLE_AND_RESPIRATORY_ELECTRON_TRANSPORT
GOCC_MITOCHONDRION
GOBP_OXIDATIVE_PHOSPHORYLATION
GOBP_ATP_METABOLIC_PROCESS
GOBP_ATP_SYNTHESIS_COUPLED_ELECTRON_TRANSPORT
GOMF_SIGNALING_RECEPTOR_BINDING
GOMF_SIGNALING_RECEPTOR_REGULATOR_ACTIVITY
BASAKI_YBX1_TARGETS_DN
DACOSTA_UV_RESPONSE_VIA_ERCC3_DN
HAMAI_APOPTOSIS_VIA_TRAIL_UP
LIAO_METASTASIS
CHANDRAN_METASTASIS_UP
KOINUMA_TARGETS_OF_SMAD2_OR_SMAD3
GOBP_REGULATION_OF_CELL_DIFFERENTIATION
REACTOME_RHO_GTPASE_EFFECTORS
HP_GLUCOSE_INTOLERANCE
GOBP_REGULATION_OF_INTRACELLULAR_SIGNAL_TRANSDUCTION
GOBP_REGULATION_OF_CELLULAR_MACROMOLECULE_BIOSYNTHETIC_PROCESS
REACTOME_TRANSLATION
GOBP_PEPTIDE_BIOSYNTHETIC_PROCESS
Pathways
NES
Hs-Beta 1
Hs-Beta 2
Hs-Beta 3
pval
D
Relative frequency
Control
Obese
T2D
Beta 1
Beta 2
Beta 3
E
Hs-Beta 1 T2D
Hs-Beta 1 Obese
379
(9.9%)
2898
(75.7%)
551
(14.4%)
F
Hs-Beta 1
R = − 0.089, p = 0.0074
Avg. Log2FC Obese
Avg. Log2FC T2D
G
Immune response
Homeostasis & Mt function
Hormone activity
Translation
PLCG2
CD99
JUND
HSP90AA1
SELENOK
HSPA5
GNAS
AGPAT5
MT-ND4L
MT-ATP8
SST
TTR
IAPP
NPY
VGF
GCG
IER3
SCG2
SOD2
RPS10
RPS26
RPL34
RPL41
RPL31
RPL35
Percent Expressed
Average Expression
Hs-Beta 2_Control
Hs-Beta 2_Obese
Hs-Beta 2_T2D

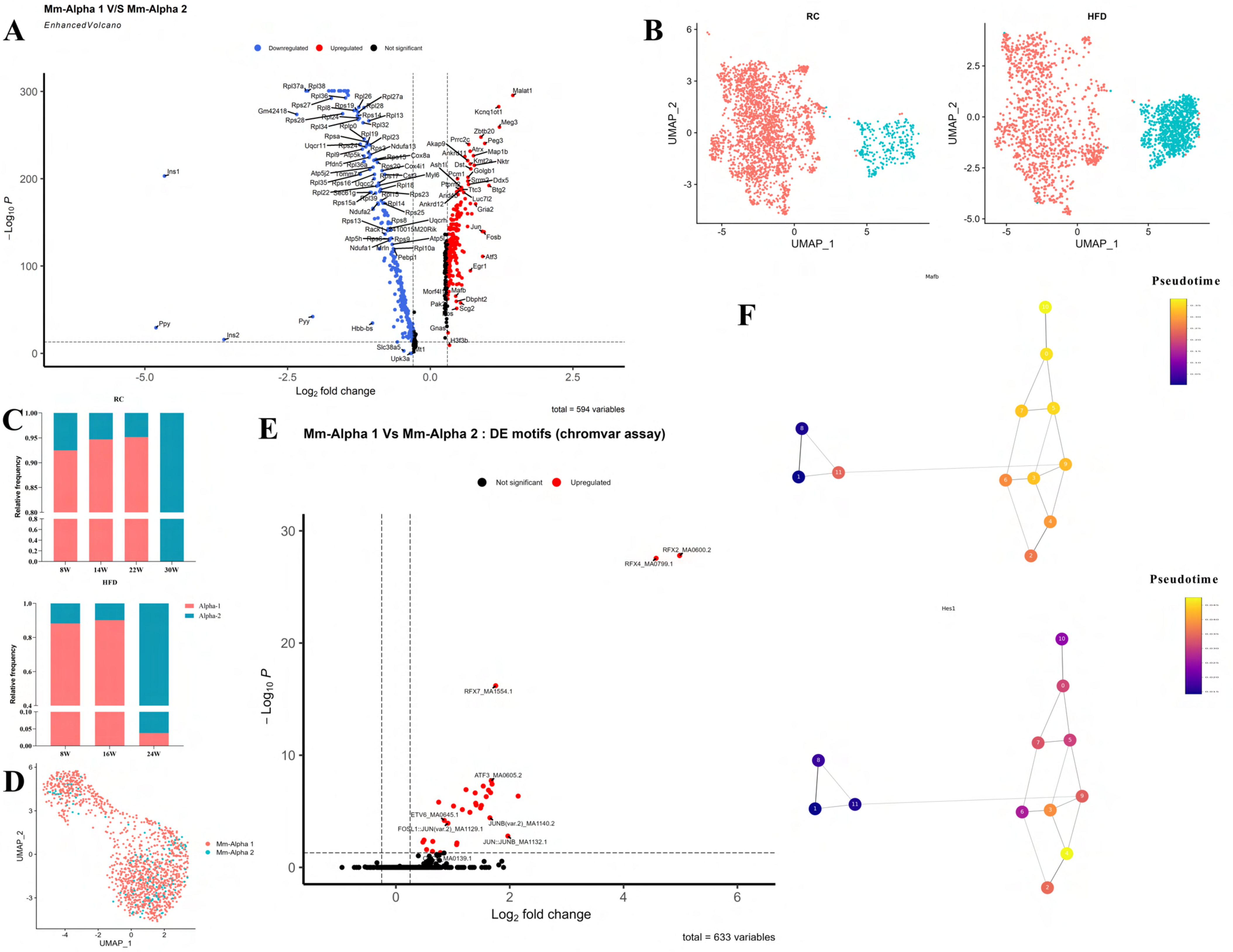
A
Mm-Alpha 1 V/S Mm-Alpha 2
EnhancedVolcano
Downregulated
Upregulated
Not significant
Malat1
Kcnq1ot1
Meg3
Zbtb20
Peg3
Map1b
Nktr
Ddx5
Btg2
Luc7l2
Gria2
Jun
Fosb
Atf3
Egr1
Mafb
Dbpht2
Scg2
H3f3b
Gnas
Ins1
Ppy
Ins2
Pyy
Hbb-bs
Slc38a5
Upk3a
Rpl37a
Rpl38
Rpl36
Rpl26
Rpl27a
Rps27
Rps19
Rpl28
Gm42418
Rps28
Rpl24
Rps14
Rpl13
Rpl32
Rpl34
Rplp0
Rpl19
Rpl23
Rpsa
Uqcr11
Ndufa13
Cox8a
Rps15
Cox4i1
Myl6
Rpl18
Rps23
Rps25
Rps8
Uqcrh
Rps9
Rpl10a
Pebp1
Ndufa1
Mrln
Rps13
Rack1
2410015M20Rik
Atp5h
Rps6
Rps16
Rpl22
Sec61g
Rpl39
Rpl14
Ndufa2
Rps15a
Rpl35
Rpl36a
Pfdn5
Atp5k
Rpl9
Rps24
Rps3
Rpl15
Tomm7
Uqcc2
Rps17
Cst3
Atp5j2
Rpl8
Prrc2c
Akap9
Ankrd11
Atrx
Kmt2a
Golgb1
Dst
Ash1l
Pcm1
Srrm2
Ptprn
Ttc3
Arid4b
Ankrd12
Morf4l1
Pak3
Fos
Mt1
Atp5l
−Log₁₀ P
Log₂ fold change
total = 594 variables
B
RC
HFD
UMAP_1
UMAP_2
C
RC
Relative frequency
8W
14W
22W
30W
HFD
Alpha-1
Alpha-2
8W
16W
24W
D
Mm-Alpha 1
Mm-Alpha 2
UMAP_1
UMAP_2
E
Mm-Alpha 1 Vs Mm-Alpha 2 : DE motifs (chromvar assay)
Not significant
Upregulated
RFX2_MA0600.2
RFX4_MA0799.1
RFX7_MA1554.1
ATF3_MA0605.2
ETV6_MA0645.1
FOSL1::JUN(var.2)_MA1129.1
JUNB(var.2)_MA1140.2
JUN::JUNB_MA1132.1
MA0139.1
− Log₁₀ P
Log₂ fold change
total = 633 variables
F
Mafb
Pseudotime
Hes1
Pseudotime

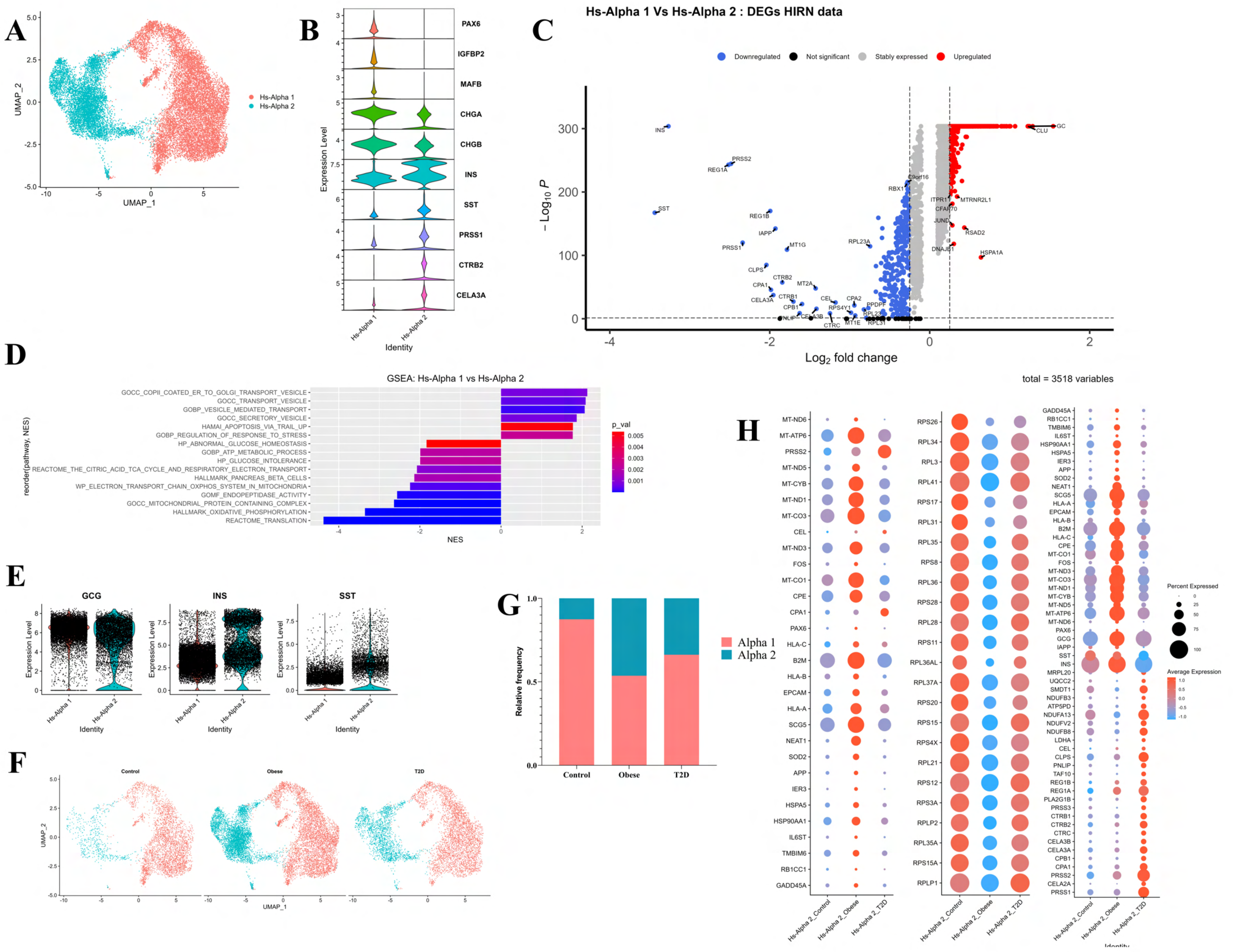

A
UMAP_1
UMAP_2
Hs-Alpha 1
Hs-Alpha 2
B
PAX6
IGFBP2
MAFB
CHGA
CHGB
INS
SST
PRSS1
CTRB2
CELA3A
Expression Level
Identity
C
Hs-Alpha 1 Vs Hs-Alpha 2 : DEGs HIRN data
Downregulated
Not significant
Stably expressed
Upregulated
− Log10 P
Log2 fold change
INS
PRSS2
REG1A
SST
REG1B
IAPP
PRSS1
MT1G
CLPS
CTRB2
CPA1
MT2A
CELA3A
CTRB1
CPB1
CEL
CPA2
RPS4Y1
PPDPF
RPL23A
C9orf16
RBX1
GC
CLU
ITPR1
MTRNR2L1
CFAP70
JUND
RSAD2
DNAJB1
HSPA1A
D
GSEA: Hs-Alpha 1 vs Hs-Alpha 2
reorder(pathway, NES)
GOCC_COPII_COATED_ER_TO_GOLGI_TRANSPORT_VESICLE
GOCC_TRANSPORT_VESICLE
GOBP_VESICLE_MEDIATED_TRANSPORT
GOCC_SECRETORY_VESICLE
HAMAI_APOPTOSIS_VIA_TRAIL_UP
GOBP_REGULATION_OF_RESPONSE_TO_STRESS
HP_ABNORMAL_GLUCOSE_HOMEOSTASIS
GOBP_ATP_METABOLIC_PROCESS
HP_GLUCOSE_INTOLERANCE
REACTOME_THE_CITRIC_ACID_TCA_CYCLE_AND_RESPIRATORY_ELECTRON_TRANSPORT
HALLMARK_PANCREAS_BETA_CELLS
WP_ELECTRON_TRANSPORT_CHAIN_OXPHOS_SYSTEM_IN_MITOCHONDRIA
GOMF_ENDOPEPTIDASE_ACTIVITY
GOCC_MITOCHONDRIAL_PROTEIN_CONTAINING_COMPLEX
HALLMARK_OXIDATIVE_PHOSPHORYLATION
REACTOME_TRANSLATION
NES
p_val
E
GCG
INS
SST
Expression Level
Identity
F
Control
Obese
T2D
UMAP_1
UMAP_2
G
Relative frequency
Control
Obese
T2D
Alpha 1
Alpha 2
H
total = 3518 variables
Hs-Alpha 2_Control
Hs-Alpha 2_Obese
Hs-Alpha 2_T2D
Identity
Percent Expressed
Average Expression

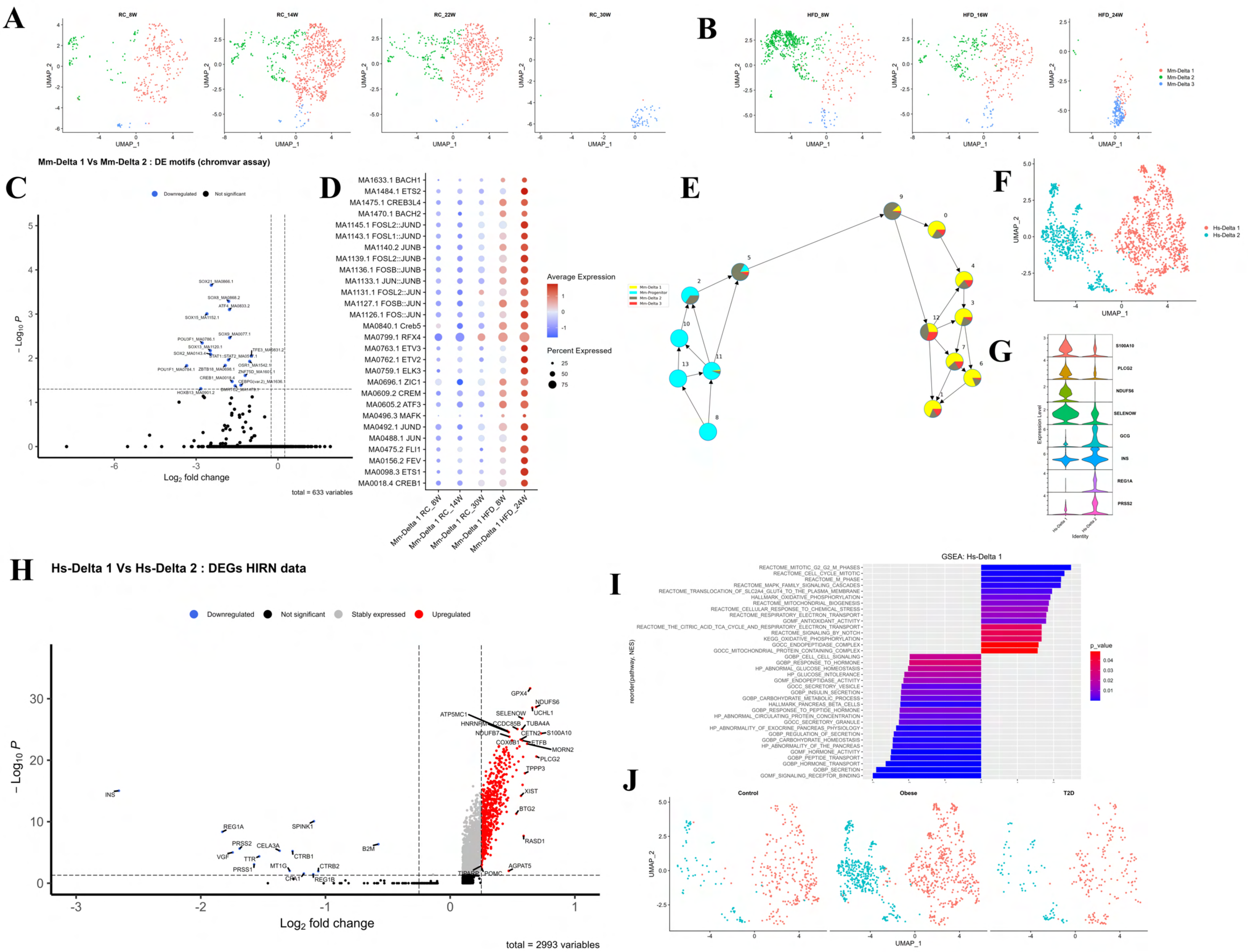
A
RC_8W
RC_14W
RC_22W
RC_30W
B
HFD_8W
HFD_16W
HFD_24W
Mm-Delta 1
Mm-Delta 2
Mm-Delta 3
UMAP_1
UMAP_2
Mm-Delta 1 Vs Mm-Delta 2 : DE motifs (chromvar assay)
C
Downregulated
Not significant
Log2 fold change
total = 633 variables
D
MA1633.1 BACH1
MA1484.1 ETS2
MA1475.1 CREB3L4
MA1470.1 BACH2
MA1145.1 FOSL2::JUND
MA1143.1 FOSL1::JUND
MA1140.2 JUNB
MA1139.1 FOSL2::JUNB
MA1136.1 FOSB::JUNB
MA1133.1 JUN::JUNB
MA1131.1 FOSL2::JUN
MA1127.1 FOSB::JUN
MA1126.1 FOS::JUN
MA0840.1 Creb5
MA0799.1 RFX4
MA0763.1 ETV3
MA0762.1 ETV2
MA0759.1 ELK3
MA0696.1 ZIC1
MA0609.2 CREM
MA0605.2 ATF3
MA0496.3 MAFK
MA0492.1 JUND
MA0488.1 JUN
MA0475.2 FLI1
MA0156.2 FEV
MA0098.3 ETS1
MA0018.4 CREB1
Mm-Delta 1 RC_8W
Mm-Delta 1 RC_14W
Mm-Delta 1 RC_30W
Mm-Delta 1 HFD_8W
Mm-Delta 1 HFD_24W
Average Expression
Percent Expressed
E
Mm-Delta 1
Mm-Progenitor
Mm-Delta 2
Mm-Delta 3
F
Hs-Delta 1
Hs-Delta 2
G
S100A10
PLCG2
NDUFS6
SELENOW
GCG
INS
REG1A
PRSS2
Expression Level
Identity
H
Hs-Delta 1 Vs Hs-Delta 2 : DEGs HIRN data
Downregulated
Not significant
Stably expressed
Upregulated
Log2 fold change
total = 2993 variables
I
GSEA: Hs-Delta 1
REACTOME_MITOTIC_G2_G2_M_PHASES
REACTOME_CELL_CYCLE_MITOTIC
REACTOME_M_PHASE
REACTOME_MAPK_FAMILY_SIGNALING_CASCADES
REACTOME_TRANSLOCATION_OF_SLC2A4_GLUT4_TO_THE_PLASMA_MEMBRANE
HALLMARK_OXIDATIVE_PHOSPHORYLATION
REACTOME_MITOCHONDRIAL_BIOGENESIS
REACTOME_CELLULAR_RESPONSE_TO_CHEMICAL_STRESS
REACTOME_RESPIRATORY_ELECTRON_TRANSPORT
GOMF_ANTIOXIDANT_ACTIVITY
REACTOME_THE_CITRIC_ACID_TCA_CYCLE_AND_RESPIRATORY_ELECTRON_TRANSPORT
REACTOME_SIGNALING_BY_NOTCH
KEGG_OXIDATIVE_PHOSPHORYLATION
GOCC_ENDOPEPTIDASE_COMPLEX
GOCC_MITOCHONDRIAL_PROTEIN_CONTAINING_COMPLEX
GOBP_CELL_CELL_SIGNALING
GOBP_RESPONSE_TO_HORMONE
HP_ABNORMAL_GLUCOSE_HOMEOSTASIS
HP_GLUCOSE_INTOLERANCE
GOMF_ENDOPEPTIDASE_ACTIVITY
GOCC_SECRETORY_VESICLE
GOBP_INSULIN_SECRETION
GOBP_CARBOHYDRATE_METABOLIC_PROCESS
HALLMARK_PANCREAS_BETA_CELLS
GOBP_RESPONSE_TO_PEPTIDE_HORMONE
HP_ABNORMAL_CIRCULATING_PROTEIN_CONCENTRATION
GOCC_SECRETORY_GRANULE
HP_ABNORMALITY_OF_EXOCRINE_PANCREAS_PHYSIOLOGY
GOBP_REGULATION_OF_SECRETION
GOBP_CARBOHYDRATE_HOMEOSTASIS
HP_ABNORMALITY_OF_THE_PANCREAS
GOMF_HORMONE_ACTIVITY
GOBP_PEPTIDE_TRANSPORT
GOBP_HORMONE_TRANSPORT
GOBP_SECRETION
GOMF_SIGNALING_RECEPTOR_BINDING
reorder(pathway, NES)
p_value
J
Control
Obese
T2D
UMAP_1
UMAP_2

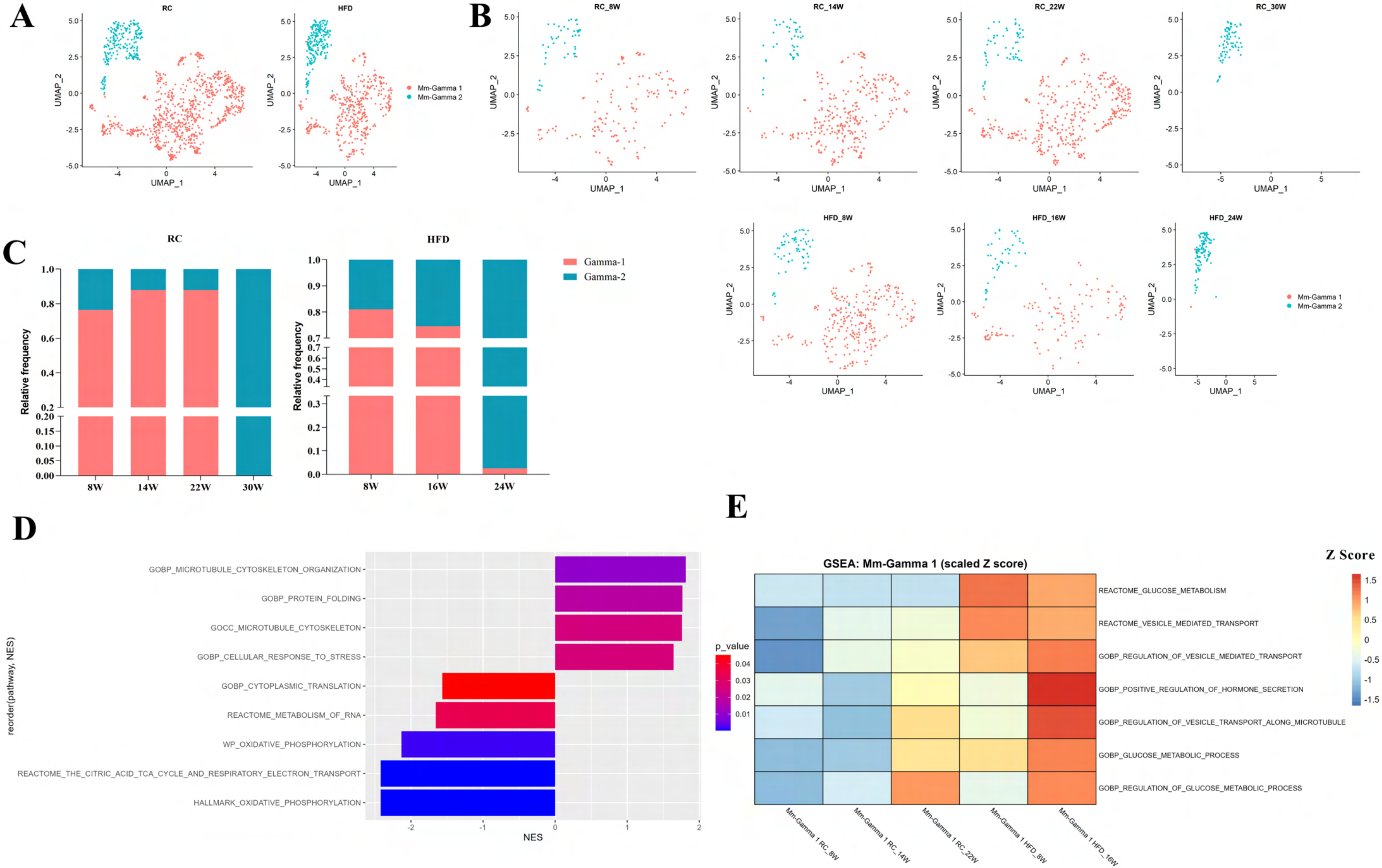
A
RC
HFD
UMAP_1
UMAP_2
Mm-Gamma 1
Mm-Gamma 2
B
RC_8W
RC_14W
RC_22W
RC_30W
HFD_8W
HFD_16W
HFD_24W
C
RC
HFD
Relative frequency
Gamma-1
Gamma-2
8W
14W
22W
30W
16W
24W
D
GOBP_MICROTUBULE_CYTOSKELETON_ORGANIZATION
GOBP_PROTEIN_FOLDING
GOCC_MICROTUBULE_CYTOSKELETON
GOBP_CELLULAR_RESPONSE_TO_STRESS
GOBP_CYTOPLASMIC_TRANSLATION
REACTOME_METABOLISM_OF_RNA
WP_OXIDATIVE_PHOSPHORYLATION
REACTOME_THE_CITRIC_ACID_TCA_CYCLE_AND_RESPIRATORY_ELECTRON_TRANSPORT
HALLMARK_OXIDATIVE_PHOSPHORYLATION
reorder(pathway, NES)
NES
p_value
E
GSEA: Mm-Gamma 1 (scaled Z score)
Z Score
REACTOME_GLUCOSE_METABOLISM
REACTOME_VESICLE_MEDIATED_TRANSPORT
GOBP_REGULATION_OF_VESICLE_MEDIATED_TRANSPORT
GOBP_POSITIVE_REGULATION_OF_HORMONE_SECRETION
GOBP_REGULATION_OF_VESICLE_TRANSPORT_ALONG_MICROTUBULE
GOBP_GLUCOSE_METABOLIC_PROCESS
GOBP_REGULATION_OF_GLUCOSE_METABOLIC_PROCESS
Mm-Gamma 1 RC_8W
Mm-Gamma 1 RC_14W
Mm-Gamma 1 RC_22W
Mm-Gamma 1 HFD_8W
Mm-Gamma 1 HFD_16W

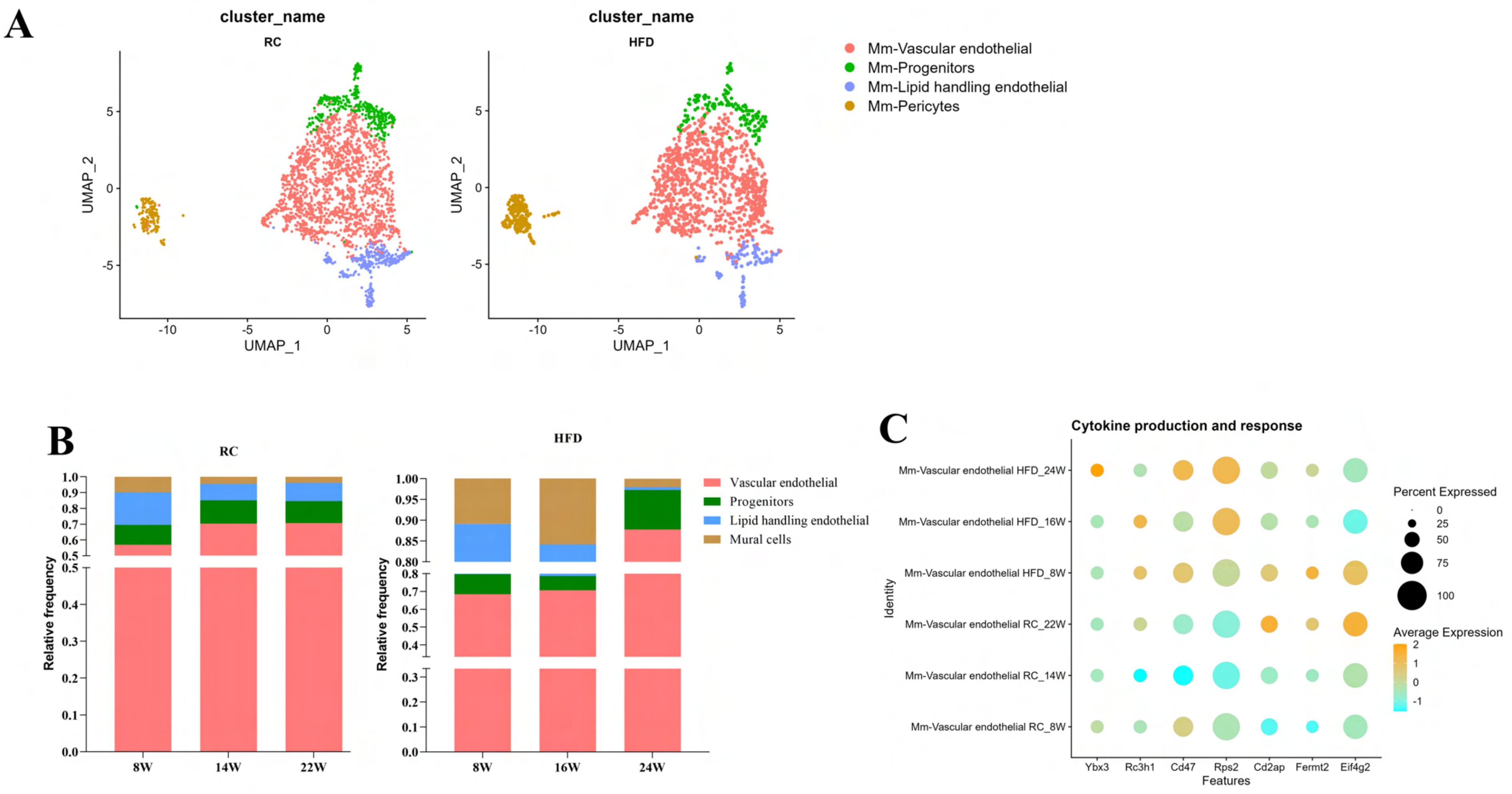
A
cluster_name
RC
HFD
UMAP_1
UMAP_2
Mm-Vascular endothelial
Mm-Progenitors
Mm-Lipid handling endothelial
Mm-Pericytes
B
RC
HFD
Relative frequency
8W
14W
22W
16W
24W
Vascular endothelial
Progenitors
Lipid handling endothelial
Mural cells
C
Cytokine production and response
Identity
Mm-Vascular endothelial HFD_24W
Mm-Vascular endothelial HFD_16W
Mm-Vascular endothelial HFD_8W
Mm-Vascular endothelial RC_22W
Mm-Vascular endothelial RC_14W
Mm-Vascular endothelial RC_8W
Ybx3
Rc3h1
Cd47
Rps2
Cd2ap
Fermt2
Eif4g2
Features
Percent Expressed
Average Expression

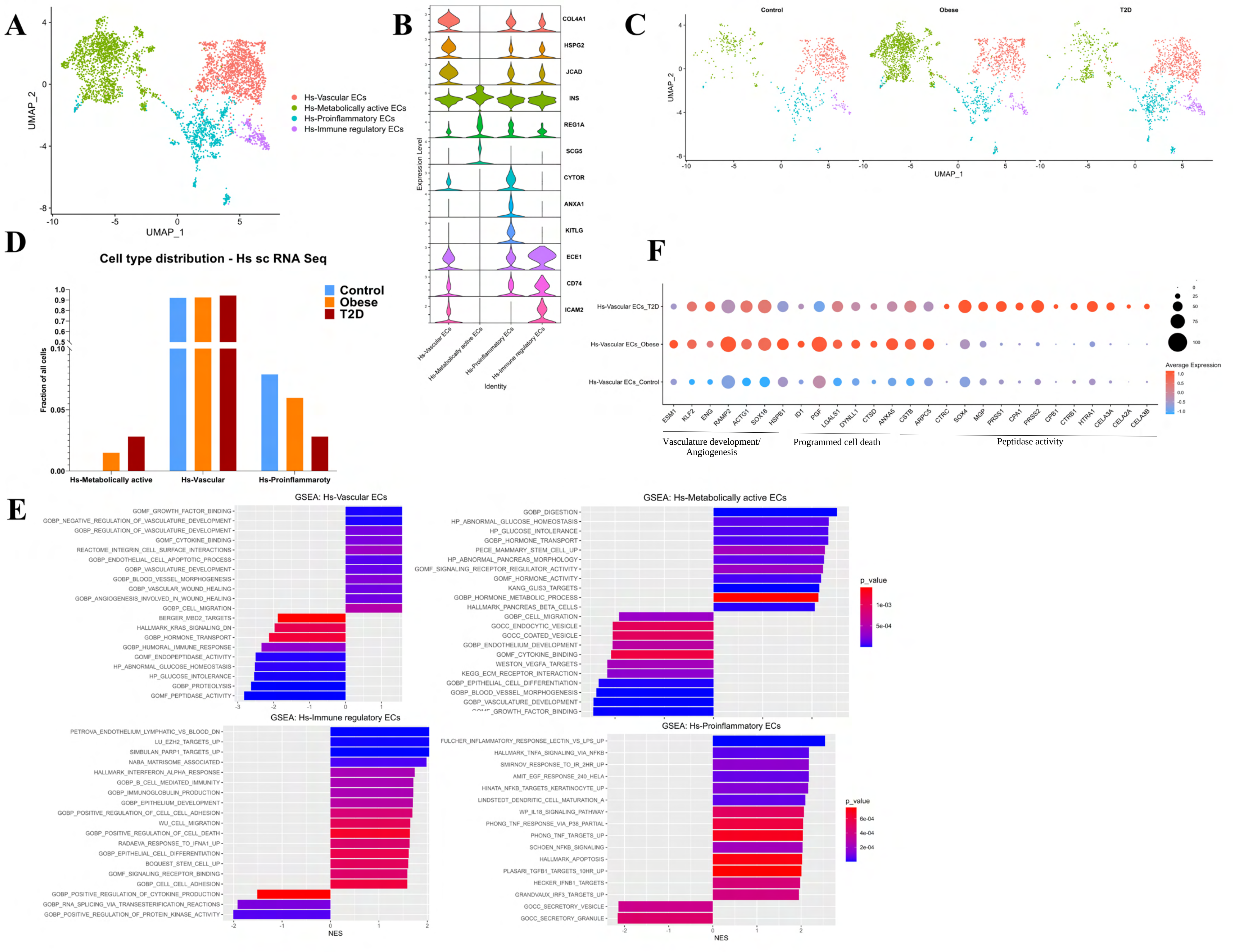

A
Hs-Vascular ECs
Hs-Metabolically active ECs
Hs-Proinflammatory ECs
Hs-Immune regulatory ECs
UMAP_1
UMAP_2
B
COL4A1
HSPG2
JCAD
INS
REG1A
SCG5
CYTOR
ANXA1
KITLG
ECE1
CD74
ICAM2
Expression Level
Identity
C
Control
Obese
T2D
D
Cell type distribution - Hs sc RNA Seq
Control
Obese
T2D
Fraction of all cells
Hs-Metabolically active
Hs-Vascular
Hs-Proinflammaroty
F
Hs-Vascular ECs_T2D
Hs-Vascular ECs_Obese
Hs-Vascular ECs_Control
Average Expression
ESM1
KLF2
ENG
RAMP2
ACTG1
SOX18
HSPB1
ID1
PGF
LGALS1
DYNLL1
CTSD
ANXA5
CSTB
ARPC5
CTRC
SOX4
MGP
PRSS1
CPA1
PRSS2
CPB1
CTRB1
HTRA1
CELA3A
CELA2A
CELA3B
Vasculature development/ Angiogenesis
Programmed cell death
Peptidase activity
E
GSEA: Hs-Vascular ECs
GOMF_GROWTH_FACTOR_BINDING
GOBP_NEGATIVE_REGULATION_OF_VASCULATURE_DEVELOPMENT
GOBP_REGULATION_OF_VASCULATURE_DEVELOPMENT
GOMF_CYTOKINE_BINDING
REACTOME_INTEGRIN_CELL_SURFACE_INTERACTIONS
GOBP_ENDOTHELIAL_CELL_APOPTOTIC_PROCESS
GOBP_VASCULATURE_DEVELOPMENT
GOBP_BLOOD_VESSEL_MORPHOGENESIS
GOBP_VASCULAR_WOUND_HEALING
GOBP_ANGIOGENESIS_INVOLVED_IN_WOUND_HEALING
GOBP_CELL_MIGRATION
BERGER_MBD2_TARGETS
HALLMARK_KRAS_SIGNALING_DN
GOBP_HORMONE_TRANSPORT
GOBP_HUMORAL_IMMUNE_RESPONSE
GOMF_ENDOPEPTIDASE_ACTIVITY
HP_ABNORMAL_GLUCOSE_HOMEOSTASIS
HP_GLUCOSE_INTOLERANCE
GOBP_PROTEOLYSIS
GOMF_PEPTIDASE_ACTIVITY
GSEA: Hs-Metabolically active ECs
GOBP_DIGESTION
HP_ABNORMAL_GLUCOSE_HOMEOSTASIS
HP_GLUCOSE_INTOLERANCE
GOBP_HORMONE_TRANSPORT
PECE_MAMMARY_STEM_CELL_UP
HP_ABNORMAL_PANCREAS_MORPHOLOGY
GOMF_SIGNALING_RECEPTOR_REGULATOR_ACTIVITY
GOMF_HORMONE_ACTIVITY
KANG_GLIS3_TARGETS
GOBP_HORMONE_METABOLIC_PROCESS
HALLMARK_PANCREAS_BETA_CELLS
GOBP_CELL_MIGRATION
GOCC_ENDOCYTIC_VESICLE
GOCC_COATED_VESICLE
GOBP_ENDOTHELIUM_DEVELOPMENT
GOMF_CYTOKINE_BINDING
WESTON_VEGFA_TARGETS
KEGG_ECM_RECEPTOR_INTERACTION
GOBP_EPITHELIAL_CELL_DIFFERENTIATION
GOBP_BLOOD_VESSEL_MORPHOGENESIS
GOBP_VASCULATURE_DEVELOPMENT
_GROWTH_FACTOR_BINDING
p_value
GSEA: Hs-Immune regulatory ECs
PETROVA_ENDOTHELIUM_LYMPHATIC_VS_BLOOD_DN
LU_EZH2_TARGETS_UP
SIMBULAN_PARP1_TARGETS_UP
NABA_MATRISOME_ASSOCIATED
HALLMARK_INTERFERON_ALPHA_RESPONSE
GOBP_B_CELL_MEDIATED_IMMUNITY
GOBP_IMMUNOGLOBULIN_PRODUCTION
GOBP_EPITHELIUM_DEVELOPMENT
GOBP_POSITIVE_REGULATION_OF_CELL_CELL_ADHESION
WU_CELL_MIGRATION
GOBP_POSITIVE_REGULATION_OF_CELL_DEATH
RADAEVA_RESPONSE_TO_IFNA1_UP
GOBP_EPITHELIAL_CELL_DIFFERENTIATION
BOQUEST_STEM_CELL_UP
GOMF_SIGNALING_RECEPTOR_BINDING
GOBP_CELL_CELL_ADHESION
GOBP_POSITIVE_REGULATION_OF_CYTOKINE_PRODUCTION
GOBP_RNA_SPLICING_VIA_TRANSESTERIFICATION_REACTIONS
GOBP_POSITIVE_REGULATION_OF_PROTEIN_KINASE_ACTIVITY
NES
GSEA: Hs-Proinflammatory ECs
FULCHER_INFLAMMATORY_RESPONSE_LECTIN_VS_LPS_UP
HALLMARK_TNFA_SIGNALING_VIA_NFKB
SMIRNOV_RESPONSE_TO_IR_2HR_UP
AMIT_EGF_RESPONSE_240_HELA
HINATA_NFKB_TARGETS_KERATINOCYTE_UP
LINDSTEDT_DENDRITIC_CELL_MATURATION_A
WP_IL18_SIGNALING_PATHWAY
PHONG_TNF_RESPONSE_VIA_P38_PARTIAL
PHONG_TNF_TARGETS_UP
SCHOEN_NFKB_SIGNALING
HALLMARK_APOPTOSIS
PLASARI_TGFB1_TARGETS_10HR_UP
HECKER_IFNB1_TARGETS
GRANDVAUX_IRF3_TARGETS_UP
GOCC_SECRETORY_VESICLE
GOCC_SECRETORY_GRANULE
p_value
NES

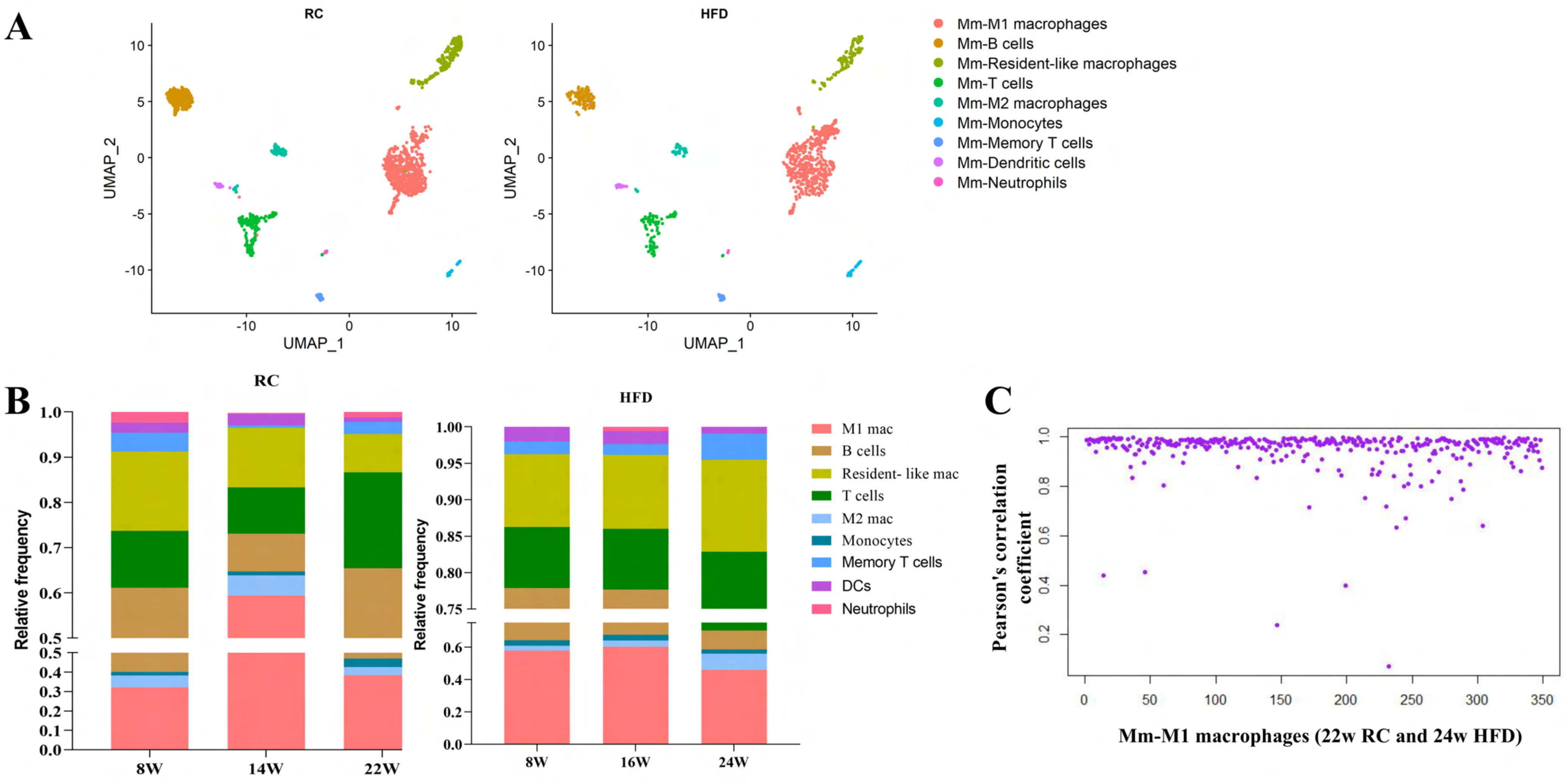
A
RC
HFD
UMAP_1
UMAP_2
Mm-M1 macrophages
Mm-B cells
Mm-Resident-like macrophages
Mm-T cells
Mm-M2 macrophages
Mm-Monocytes
Mm-Memory T cells
Mm-Dendritic cells
Mm-Neutrophils
B
RC
HFD
Relative frequency
8W
14W
22W
16W
24W
M1 mac
B cells
Resident- like mac
T cells
M2 mac
Monocytes
Memory T cells
DCs
Neutrophils
C
Pearson's correlation coefficient
Mm-M1 macrophages (22w RC and 24w HFD)

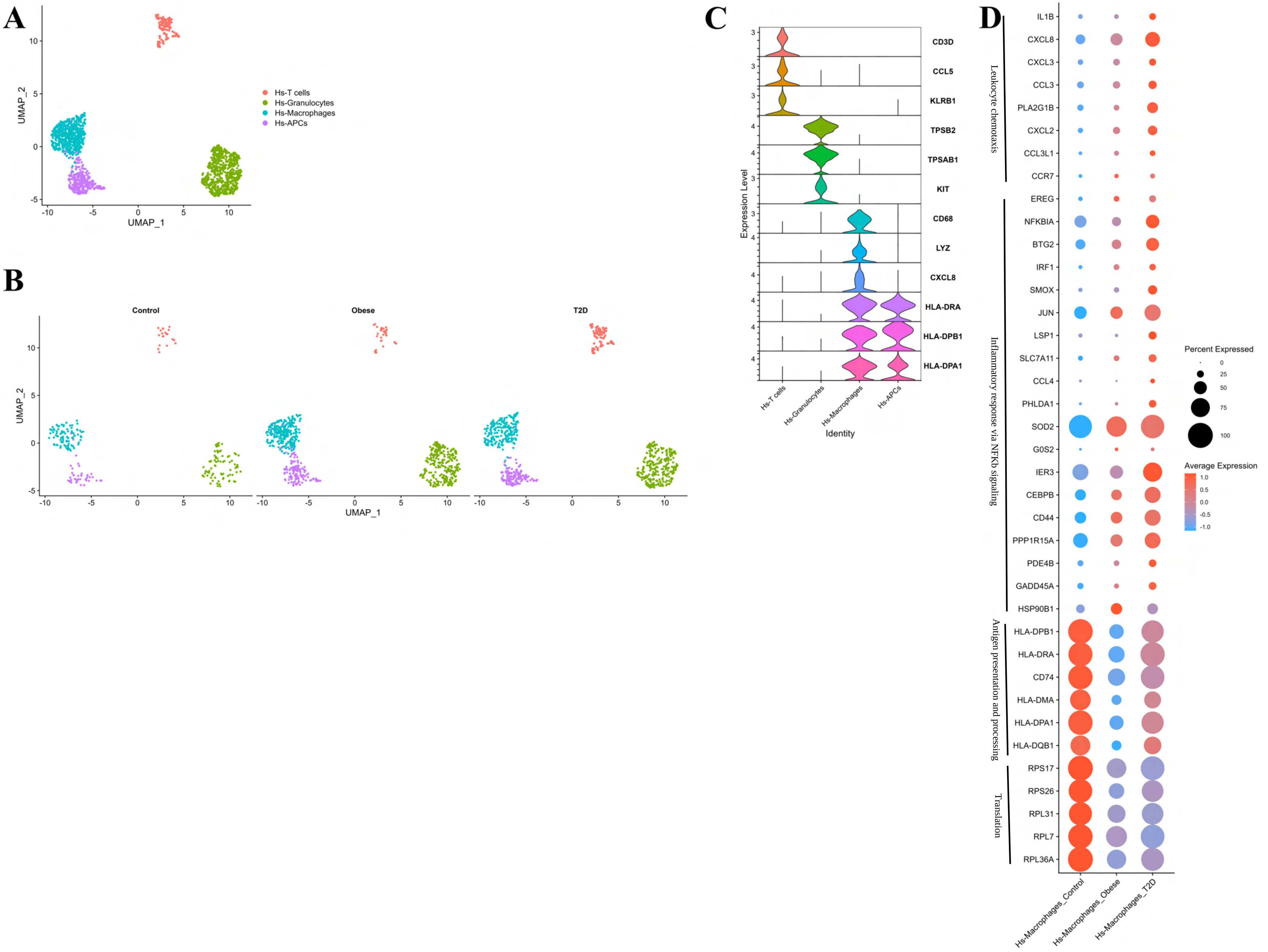

A
B
C
D
UMAP_1
UMAP_2
Hs-T cells
Hs-Granulocytes
Hs-Macrophages
Hs-APCs
Control
Obese
T2D
Expression Level
Identity
CD3D
CCL5
KLRB1
TPSB2
TPSAB1
KIT
CD68
LYZ
CXCL8
HLA-DRA
HLA-DPB1
HLA-DPA1
Leukocyte chemotaxis
Inflammatory response via NFKb signaling
Antigen presentation and processing
Translation
IL1B
CXCL8
CXCL3
CCL3
PLA2G1B
CXCL2
CCL3L1
CCR7
EREG
NFKBIA
BTG2
IRF1
SMOX
JUN
LSP1
SLC7A11
CCL4
PHLDA1
SOD2
G0S2
IER3
CEBPB
CD44
PPP1R15A
PDE4B
GADD45A
HSP90B1
HLA-DPB1
HLA-DRA
CD74
HLA-DMA
HLA-DPA1
HLA-DQB1
RPS17
RPS26
RPL31
RPL7
RPL36A
Percent Expressed
Average Expression
Hs-Macrophages_Control
Hs-Macrophages_Obese
Hs-Macrophages_T2D

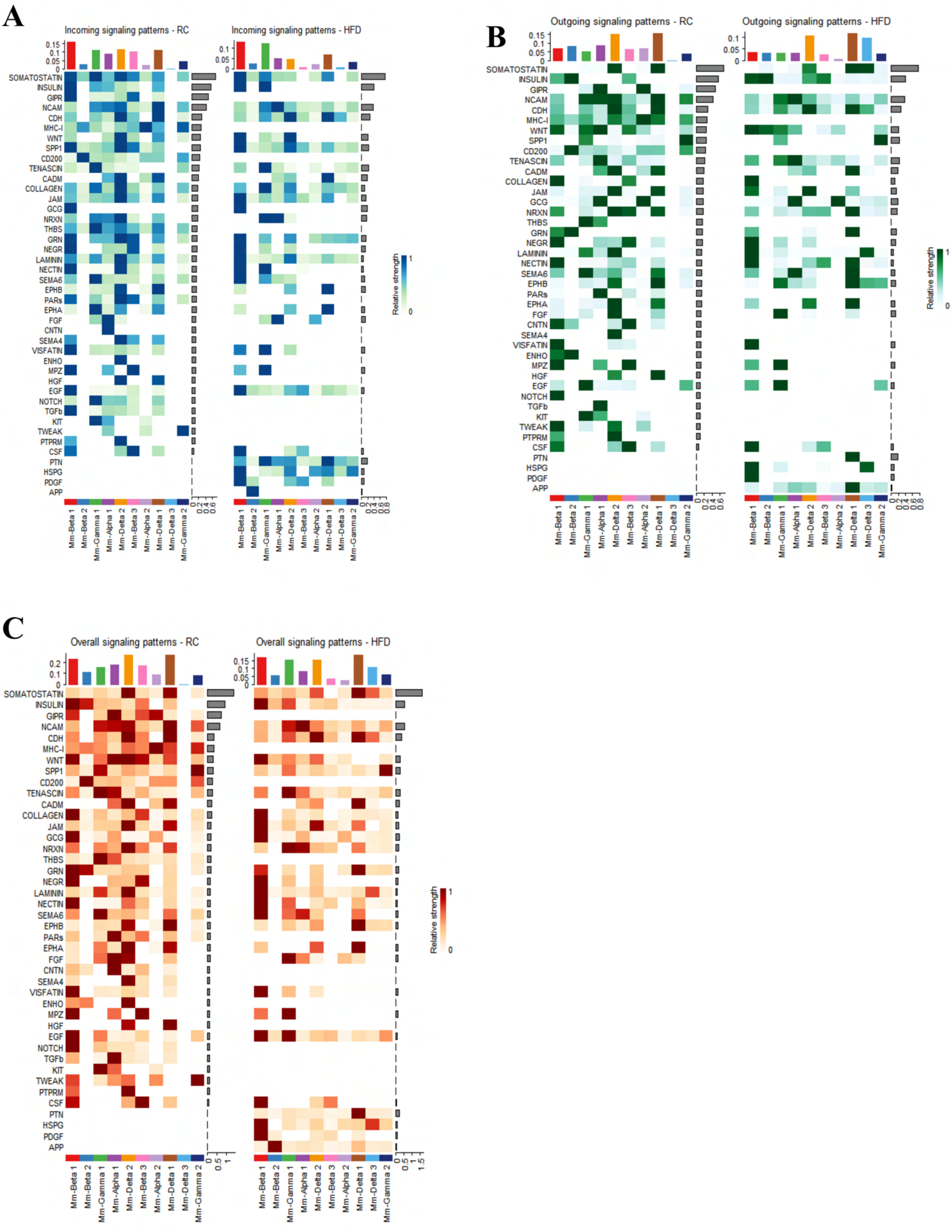

A
Incoming signaling patterns - RC
Incoming signaling patterns - HFD
B
Outgoing signaling patterns - RC
Outgoing signaling patterns - HFD
C
Overall signaling patterns - RC
Overall signaling patterns - HFD
SOMATOSTATIN
INSULIN
GIPR
NCAM
CDH
MHC-I
WNT
SPP1
CD200
TENASCIN
CADM
COLLAGEN
JAM
GCG
NRXN
THBS
GRN
NEGR
LAMININ
NECTIN
SEMA6
EPHB
PARs
EPHA
FGF
CNTN
SEMA4
VISFATIN
ENHO
MPZ
HGF
EGF
NOTCH
TGFb
KIT
TWEAK
PTPRM
CSF
PTN
HSPG
PDGF
APP
Mm-Beta 1
Mm-Beta 2
Mm-Gamma 1
Mm-Alpha 1
Mm-Delta 2
Mm-Beta 3
Mm-Alpha 2
Mm-Delta 1
Mm-Delta 3
Mm-Gamma 2
Relative strength

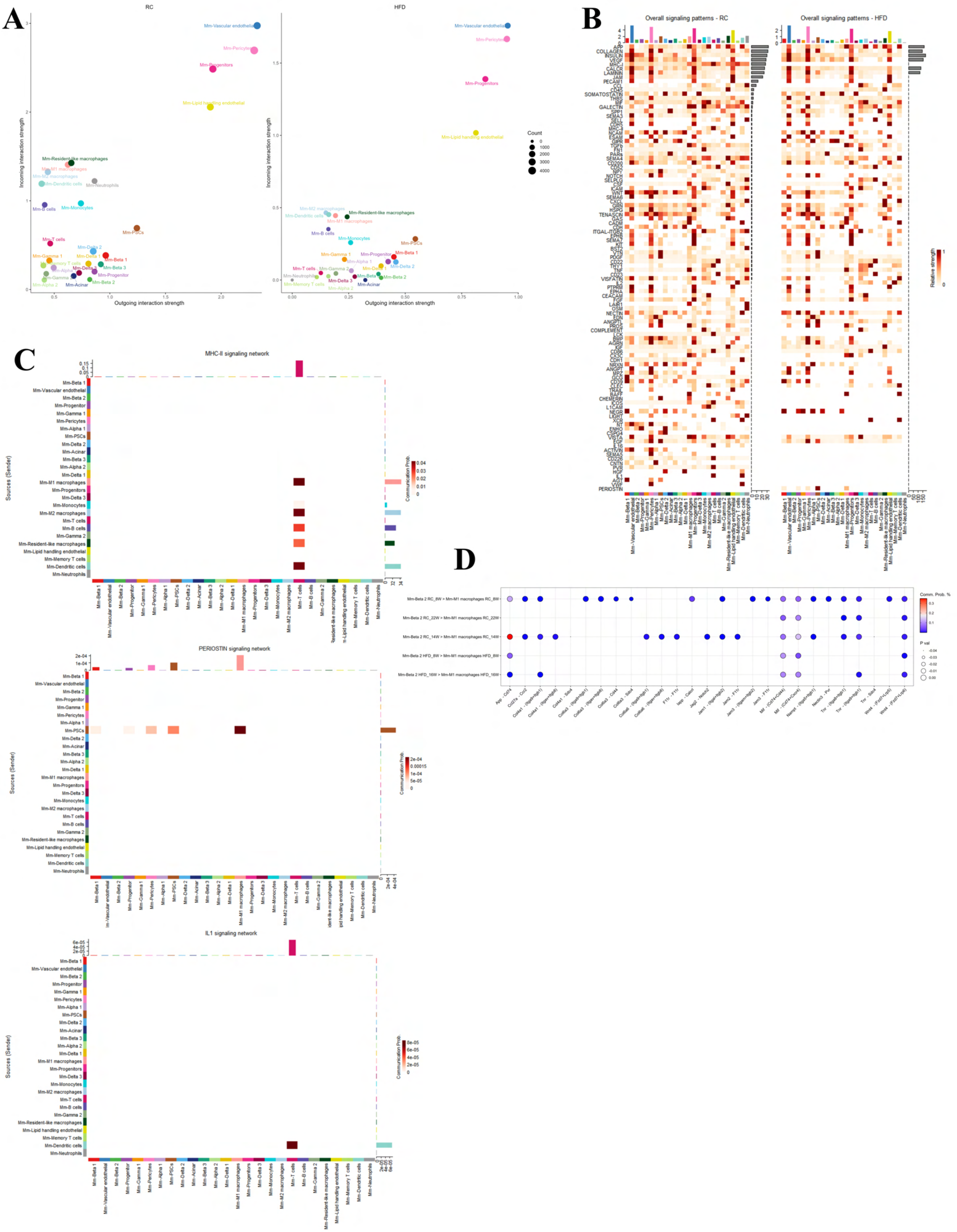
A
RC
HFD
Incoming interaction strength
Outgoing interaction strength
Count
0
1000
2000
3000
4000
Mm-Vascular endothelial
Mm-Pericytes
Mm-Progenitors
Mm-Lipid handling endothelial
Mm-Resident-like macrophages
Mm-M1 macrophages
Mm-M2 macrophages
Mm-Dendritic cells
Mm-Neutrophils
Mm-B cells
Mm-Monocytes
Mm-PSCs
Mm-T cells
Mm-Delta 2
Mm-Gamma 1
Mm-Delta 1
Mm-Beta 1
Mm-Memory T cells
Mm-Alpha 1
Mm-Delta 3
Mm-Beta 3
Mm-Progenitor
Mm-Gamma 2
Mm-Beta 2
Mm-Alpha 2
Mm-Acinar
B
Overall signaling patterns - RC
Overall signaling patterns - HFD
Relative strength
C
MHC-II signaling network
Sources (Sender)
Communication Prob.
PERIOSTIN signaling network
IL1 signaling network
D
Comm. Prob. %
P val
Mm-Beta 2 RC_8W > Mm-M1 macrophages RC_8W
Mm-Beta 2 RC_22W > Mm-M1 macrophages RC_22W
Mm-Beta 2 RC_14W > Mm-M1 macrophages RC_14W
Mm-Beta 2 HFD_8W > Mm-M1 macrophages HFD_8W
Mm-Beta 2 HFD_16W > Mm-M1 macrophages HFD_16W

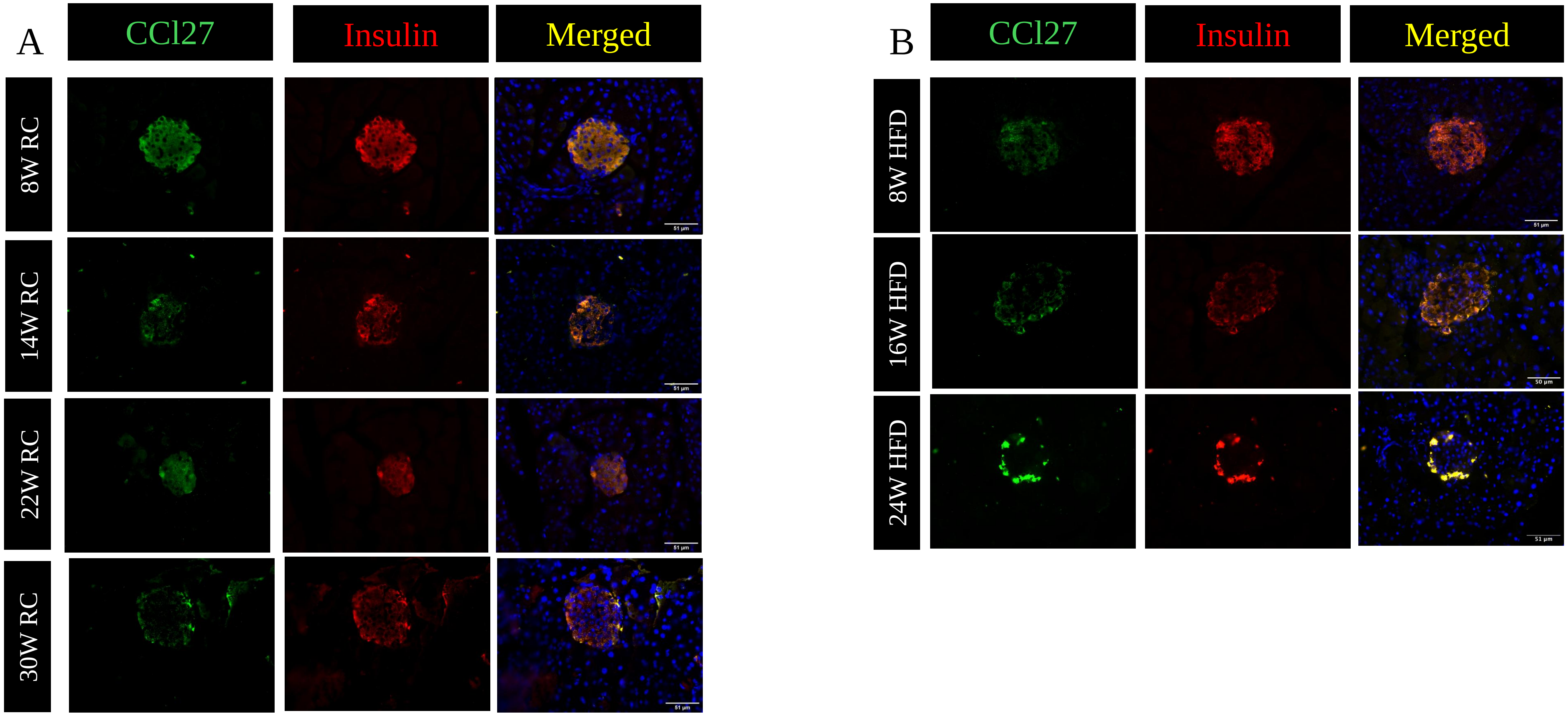
A
CCl27
Insulin
Merged
8W RC
14W RC
22W RC
30W RC
B
CCl27
Insulin
Merged
8W HFD
16W HFD
24W HFD
